# Programmable DNA integration with New-to-Nature tools using Computational Protein Design

**DOI:** 10.64898/2026.05.29.728539

**Authors:** Hailey M. Wallace, Seong Guk Park, Adam Smiley, Tuba Şevik, Shubham Dubey, Shirin Fatma, Samuel Chau-Duy-Tam Vo, Ethan Creed, Alexandre Zanghellini, Elizabeth H. Kellogg

**Affiliations:** Center for Protein Design, Center of Excellence for Data Driven Discovery, St Jude Children’s Research Hospital, Memphis, TN; Department of Structural Biology, St. Jude Children’s Research Hospital, Memphis, TN; Arzeda Corporation, Seattle, WA

## Abstract

Programmable integration of large DNA cargo (≥ 2 kb), without inducing double-strand breaks, remains challenging for genome editing technologies. Current approaches have limitations in programmability, depend on co-delivery of multiple components, require multiple enzymatic steps, or have variable on-target editing outcomes. Here, we address this challenge using *de novo* protein design to create highly active, new-to-nature RNA-guided transposons. Our strategy exploits the modular architecture of CRISPR-associated transposons (CASTs), reconfiguring their conserved transposition machinery to interface with widely adopted Cas9. The resulting system, which we call NovoCAST, simplifies the CAST architecture from eight distinct proteins to four, establishing the simplest CAST described to date. NovoCAST exhibits sharply defined integration profiles, a 500-fold increase in activity relative to the parental PmcCAST, and general programmability. Using structural and biochemical analyses, we confirmed that the designed proteins fold and function as intended. Finally, we demonstrate robust programmable genomic integration in human cells highlighting its broad potential applications in research and therapeutics. Together, these results establish *de novo* protein design as a powerful strategy for engineering efficient genome-editing systems and for coupling CRISPR-mediated DNA recognition to heterologous functions through *de novo* designed protein interfaces.

## Introduction

Over the past decade, genome editing has advanced from concept to clinical reality, enabling transformative therapies for genetic diseases such as sickle cell disease (Frangoul et al. 2021). Base editing is well suited for single-base corrections (Gaudelli et al. 2017), whereas prime editing enables small insertions and deletions of less than 100 base pairs (bp) (Anzalone et al. 2019). On the other hand, precise integration of large DNA cargo is required for many therapeutic applications, such as restoration of gene function using ‘superexon’ strategies and targeted insertion of genetic cassettes into endogenous loci for cancer immunotherapies (Eyquem et al. 2017). Current approaches for the integration of large DNA cargo remain limited. Nuclease-mediated homologous recombination enables targeted insertion of large DNA payloads (Mali et al. 2013; Cong et al. 2013), but this process relies on DNA double-strand breaks, which are inefficient and commonly result in cytotoxicity and large chromosomal rearrangements due to non-homologous end-joining (Kosicki et al. 2018). Prime editing can be coupled to site-specific integrases, but this approach requires a complex editing design involving numerous components and remains susceptible to indel byproducts (Yarnall et al. 2023; Pandey et al. 2025). R2 retrotransposons are challenging to reprogram, exhibit limited processivity, and often result in partial integrations (Zhang et al. 2025), whereas bridge recombinases, albeit promising, are constrained by short target site recognition sequences (Durrant et al. 2024), limiting their ability to uniquely target sites across the human genome.

In contrast, CRISPR-associated transposases (CASTs) provide an elegant solution to this central challenge in genome engineering by enabling RNA-guided integration of large DNA cargo without introducing double-strand breaks. Whereas canonical transposases either integrate randomly throughout the genome or exhibit strict specificity for a single location (Craig 2015), recent bioinformatic analyses have identified transposases in bacteria and archaea that have co-opted CRISPR-Cas machinery to achieve RNA-guided integration (Peters et al. 2017). CAST systems therefore provide a general strategy for programmable genomic integration (PGI), enabling targeted integration of large DNA cargo at user-defined genomic loci. Several CASTs have demonstrated strong on-target integration specificity and well-defined integration profiles, achieving up to 99% on-target integration for select guide RNAs (gRNAs) (Strecker et al. 2019; Klompe et al. 2019). CAST targeting is mediated by long gRNAs (20–32 nt) (Saito et al. 2021), which are sufficient to uniquely target sites in the human genome (Hsieh and Peters 2024). Together, these properties make CASTs a promising tool for PGI.

All CASTs encode a conserved set of proteins: a CRISPR effector as the targeting module, a bridging protein TniQ, and Tn7-like transposition machinery consisting of the transposase TnsB, endonuclease TnsA, and AAA+ ATPase TnsC (**Figure 1A**) (Hsieh and Peters 2024). This multi-component architecture often results in low activity in heterologous contexts, including human cells (Liu et al. 2025; Lampe et al. 2024). In addition, CAST activity often depends on bacterial-encoded factors, which are challenging to identify (Schmitz et al. 2022; Walker et al. 2023). Comprehensive surveys of CAST orthologs have yet to reveal a system that is both robust and portable across organisms, motivating efforts to engineer improved variants.

**Figure 1.**
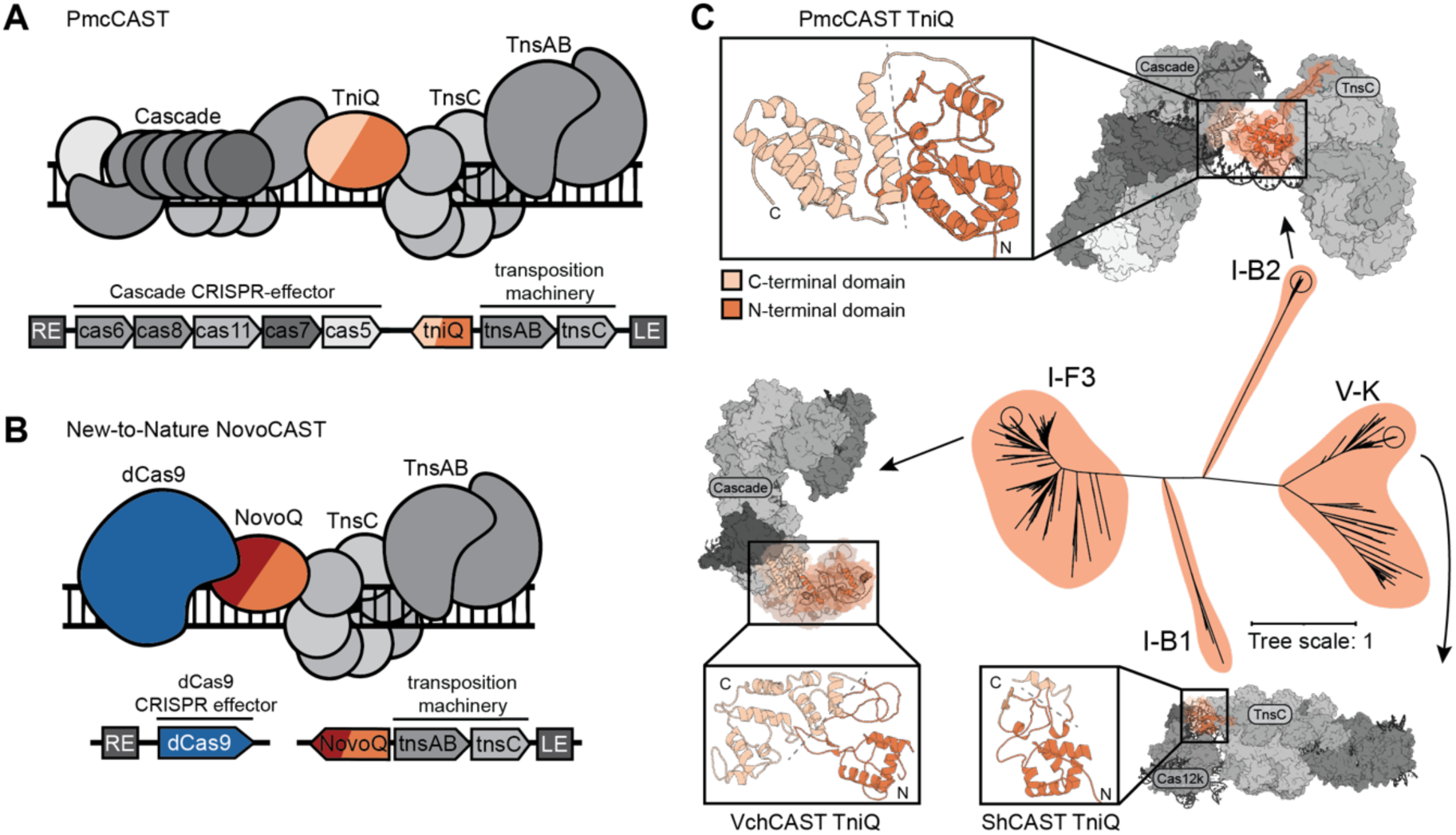
Computational protein design strategy leverages TniQ modularity to design a minimal, new-to-nature CAST. **(A) Architecture of PmcCAST.** Top: Conceptual illustration of the PmcCAST integration complex, including target DNA (black lines), Cascade CRISPR-effector (shades of gray, cas6, cas8, cas11, cas7, and cas5), TniQ (beige/orange), and the transposition machinery (shades of gray, TnsAB and TnsC). Bottom: Diagram showing organization of PmcCAST genes, including transposon right end (RE)/left end (LE) and gene orientations indicated as found in the native operon. TniQ is shaded in beige and orange, corresponding to the C-terminal domain and the N-terminal domain, respectively. **(B) Architecture of the computationally designed NovoCAST.** Proposed architecture of the NovoCAST integration complex (top) and gene organization (bottom), using the same convention as panel A. dCas9, a nuclease-defective SpCas9, is colored blue. NovoQ is a de novo designed protein that retains the conserved N-terminal domain (orange) while incorporating a *de novo* C-terminal domain (red). **(C) Structural and evolutionary diversity of TniQ across CAST subtypes.** Phylogenetic tree of CAST TniQ, branches (shaded orange) are labeled with the CAST subtype (starting from the top clade and proceeding clockwise: Types I-B2, V-K, I-B1, and I-F3). Circles indicate location of representatives whose structures are shown (indicated with arrows): Type I-B2 (PmcCAST), PDB: 8FF4; Type V-K (ShCAST), PDB: 8EA3; Type I-F3 (VchCAST), PDB: 6PIJ. There is currently no structure of a Type I-B1 CAST. N- and C-termini of TniQ are labeled N and C, respectively. The N-terminal domain and C-terminal domain are colored orange and beige, respectively.

Current engineering strategies to improve CASTs as a gene editing tool have yielded limited success. Simple fusions reduce the number of components, but have failed to enhance on-target activity, and the generality of this approach has not been demonstrated (Schargel et al. 2025; Tou et al. 2023). Directed evolution can improve integration activity but does not fundamentally simplify native systems (Witte et al. 2025). These limitations suggest that enhancing CAST activity in heterologous systems will require more extensive engineering of their architectures. Recent work indicates that target-site recognition of CASTs may be limited by their native CRISPR effector (Lampe et al. 2024), suggesting that replacing that component could both enhance CAST integration efficiencies and simplify the multi-component architecture.

Here, we use *de novo* protein design to replace the native CRISPR effector, a multi-component complex called Cascade, of the Type I-B2 CAST (PmcCAST) with a designed variant of nuclease-defective *Streptococcus pyogenes* Cas9 (dCas9) (Jinek et al. 2012). With this strategy, through four rounds of computational design we generate a new-to-nature PGI tool, termed NovoCAST, that enables programmable integration across multiple therapeutically relevant loci in the human genome. Together, these results establish *de novo* protein design as a general strategy for coupling CRISPR effectors to functions beyond DNA cleavage, enabling next-generation genome-editing systems with therapeutic potential.

## Results

### Computational protein design strategies to simplify CAST architecture

In general, our strategy follows a design-build-test-learn framework: designs are generated computationally based on structural and mechanistic hypotheses, encoded as DNA libraries, and evaluated in a quantitative, high-throughput integration screen to inform subsequent design rounds. We selected PmcCAST for its inherent designability; its recruitment complex has been structurally characterized (Wang et al. 2023), and TniQ engages Cascade primarily through protein–protein interactions (**Figure S1A**). This presents a more tractable interface than systems in which TniQ predominantly relies on protein–RNA interactions (**Figure S1B**) or forms a homodimer (**Figure S1C**). Successful redesign of this interface would replace the five-protein Cascade complex with a single protein, dCas9 (**Figure 1B**), dramatically reducing the complexity of the targeting module.

The evolutionary diversity of TniQ suggests that variants can be extensively altered while retaining the capacity to selectively activate transposition machinery. TniQ serves multiple functions: **1.** to physically bridge by associating with the CRISPR effectors (Halpin-Healy et al. 2020), **2.** to recruit transposition machinery to the target-site (J.-U. Park et al. 2023; Schmitz et al. 2022) and **3.** to allosterically initiate integration (George et al. 2023). These multiple functionalities are associated with different domains of TniQ: the N-terminal domain interacts with TnsC, whereas the C-terminal domain mediates selective association with distinct CRISPR effectors (**Figure 1C**). This structural partitioning is reflected in TniQ’s sequence conservation: the N-terminal domain is more highly conserved, consistent with its role in promoting transposition, whereas the C-terminal domain exhibits variability (**Figure S2**). Based on these observations, we reasoned that retaining the N-terminal domain while designing a *de novo* C-terminal domain would enable engineering of a new-to-nature CAST that relies on a CRISPR effector of choice for targeting.

### NovoCASTs achieve fifty-fold increase in activity over PmcCAST

In total, four rounds of design were spent optimizing three different proteins (here, dCas9 and TniQ; in later sections, TnsAB) to create and optimize our new-to-nature CAST, which we term NovoCAST. We first computationally modeled the system by aligning dCas9 with PmcCAST target DNA (**Figure 2A**), retaining the N-terminal domain of TniQ (residues 1–233, **Figure 2A & S3**). To produce the *de novo* designs (**Figure 2B**), hereafter referred to as NovoQ, we implemented a two-stage computational pipeline. In the first stage, diverse backbone geometries were generated for NovoQ using RFDiffusion (Watson et al. 2023) with hotspots guiding the diffusion trajectories towards an exposed and accessible surface on dCas9 in the PAM-binding domain. This was followed with sequence design using LigandMPNN (Dauparas et al. 2025) in the context of dCas9, gRNA, and target DNA (**Figure 2C**). Additionally, five polar positions on the dCas9 binding surface were co-designed to allow for more favorable protein-protein interactions at the NovoQ–dCas9 interface (**Methods & Figure S4A).** Resulting designs were evaluated using physics-based scoring with Rosetta (Alford et al. 2017) and folding-based metrics with Chai-1 (Team et al. 2024), producing 441 top-scoring design candidates. The second stage iteratively refines sequence designs, using top-scoring candidates from stage one as seeds (**Figure 2C**) and prioritizing variants with high self-consistency between predicted and designed structures (**Methods & Figure 2D**). Across both stages, 2,000 structurally and sequence-diverse NovoQ designs (**Figure S5**) were selected for initial experimental screening. Selected designs feature well-packed hydrophobic cores, minimal buried unsatisfied hydrogen bonds, interface shape complementarity, and favorable predicted binding energy (**Figure S6**). Our computationally designed model of the fully assembled NovoCAST integration complex (**Figure S7**) predicts that integration events occur roughly 67 bp from the protospacer adjacent motif (PAM) sequence.

**Figure 2.**
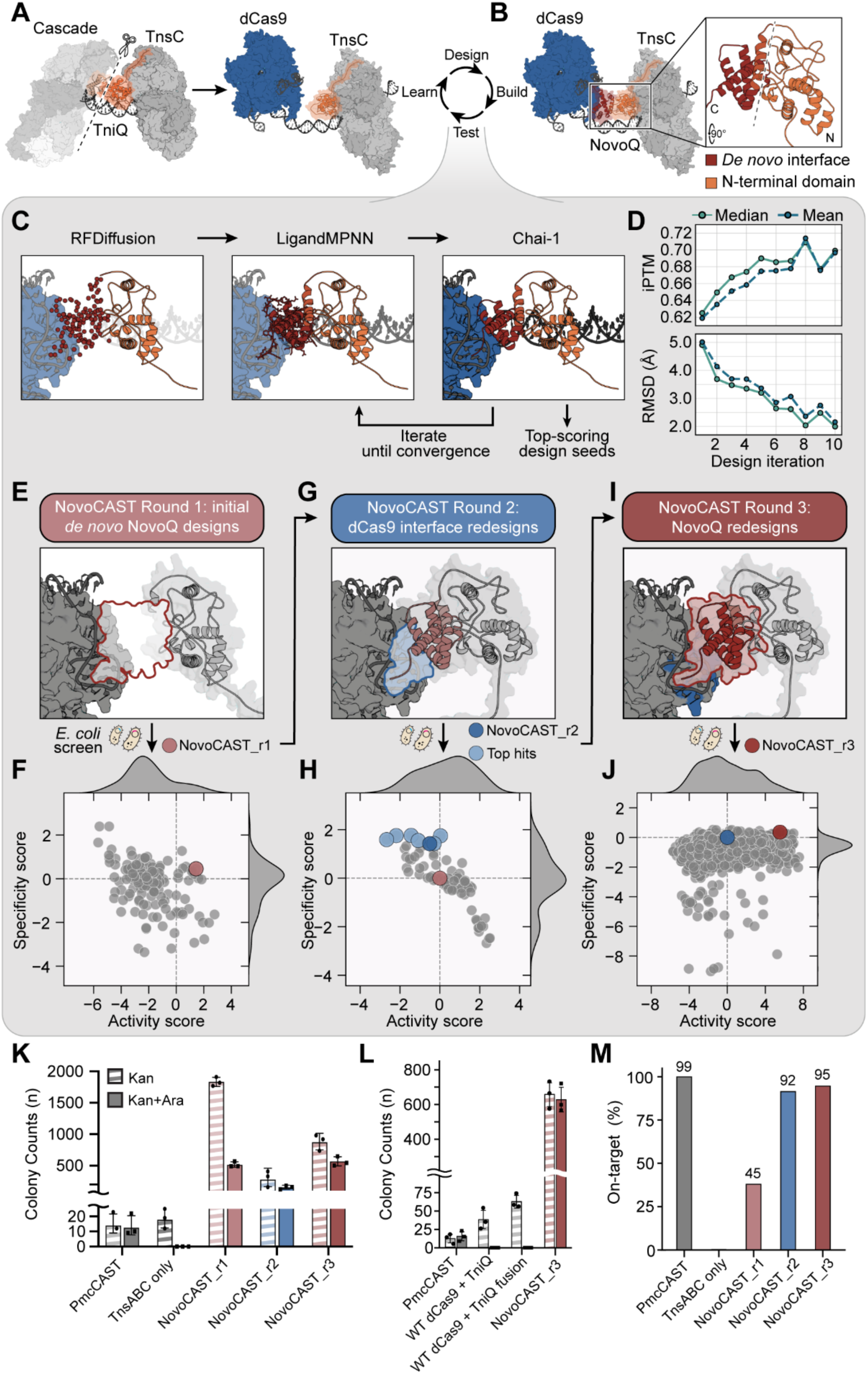
Iterative design–build–test–learn optimization of the NovoCAST system. **(A) Starting point for NovoCAST computational design.** PDB: 8FF4 (gray) is used as a starting point. Structural components are labeled. TniQ (colored as in Fig. 1) is truncated to remove its C-terminal domain (beige; indicated by scissors and a dashed line), retaining the N-terminal domain (orange ribbon). Target DNA is colored black. dCas9 (blue; PDB: 7OXA) docked on target DNA represents the starting point for design (black). **(B) Example of a complete designed system.** Inset: cartoon representation of NovoQ, with the N-terminal domain (orange) and *de novo* interface (red). The relative orientation with respect to overview is indicated by the rotation axis shown at bottom left. N and C indicate N-terminus and C-terminus, respectively. (C-J) Illustration of the design-build-test-learn pipeline. **(C) The computational design workflow.** Three stages are illustrated from left to right. **Left:** RFDiffusion generates *de novo* protein backbones, visualized as red spheres corresponding to sampled backbone coordinates that extend the fixed N-terminal domain (orange cartoon) to the putative binding surface of dCas9 (blue surface). **Middle:** LigandMPNN assigns amino acid sequences (shown in stick representation) onto the backbones of the diffused region, producing NovoQ sequence candidates with a putative dCas9-recognition domain (blue surface) at the target DNA (gray). **Right:** Chai-1 predicts the folded structures of the NovoQ-dCas9 complex, including NovoQ (red/orange), dCas9 (blue), and nucleic acids (black), enabling evaluation of the designed interface and structural plausibility. Designs are iteratively cycled through these steps (indicated by arrows), with top-scoring candidates selected as seeds for subsequent rounds until sequence-structure convergence is achieved. **(D) *In silico* design convergence across iterations.** Predicted interface TM-score (ipTM, top) and Root Mean Squared Deviation (RMSD in Angstroms, bottom) between designed and predicted models are shown as a function of design iterations (depicted in panel C). **(E) NovoCAST Round 1 strategy.** Red outline highlights the target of round 1: the *de novo* designed interface that extends the core region of NovoQ. **(F) Screening result of NovoCAST Round 1 designs**. Activity scores (x-axis) and specificity scores (y-axis) are shown for 76 NovoQ designs (gray points). Dotted lines indicate x = 0, y = 0. Conventions remain the same across panels F, H, and J. The top-performing NovoQ_r1-dCas9_r1 candidate (collectively, NovoCAST_r1) is colored rose. Histograms of activity and specificity scores are shown on the top and right sides of the plot, respectively. The *E. coli* icon indicates data from the bacterial screen. **(G) NovoCAST Round 2 strategy.** Target region for design is shaded in blue. An example of a NovoQ design is shown in cartoon for context (brown). **(H) Screening result of NovoCAST Round 2 dCas9 variants** (gray). The top seven dCas9 variants that were used for further optimization are shaded light blue. The final variant (dCas9_r2) that was selected from Round 3 screening is highlighted in dark blue. NovoCAST_r1(rose) is shown as a reference for comparison. **(I) NovoCAST Round 3 strategy.** The redesigned region of NovoQ is highlighted in red. **(J) Screening of NovoCAST Round 3 designs.** All designs shown in gray. NovoCAST_r3 is shown as a reference (dark blue). NovoQ_r3-dCas9_r2 (collectively, NovoCAST_r3), is highlighted in red. **(K) *In vivo* transposition assays in *E. coli* for top NovoCAST designs from each round.** Colony counts (n = number, y-axis) under kanamycin selection (striped bars) report overall integration activity, while kanamycin + arabinose selection (solid bars) report on-target integration activity. On the x-axis, transposition assays were performed under the same conditions individually for: PmcCAST, TnsABC only, NovoCAST_r1, NovoCAST_r2, and NovoCAST_r3 (shown left to right, respectively). The y-axis is broken to accommodate a large change in dynamic range. All data in panels (K-L) represent the mean ± standard deviation; n = 3 for each bar. **(L) Control transposition assays**. Transposition assays were performed under the same conditions individually for PmcCAST, WT dCas9-WT TniQ with TnsABC, a WT dCas9-WT TniQ fusion with TnsABC, and NovoCAST_r3 (shown left to right, respectively). **(M)** Genome-wide on-target specificity (percent on-target reads) across design rounds. Tagmentation assays were performed individually for PmcCAST, TnsABC only, NovoCAST_r1, NovoCAST_r2, and NovoCAST_r3 (shown left to right, respectively).

To determine relative integration activity and specificity of our NovoCAST Round 1 designs (**Figure 2E**), we used our previously developed bacterial screen (S. G. Park et al. 2025). Briefly, a kanamycin resistance cassette serves as the donor DNA (1.9 kb), such that kanamycin selection (Kan) reports overall integration activity. On-target integration is enriched by dual kanamycin and arabinose selection (Kan+Ara), which selects for disruption of the arabinose-inducible *ccdB* toxin gene. Activity and specificity scores are calculated as the log2-fold change in variant frequency compared to the input library. We observed robust and quantifiable on-target integration activity in 76 unique NovoQ-dCas9 designs (3.8% of the 2,000) (**Figure 2F**). We selected the NovoQ_r1 and dCas9_r1 pair as the first NovoCAST candidate (NovoCAST_r1) because this combination showed the strongest overall performance for on-target integration activity within the design pool. Next-generation sequencing (NGS) revealed an on-target specificity of 45.4% for NovoCAST_r1 (**Figure S8A**), compared to 99.9% for the best-performing PmcCAST gRNA (**Figure S9**).

The second round of design is based on the premise that strengthening the engineered NovoQ–dCas9 interface would enhance on-target specificity. Consistent with this, expanded redesign of the dCas9 interface (**Figure 2G**) yielded 89 variants, of which 32 (36%) improved on-target specificity without substantially compromising activity (**Figure 2H**; scores relative to NovoCAST_r1). Among the top-performing dCas9 variants (**Figure 2H**, light blue), the NovoQ_r1–dCas9_r2 pair (NovoCAST_r2; **Figure 2H**, dark blue **& S4B**) achieved 91.5% on-target specificity by genome-wide integration profiling (**Figure S8B**).

In a third round of design, we redesigned NovoQ_r1 in the context of the best-performing dCas9 variants from Round 2 (**Figure 2I**). We generated and screened 500 additional NovoQ designs against ten possible dCas9 variants (see Methods and **Figure S10**). Of 406 NovoQ designs with quantifiable on-target integration activity (**Figure 2J**; scores relative to NovoCAST_r2), 237 (58.4%) outperformed NovoQ_r1. We selected a Round 3 pair, NovoQ_r3–dCas9_r2 (NovoCAST_r3; red), for further characterization based on its increased activity (score = 5.54) relative to NovoCAST_r2 (**dark blue; Figure 2H**).

To validate the screen results, we compared the lead NovoCAST candidate from each design round to parental PmcCAST using *in vivo* bacterial transposition assays. Although PmcCAST has high targeting specificity (**Figure S9**), removal of Cascade reveals a latent TnsABC-driven off-target integration pathway (**Figure 2K**). Top NovoCAST candidates from design rounds 1–3 are 20–200-fold more active than PmcCAST, with the final design, NovoCAST_r3, achieving a 50-fold increase in activity while maintaining targeting specificity (**Figure 2K**). This is directly attributable to the designed NovoCAST interface, as neither simple substitution of Cascade for dCas9, nor fusion of dCas9 to PmcCAST TniQ, yields detectable on-target activity (**Figure 2L**). Consistent with our *in vivo* data, on-target integration specificity progressively improves across the three rounds of design, increasing from 45.4% for NovoCAST_r1 to 91.5% for NovoCAST_r2 (**Figure S8B**) and finally 94.6% for NovoCAST_r3 (**Figures 2M and 3A**). In agreement with our computational model (**Figure S7**), donor DNA integrated at a mean distance of 70 bp downstream of the PAM across all design rounds (**Figure 3B**), indicating that on-target integration is governed by the precise positioning of each component within the fully assembled transpososome. NovoCAST also exhibits striking orientation specificity, with 99.96% of integrations occurring in the L–R orientation relative to the PAM (**t-LR, Figure 3C**). This behavior mirrors that of its native precursor, PmcCAST, which inserts almost exclusively with its left end proximal to the attachment site (Wang et al. 2023).

**Figure 3.**
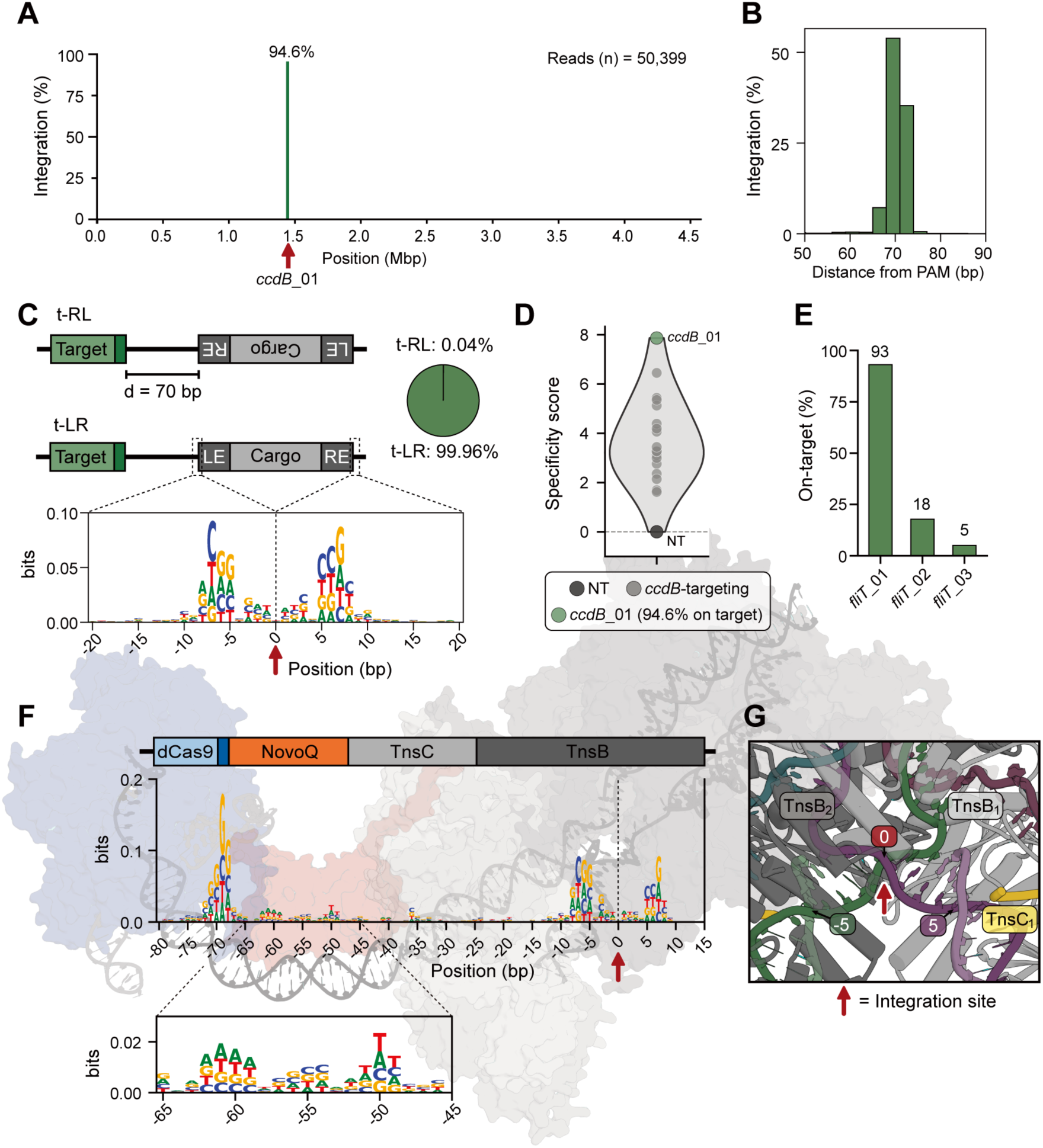
Sequence determinants of NovoCAST integration reveal dCas9-dependent, RNA-guided targeting with defined spacing, orientation, and PAM signatures. **(A) Genome-wide integration profile of NovoCAST_r3 mapped in *E. coli* using the *ccdB*_01 gRNA.** The x-axis shows genomic position (Mbp) and the y-axis shows the normalized mapped integration reads as integration (%). A strong peak with 94.6% of mapped reads is observed at the target locus, *ccdB_01* (red arrow) in *ccdB*. Data are from the same dataset shown in Figure 2M. **(B) Distribution of integration distances relative to the dCas9 PAM at the *ccdB*_01 target site.** Distances are calculated for the integration events shown in (A) with respect to the genomic position of the PAM. **(C) Diagram of the expected and observed simple insertion products for NovoCAST_r3.** The two possible simple insertion products, t-RL and t-LR, are diagrammed with RE, LE, and cargo indicated, dark and light gray respectively. Target sequence is illustrated in green, and the PAM motif colored dark green. The pie chart to the left illustrates orientation of integration events. The sequence logo below centers on the integration site (red arrow) for *fliT* off-target integration events, indicating local sequence preferences associated with the transposition machinery. The red arrow indicates the insertion position. **(D) A *ccdB*-targeting gRNA screen of NovoCAST.** Mean specificity scores over two replicates are shown on the y-axis. Specificity is normalized to the non-targeting control (NT), a non-targeting spacer that does not match any genomic sequence in *E. coli*, shown in black. *ccdB*_01 is highlighted in green and other *ccdB-*targeting guides are shown in gray. The violin plot illustrates the distribution of specificity scores for each gRNA. Dotted lines indicate 0 on the x- and y-axis. **(E) Tagmentation-based profiling of three gRNAs targeting the non-essential flagellar gene *fliT* in *E. coli*.** Tn5-based genomic profiling, on-target specificity of three guides: fliT_01 (93.1%), fliT_02 (17.8%), and fliT_03 (5.0%). **(F) Sequence logos derived from off-target integration windows for the *fliT*-targeting gRNAs, aligned relative to the integration site (red arrow).** The colored domains behind the main sequence logo indicates the expected DNA footprint of dCas9 (light blue), PAM (dark blue), NovoQ (orange), TnsC (light gray), and TnsAB (dark gray) for NovoCAST. A strong GG enrichment is observed approximately 65-70 bp upstream of the integration site, consistent with the location expected for a dCas9 PAM in a productive NovoCAST complex (scaled to match DNA footprint and shown in transparent surface in the background). The inset provides a detailed view of the region corresponding to the footprint of NovoQ (positions −65 to −45), highlighting the sequence signature observed from off-target integration events. **(G) Structure of integration site reveals protein-DNA interactions important for target DNA distortions.** TnsAB structure (PDB: 9BW1) shown in cartoon. Protein displayed in cylinder/ribbon, nucleic acid shown in cartoon. Two TnsB protomers (gray, labeled) shown at the integration site (indicated with red arrow). 0 indicates the same position as shown in panel F. The TnsC C-terminal tail (yellow) is positioned adjacent to TnsB_1_, making contacts near the active site in close proximity to the target DNA 5 bp away (labeled) from the insertion site, coinciding with the observed sequence preferences in panel F.

### NovoCAST Mediates Programmable, dCas9-Dependent Integration

To test the general programmability of NovoCAST, we screened 20 gRNAs targeting the *ccdB* reporter gene of our *E. coli* strain (S. G. Park et al. 2025). Consistent with CRISPR–Cas9 and native CAST systems (Doench et al. 2016; Strecker et al. 2019), integration specificity varied across gRNAs; however, every guide significantly outperformed the non-targeting (NT) control, highlighting the robust programmability of NovoCAST (**Figure 3D**). The most specific gRNA to *ccdB*, referred to as *ccdB*_01 (green, **Figure 3D**), is the same gRNA characterized earlier (**Figure 3A**). To assess targeting flexibility, we designed three gRNAs targeting the non-essential *E. coli* flagellar assembly gene, *fliT* (Juhas et al. 2014). Flagellar genes represent strategically advantageous insertion sites: though dispensable for growth, they govern motility and biofilm formation (Haiko and Westerlund-Wikström 2013) and rank among the most transcriptionally active and insertion-tolerant loci in the *E. coli* genome (Juhas et al. 2014). Across the three tested gRNAs, we observed integration specificities of 93%, 18%, and 5% (**Figures 3E and S11**), underscoring that gRNA choice is critical but does not completely determine targeting specificity.

We next sought to characterize the intrinsic sequence preferences of the NovoCAST transposase and their contribution to target-site selection. DDE transposons, a transposon superfamily encompassing Tn7-like elements, exhibit characteristic target-site sequence preferences which can strongly influence both target selection and integration positioning (Halling and Kleckner 1982). To this end, we generated sequence logos from genome-wide NovoCAST_r3 integration sites using the three *fliT*-targeting gRNAs (**Figure 3F**). Integration sites displayed two enriched motifs: a palindromic target motif (PTM) at the insertion site (position 0) and a PAM motif 70 bp away (position −70, **Figure 3F**). The PTM persisted in TnsABC-only insertions, whereas the PAM motif did not (compare panels A and B, **Figure S12**), indicative of intrinsic transposase-driven sequence preferences and a dCas9-dependent preference for the NGG PAM (Jinek et al. 2012). Target-site sequence preferences typically reflect transposase-DNA interactions critical for target-DNA bending and, ultimately, DNA integration (Arinkin et al. 2019). These results are consistent with prior work showing that target-site preferences are largely dictated by the transposase and can thus be used to guide the selection of optimal integration sites (George et al. 2023).

Intriguingly, closer examination of the strongly preferred off-target sites of NovoCAST identified clear gRNA-like sequences at the expected distance and orientation (**Figure S13)**. We reasoned that off-target integration hotspots likely arise from transient dCas9 stalling during target search, consistent with prior studies demonstrating that dCas9 occupies off-target sites with partial gRNA complementarity, particularly when the PAM-proximal seed region is intact (Kuscu et al. 2014; Wu et al. 2014). Supporting this idea, off-target sites contain putative gRNA matches within the PAM-proximal seed region (**Figure S13**).

### Sequence preferences reflect features of integration complex assembly

A central mechanistic question in CAST biology is why TnsC, an accessory AAA+ ATPase, is required for transposition when TnsAB catalyzes the strand-transfer reaction. In CAST systems, TnsC plays important roles in target-site selection (J.-U. Park et al. 2021) and in recruiting the transposase to the target site (J.-U. Park et al. 2022). Our findings raise the possibility that, in PmcCAST, TnsC may serve an additional role by directly contacting target DNA. Close inspection of the PTM sequence logo (**Figure 3C**) reveals strong sequence preferences 5-7 base pairs away from the insertion site, indicative of strong DNA distortions and, potentially, protein-DNA interactions. Consistent with this, the TnsC C-terminal tail in the previously published PmcCAST TnsABC structure (PDB: 9BW1) closely associates with bases at positions 5 and 6, coinciding with a sharp bend in the target DNA (**Figure 3G**), and removal of this portion of TnsC (residues 350 – 383) results in complete loss of transposition (Wang et al. 2023). We also observe a weak A-T preference at positions −62 to −49 (**Figure 3F**), coinciding with the presumed binding footprint of TniQ. Given that A-T rich sequences are associated with increased DNA flexibility and distortion, this preference suggested to us that TniQ binding may also bend target DNA.

### Cryo-EM structure of the NovoCAST_r3 complex confirms the accuracy of the designed assembly

To validate our *in silico* models, we determined the cryo-EM structure at 3.2 Å overall resolution (FSC = 0.143) of the NovoCAST_r3 complex harboring: NovoQ_r3, dCas9_r2, gRNA, TnsC, and target DNA (**Figure 4A, Figure S14**). We directly docked an AlphaFold3 predicted model of this assembly (**Figure S15**) into the density map and performed real space refinement followed by manual inspection and rebuilding (**Figure S16**). The structure (**Figure 4B**) confirms that NovoQ_r3 adopts the intended position and fold (**Ca RMSD = 1.13 Å, Figure 4C**) between dCas9_r2 and TnsC **(global Ca RMSD = 2.1 Å, Figure S17**). Compared to the parental protein (PmcCAST TniQ), the NovoQ_r3 C-terminal domain adopts a distinct fold (TM-score = 0.43, **Figure S18**). Importantly, our cryo-EM structure reveals a hydrophobic pocket of interactions at the NovoQ_r3–dCas9_r2 interface (**Figures 4C & 4D**), mainly stabilized by NovoQ_r3 residues: Y306, L313, F315, I319, M310, A245, L249, V248, L252 and dCas9_r2 residues: I1196, L1194, L1191, V1146, and L1148. Thus, the NovoCAST_r3 design folds as predicted, providing strong validation of our design strategy. In addition to validating the designed NovoCAST architecture, the structure also reveals DNA distortions associated with the previously mentioned NovoQ binding footprint **(inset, Figure 3F).** Our top design displays a degree of target DNA bending reminiscent of that observed in the PmcCAST complex (**Figure S19**) (Wang et al. 2023). DNA distortions have previously been implicated as regulatory features of Tn7-like transposases to facilitate recruitment and assembly of the transposition machinery (Kuduvalli et al. 2001). The similarity of DNA distortions suggests that NovoCAST may retain mechanistic features of Tn7-like transposases to facilitate efficient integration.

**Figure 4.**
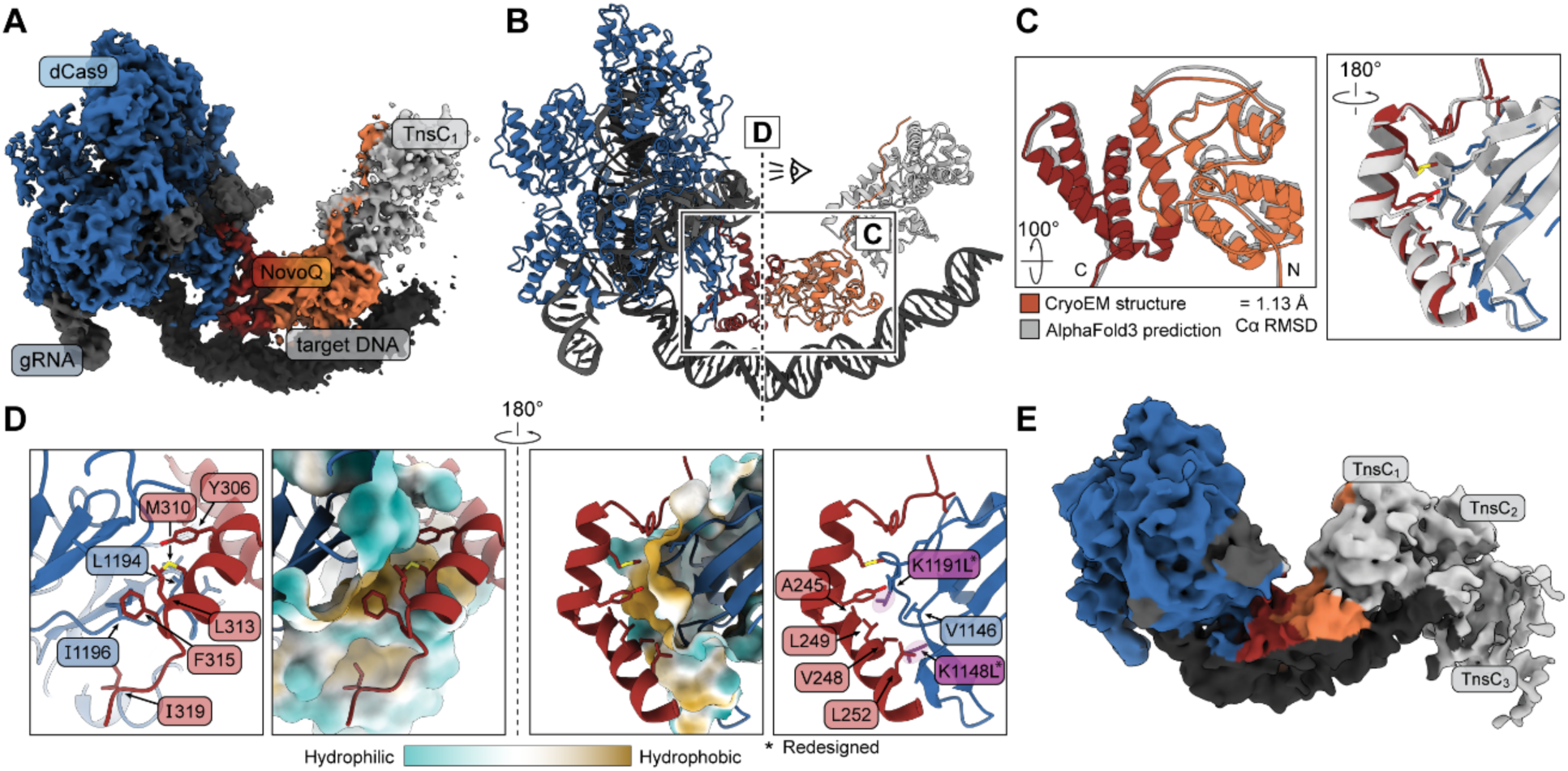
Cryo-EM structure of the NovoCAST_r3 complex confirms accuracy of NovoCAST designs. **(A) High-resolution cryo-EM density map of the NovoCAST_r3 complex**, consisting of dCas9_r2 (blue), NovoQ_r3 (orange/red), gRNA (gray), target DNA (black), and one resolved TnsC protomer (TnsC_1_, light gray). **(B) NovoCAST_r3 complex atomic model.** Colors match that of panel A. Structure is shown in cartoon. The boxed region indicates the detailed view of the NovoQ-dCas9 interface shown in panels **(C-D).** Dotted line and eye indicate viewing direction for panel D. **(C) Comparison of NovoQ_r3 predicted and experimentally determined structure.** Overlay shows AlphaFold3 predicted complex structure (white) superimposed on experimentally determined structure (colors as in panels A-B). Cα RMSD indicated below the structural superposition (1.13 Å). Rotations are with respect to the overview in panel B. N and C-termini are labeled N and C, respectively. **(D) Detailed view of NovoQ_r3-dCas9_r2 interface.** Representative side chains (far left and far right panels) on NovoQ_r3 (red) and dCas9_r2 (blue) that contribute to interface packing are displayed as sticks from two viewpoints rotated by 180°. NovoQ_r3 residues displayed include A245, V248, L249, L252, Y306, M310, L313, F315, and I319. dCas9_r2 residues displayed include: V1146, L1194, I1196, K1148L, and K1191L, where K1148L, and K1191L represent two of the designed substitutions positioned directly at the interface. Redesigned dCas9 residues are indicated with *. The center panels show a surface representation of the refined model of dCas9_r2 colored by hydrophilicity/hydrophobicity (scale shown on bottom). **(E) Cryo-EM density of NovoCAST_r3 reveals three TnsC protomers (TnsC_1_-TnsC_3_).** A subset of particles containing a more complete TnsC ring resulted in a 7.3 Å cryo-EM map, shown as a colored surface. Colors follow conventions laid out in panel A. The additional density in the low-resolution map corresponds to TnsC protomers TnsC_2_ and TnsC_3_.

Downstream of the engineered NovoQ–dCas9 interface, we identify clear density corresponding to a TnsC protomer bound at the expected position (**Figure 4A**). Further 2D classification identifies a subset of particles with additional density corresponding to TnsC (**Figure S14**). These particles were used to reconstruct a 7.3 Å cryo-EM map, sufficient to visualize a total of 3 TnsC protomers bound to target DNA (**Figure 4E**). Indeed, the arrangement of three TnsC protomers docked in this map matches the helical parameters expected for the canonical heptameric TnsC assembly observed in its precursor, PmcCAST (Wang et al. 2023) (**Figure S20**), indicating that NovoCAST supports both recruitment and nucleation of the TnsC filament architecture required for transposition.

### Programmable Genomic Integration in Human Cells Using NovoCAST

To assess the potential of NovoCAST for bioengineering and therapeutic applications, we evaluated its capacity for PGI in human cells. We constructed mammalian expression vectors encoding NovoCAST_r3 and co-transfected them into HEK293FT cells with a donor plasmid (∼2.1 kb cargo) to perform both episomal (plasmid-to-plasmid, using a co-transfected pTarget containing *ccdB*) and genomic (plasmid-to-genome) integration assays (**Figure 5A**). We could detect *ccdB*-targeted episomal integration mediated by NovoCAST_r3 by junction PCR (**Figure 5B**), with the expected integration spacing profiles (**Figure 5C**). These results demonstrate that NovoCAST retains functional activity in a mammalian context.

**Figure 5.**
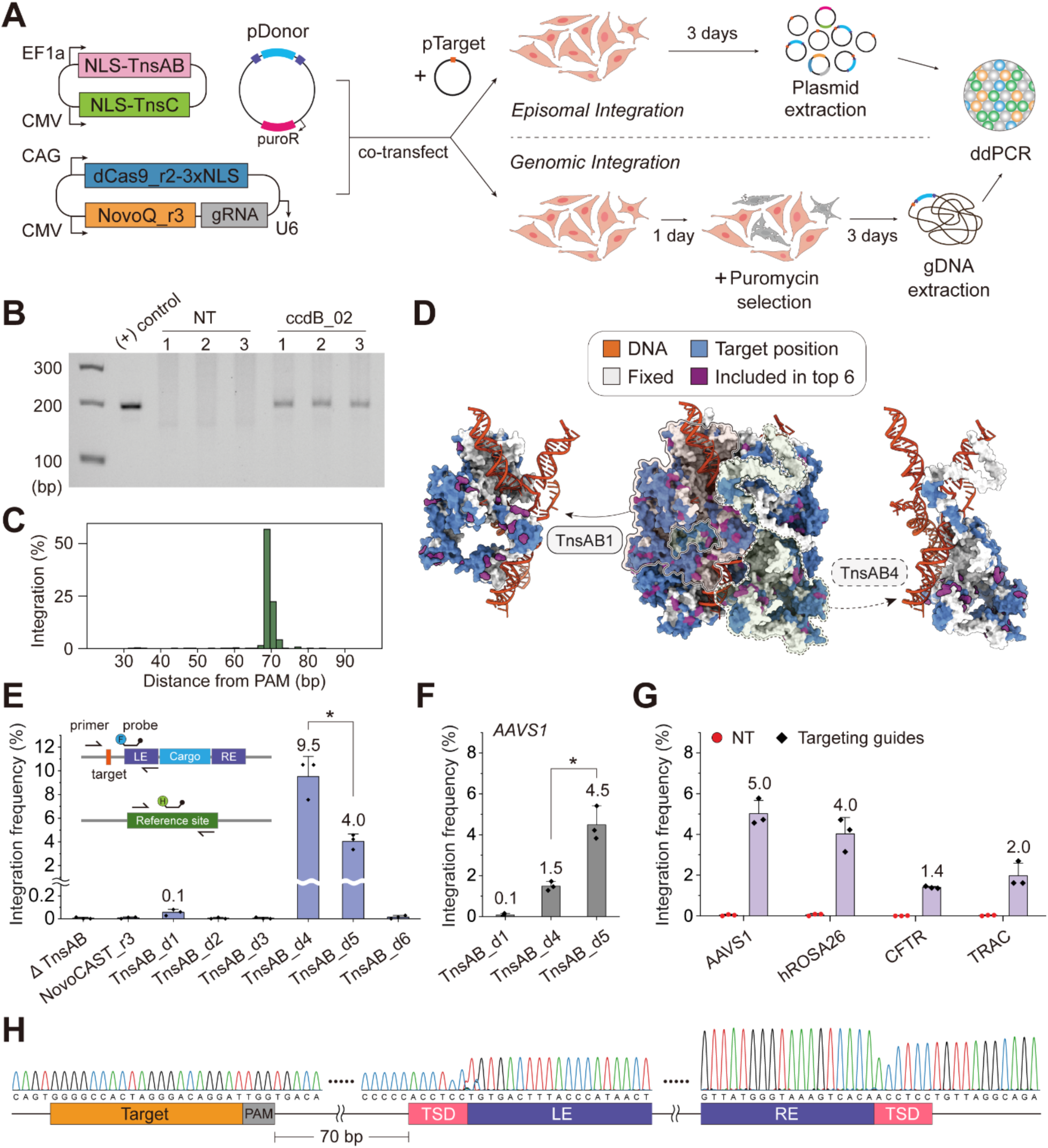
NovoCAST can efficiently integrate donor DNA into the target site in human cells. **(A)** Schematic illustration of episomal integration (top left) and genomic integration assays (bottom left) in human cells. Mammalian expression vectors encoding NovoCAST components with NLS tags under separate promoters were co-transfected with a pDonor plasmid into HEK293FT cells. For the episomal integration assay, a pTarget plasmid harboring the target site (orange) was additionally co-transfected. For the genomic integration assay, cells underwent puromycin selection to remove non-transfected cells. Plasmids or gDNAs were extracted from transfected cells, and integration frequencies were quantified by ddPCR. **(B)** Agarose gel image showing junction PCR of the LE junction. The sizes of the DNA ladder (lane 1) are indicated on the left side. (+) control represents a mock integration sample. NT, non-targeting guide; *ccdB*_02. Lane 3-5 and Lane 6-8 represent triplicates. **(C)** Distribution of integration distances relative to the PAM at the target site. Distances were calculated for the integration events shown in (B), relative to the position of the PAM on the pTarget-*ccdB* sequence. **(D)** Redesign target residues mapped onto the AlphaFold3-predicted atomic model of TnsAB in the DNA-bound homo-tetramer state. White, fixed; blue, redesign targets; purple, redesigned positions in the top 6 designs; orange, DNA. Individual structures of TnsAB1 and TnsAB4 protomer extracted from the tetramer are indicated by arrows. **(E)** Episomal integration frequencies of NovoCAST with six TnsAB redesigns, shown together with a schematic of the ddPCR assay. Integration frequency was quantified using junction-specific primers and a probe (F, FAM) and normalized to reference (pTarget backbone or RPP30) site-specific primers and a probe (H, HEX). * *p* = 0.014 (unpaired two-tailed Welch’s t-test). **(F)** Quantification of AAVS1-targeted integration in HEK293FT cells expressing TnsAB_d1, TnsAB_d4, or TnsAB_d5. βTnsAB represents a deletion of TnsAB. * *p* = 0.026 (unpaired two-tailed Welch’s t-test). **(G)** Genomic integration frequencies of NovoCAST_r4 across four different target loci. All data in panels **E-G** represent the mean ± standard deviation; n = 3 for each bar. WT, wild type; NT, non-targeting gRNA; The *AAVS1* integration data were generated independently from those shown in panel **F**. **(H)** Sequencing chromatograms of LE and RE junction PCR products derived from AAVS1-targeted integration (panel G). Each colored trace represents a nucleotide: green, A; red, T; blue, C; black, G. The target site, PAM, LE, RE, and TSD are indicated in boxes below the corresponding sequences. Wavy lines indicate omitted regions within the insertion window (70 bp) and portions of the donor DNA due to space constraints.

However, unlike in *E. coli*, the integration activity of NovoCAST_r3 in human cells was below the limit of detection by ddPCR. This could be due to the instability of TnsAB in human cells, as western blot analysis showed low expression levels (lane 9; **Figure S21A**), and biochemical purification revealed substantial aggregation (**Figure S22**). Since transposase engineering can substantially improve integration activity (S. G. Park et al. 2025), we pursued an additional round of computational protein design (Round 4), focused on improving TnsAB stability and function. We used LigandMPNN (Dauparas et al. 2025) to stabilize the active conformation of TnsAB as predicted by AlphaFold3 (“Accurate Structure Prediction of Biomolecular Interactions with AlphaFold 3 | Nature,” n.d.) (**Figure 5D)**. Because structure-based sequence optimization can over-stabilize at the expense of activity (Sumida et al. 2024), we included per-residue sampling bias towards active TnsAB homologs (**Methods**). Residues distal from DNA, active sites, disordered loops, or highly conserved regions were subjected to redesign (**Figure 5D**).

Given the absence of a suitable high-throughput assay in human cells, we directly tested six top-scoring TnsAB designs, each containing between 13–36 mutations (purple; **Figure 5D**), using our episomal integration assay followed by ddPCR (**Figure 5A**). Remarkably, three out of six designs significantly outperformed NovoCAST_r3 (**Figure 5E**), supporting the idea that TnsAB is a promising engineering target. The top-performing TnsAB variants TnsAB_d4 and TnsAB_d5, achieved episomal integration efficiencies of 9.5% and 4.0%, respectively, as measured by ddPCR (**Figure 5E**).

Next, we investigated whether NovoCASTs with TnsAB redesigns are capable of performing PGI in human cells (**Figure 5A**). As a proof of concept, we targeted the *AAVS1* safe-harbor locus using NovoCAST with either TnsAB_d4 or TnsAB_d5. We obtained genomic integration efficiency of 1.5% with TnsAB_d4 and 4.5% with TnsAB_d5 (**Figure 5F**). To examine whether the observed improvements reflect intrinsic properties of the redesigned transposase or host-specific effects, we tested these variants in *E. coli* and observed a consistent 2–10-fold increase in activity relative to NovoCAST_r3 (**Figure S23**), albeit with slight reduction in specificity (**Figure S24**). The enhancement appears to be driven primarily by increased intrinsic activity rather than by elevated expression, since the designs have comparable expression levels in human cells (**Figure S21B**). We also tested whether our redesigns affected TnsA catalytic activity. TnsA is an endonuclease that, in PmcCAST, is fused to TnsB and required for transposition (Saito et al. 2021); its catalytic activity promotes simple insertions over cointegrates by specifically cleaving flanking genomic DNA. NovoCAST_r4 produced 15.7% cointegrates in *E. coli* (**Figure S25**), comparable to native PmcCAST (Saito et al. 2021), suggesting that TnsAB_d5 preserves this key functional property.

The top design, NovoCAST_r4, is created with three engineered proteins: NovoQ_r3, dCas9_r2 and TnsAB_d5. NovoCAST_r4 shows ∼500-fold higher activity in *E. coli* compared to PmcCAST, reflecting a ∼50-fold improvement over NovoCAST_r3 (**Figure 2K**) and an additional ∼10-fold improvement from the optimized TnsAB_d5 (**Figure S23**). Finally, to demonstrate the potential therapeutic relevance of NovoCAST, we evaluated an additional safe-harbor locus (*hROSA26*) as well as two therapeutically relevant loci in *CFTR* (cystic fibrosis transmembrane conductance regulator) and *TRAC* (T cell receptor alpha constant). We chose sites in the first intron or upstream of exon 1, which are commonly targeted for therapeutic applications. NovoCAST_r4 achieved 5.0% and 4.0% integration frequencies at *AAVS1* and *hROSA26*, and 1.4% and 2.0% at *CFTR* and *TRAC*, respectively (**Figure 5G**). No integration was detected in non-targeting (NT) controls, confirming that the observed integration is driven by RNA-guided activity of NovoCAST. Sequencing chromatograms of the LE and RE junctions at the *AAVS1* locus showed integrations occurring 70 bp downstream of the PAM (**Figure 5H**). A target site duplication (TSD), clear evidence of transposase-mediated integration, was also detected (pink box; **Figure 5H**). Fluctuations at the TSD and junction boundaries (both LE and RE junctions) reflect slight variation in the insertion window (69-71 bp) and TSD length (5-6 bp). Collectively, these results indicate that NovoCAST_r4 maintains consistent integration features across both bacterial (*E. coli*) and mammalian (human) systems.

## Discussion

Our results establish computationally designed NovoCAST as a programmable, new-to-nature, large-cargo (≥ 2 kb) genome integration tool that significantly outperforms natural CAST systems in bacterial and human cells. CAST systems have previously required extensive screening, vector and tag optimization, and the introduction of putative host factors such as ClpX to achieve low levels of integration (1–5%) in human cells (Liu et al. 2025; Lampe et al. 2024). Remarkably, our best designs match these efficiencies without such optimization. Although integration efficiencies remain ∼10-fold lower than those of evolved CAST systems (10–30%), we anticipate that standard optimizations for human cell activity, such as NLS tag engineering (Suzuki et al. 2016), could significantly improve activity further.

Cumulatively, our designs achieve ∼500-fold higher activity than the natural system, comparable to gains reported from directed evolution approaches (Witte et al. 2025). In contrast to approaches based on bioinformatic discovery and directed evolution, a rational computational protein design approach enables specific customization of properties, such as size and architecture, and results in transferable design principles, while avoiding the unknown variables (such as host factors) that often constrain the portability of natural CAST systems. The modular use of well-characterized components such as dCas9 is an added benefit, enabling straightforward extension of the system to altered PAM specificities (Kleinstiver et al. 2015) and incorporation of activity-enhancing variants (Vakulskas et al. 2018; Ikeda et al. 2019; Kleinstiver et al. 2016). Together, these results establish rational computational design as a compelling alternative to prevailing optimization strategies.

CRISPR effector replacement yields the simplest CAST to date, comprising only four components, with further simplification likely achievable. The observation that TniQ’s N-terminal domain, and its positioning within the assembly, are important for transposition illustrates how mechanistic understanding of biological systems can directly inform structure-based engineering strategies. Accordingly, smaller effectors, such as TnpB (Altae-Tran et al. 2021), may provide a route to additional minimization, and the N-terminal domain of TniQ could potentially be incorporated directly into the effector. As in other CAST systems, TnsC is required for transposase recruitment and activation at the target site (J.-U. Park et al. 2022; Hoffmann et al. 2022). Here, we identify a region of TnsC that may stabilize target DNA distortion, a role typically attributed to the transposase. These findings raise the possibility that the transposase and TnsC cooperatively remodel target DNA, which may open new avenues for further development of novel CAST systems.

Other engineered PGI tools, including dCas9 transposase fusions (Kovač et al. 2020) or even fusions with CAST transposition machinery (Schargel et al. 2025) display promiscuous targeting, likely arising from loose coupling of independently functional modules via unstructured linkers. Although integration in our system appears to be tightly coupled to dCas9, we nevertheless observe off-target activity. Off-targets appear to be driven by partial matches to gRNA, which are commonly observed in dCas9 genome-wide binding profiles (Wu et al. 2014). These observed off-target profiles likely arise from the positioning of the NovoQ–dCas9 interface which is distal to R-loop sensing. Therefore, achieving greater targeting specificity will require both the repositioning of TniQ and the incorporation of negative design principles (Davey and Chica 2012) to discriminate between R-loop-dependent conformational states of dCas9, as in natural CAST systems (J.-U. Park et al. 2023; Wang et al. 2023).

Close agreement between our top design and the experimental cryo-EM map validates the accuracy of the design approach and its underlying hypotheses. Although target DNA distortion was not an explicit design objective, it is present in our top NovoQ designs and is observed in the experimental structure. DNA distortion has previously been shown to be required for assembly of the ShCAST integration complex (J.-U. Park et al. 2023). The emergence of this feature in our active designs supports the idea that target DNA deformation may be a prerequisite for functional coupling in these systems. However, the precise role of DNA distortion remains incompletely understood and will require further investigation to elucidate how DNA structure might be coupled to biological activity in these systems.

Robust integration activity in human cells was achieved by jointly optimizing the CRISPR effector and stabilizing the transposase (TnsAB). Transposition proceeds through an isoenergetic reaction driven by increasing the stability of the nucleoprotein assembly (Yin et al. 2012, 2016; Montaño et al. 2012). On this basis, we hypothesized that the post-integration complex would represent the most appropriate target for design and find that this strategy yields substantial gains in activity. Notably, the largest improvements in human cells arise from TnsAB redesign despite limited exploration of sequence space, with only six constructs tested. Expanded exploration of TnsAB redesign, combined with standard optimization approaches and continued study of nucleosome-dependent effects on integration, is likely to further improve translational application of CAST systems to human genome-editing.

The power of structure-based computational protein design lies in its ability to encode biological function from first principles; a capability that distinguishes it from large language models, which can generate sequence diversity but cannot rationally encode novel function. Our design strategy imposes stringent constraints: the *de novo* protein must preserve allosteric function, stably engage dCas9, and maintain a well-folded core while accommodating a domain from the precursor, PmcCAST. The success of this design approach provides, to our knowledge, the first demonstration that *de novo* protein design can orchestrate a coordinated molecular process in a stepwise manner within a genome-editing system. Furthermore, our results suggest that designing coordinated molecular machines, rather than loosely linked assemblies, is a promising and viable avenue to developing highly specific and active PGI tools.

In conclusion, the convergence of mechanistic insight into RNA-guided systems with rapid advances in protein design establishes a foundation for the *de novo* engineering of genome-editing technologies. We envision that these systems can be further deconstructed into modular domains, creating a “parts” library from which RNA-guided systems with tailored properties and novel biological functions can be assembled. This framework opens new opportunities in areas such as RNA editing and epigenome editing, where clinically viable tools remain limited. Beyond size, properties such as immunogenicity and manufacturability can be precisely tuned through design. Advances in DNA synthesis and assembly further suggest that increasingly complex architectures will soon be accessible (Hoose et al. 2023), expanding the scope of what can be engineered. Together, we believe these developments mark a transition from discovering genome-editing systems in nature to designing them with intent.

## Data Availability

The cryo-EM map and atomic coordinates reported in this study have been deposited in the Electron Microscopy Data Bank (EMDB) and the Protein Data Bank (PDB) under accession codes **EMDB: EMD-76812** and **PDB: 12WP** (NovoQ–dCas9–TnsC complex) and **EMDB: EMD-77211** (NovoQ–dCas9–TnsC, low resolution map with TnsC oligomerization). The raw high-throughput sequencing data are available from NCBI SRA (**accession PRJNA1468775**). All other data supporting the findings of this study are available within the paper and its Supplementary Information.

## Contributions

E.H.K. and A.Z. conceived and supervised the research and contributed to writing the manuscript. H.M.W. conducted bioinformatic analyses. H.M.W. developed design pipelines for NovoCAST design rounds 1-3 and the regression models. A.S. designed and generated plasmid libraries for design rounds 1-3 and gRNA screens. S.G.P. and A.S. performed high-throughput screens and transposition assays in *E. coli*. A.S. and H.M.W. designed and analyzed guides for *fliT*-targeting guide screen. S.G.P., A.S., S.V., and H.M.W. performed and characterized integration profiles. H.M.W. and E.C. analyzed off-target hotspots. S.G.P. designed and performed human cell experiments. S.G.P. and T.S. carried out ddPCR. S.F. and S.D. purified NovoCAST components, performed DNA-based pulldowns, and prepared cryo-EM grids. S.D. collected the cryo-EM data. H.M.W. and S.D. processed, analyzed, and refined the atomic models. H.M.W. generated sequences for TnsAB designs. H.M.W. and S.G.P. contributed to writing the initial manuscript draft. All authors contributed to manuscript writing.

## Acknowledgements

We thank Shengdar Tsai, Kasey Jividen, Jackie Chyr, and Azusa Matsubara for helpful discussions related to vector design, genomic profiling, characterization of integration events, and experimental feedback in human cells. We thank Esteban Dodero-Rojas for initial discussion on the design strategy and pipeline support. We thank Inês Chen, M Madan Babu, Babis Kalodimos, Shengdar Tsai, Eli Fritz McDonald, Rian Kormos, Sitaram Gayatri, and Yun-Nam Choi for valuable feedback during writing. This research is supported by the Pew Charitable Trusts, NIH NIGMS 7R01GM144566, and the Cystic Fibrosis Foundation (E.H.K.). We also thank the St. Jude Hartwell Center for Bioinformatics & Biotechnology which is supported in part by ALSAC and the National Cancer Institute grant P30 CA021765 for their help conducting next-generation sequencing, the St. Jude Cryo-EM Center, and St. Jude Research Information Services for providing high-performance scientific computing systems and computational resources that contributed to the research in this manuscript. The funders had no role in study design, data collection and analysis, decision to publish, or preparation of the manuscript.

## Methods

### Phylogenetic analysis of CAST TniQ proteins

To identify TniQ homologs that are associated with CAST systems, CAST gene neighborhoods and gene accession codes were first obtained from a previously reported CAST database (Rybarski et al. 2021). For genomes lacking complete protein annotations, open reading frames were predicted and translated using Prodigal (Larralde 2022). Because the source CAST dataset contained annotation noise and multiple putative TniQ annotations per CAST gene neighborhood, all predicted proteins in the dataset were independently re-screened using a hidden Markov model (HMM) corresponding to the conserved TniQ domain (PF06527) that was constructed from TniQ proteins from functionally characterized CAST types (Types I-B, I-F, and V-K). Candidate proteins were screened using HMMER v3.4 (Finn et al. 2011) with default settings, and the corresponding matches were filtered using a domain E-value threshold of ≤ 1e-3 and a domain bit score ≥ 25.

Because TnsD proteins also contain a conserved TniQ domain but function as sequence-specific DNA-targeting proteins rather than recognizing RNA-guided CRISPR effectors, candidate proteins were additionally filtered by protein sequence length. Proteins exceeding the characteristic length of canonical CAST TniQ proteins (>300 amino acids for type V-K; >450 amino acids for types I-B and I-F) were removed from the dataset. Candidate proteins lacking recognizable CRISPR effectors and CAST-associated proteins were also excluded. Each retained TniQ homolog was mapped to its associated CRISPR effector, and selected sequences were validated using CRISPRCasTyper (Russel et al. 2020). TniQ homologs associated with non-canonical CAST types or divergent CRISPR systems (Types I-C, IV, or unclassified) were removed from the final dataset to restrict the analysis to well-characterized CAST types.

TniQ sequences corresponding to CAST Types I-B2, V-K, and I-F3a were used as queries to identify additional homologs using PSI-BLAST (NCBI BLAST+ v2.17.0) (Altschul et al. 1997) against the nr database (version 11/25/2025) using an E-value of 0.005, converging by the third iteration. These additional homologs were incorporated into the final alignment to anchor the major phylogenetic clades. Redundant sequences were removed using CD-HIT with a 99% sequence identity threshold (Li and Godzik 2006). The resulting non-redundant TniQ sequences were aligned using MAFFT with default parameters (Katoh and Standley 2013). The maximum likelihood phylogenetic tree was constructed using PhyML (Guindon et al. 2010), and the tree visualization was generated using the Interactive Tree of Life (Letunic and Bork 2021). The final dataset used for phylogenetic analysis contained 709 CAST-associated TniQ sequences is provided in **Supplementary Table 1**.

### Computational design of NovoQ

#### Starting model

The crystal structure of *Streptococcus pyogenes* Cas9 bound to target DNA with a chimeric RNA-DNA guide (PDB: 7OXA) was used as the dCas9 scaffold for NovoQ binding and design. The cryo-EM structure of the Type I-B2 CAST (PmcCAST) Cascade–DNA–TniQ–TnsC complex (PDB: 8FF4) was used as the source of the TniQ scaffold and target DNA geometry.

To generate the starting model for NovoCAST design, the target DNA from the dCas9 scaffold was superimposed onto the target DNA from the PmcCAST structure using `super` command in PyMOL (version 3.1.7.2) (Schrödinger, LLC, n.d.). Following DNA superposition, the Cascade complex was removed while retaining the TniQ scaffold and target DNA coordinates from the PmcCAST structure. The resulting models were analyzed to ensure: 1. preservation of the target DNA geometry observed in PmcCAST, 2. an exposed and accessible interaction surface on dCas9 for NovoQ recognition, and 3. sufficient spatial separation between dCas9 and the retained TniQ scaffold to accommodate a de novo designed C-terminal domain (**Figure S3**). The resulting superimposed model positioned the N-terminal domain of TniQ and dCas9 approximately 20 Å apart and was used as the starting template for subsequent computational protein design.

#### Backbone generation

To generate initial NovoQ backbones for de novo interface design, the PmcCAST TniQ scaffold from the starting model was truncated to remove the native C-terminal effector-recognition region (residue positions 309-329), retaining residues 1-308. To reduce computational cost during diffusion-based backbone generation, the dCas9 scaffold was trimmed to a contiguous region encompassing the intended NovoQ interaction surface and the surrounding local structural environment. The retained dCas9 residues corresponded to positions 1-57 and 1094-1365.

Because the dCas9 target DNA and gRNA occupy space directly adjacent to the intended NovoQ interaction surface, and nucleic acids could not be included with the RFDiffusion input model at the time, a placeholder poly-glycine chain was generated and positioned along the nucleic acid coordinates to sterically occlude these regions during diffusion. This placeholder chain was generated by tracing glycine residues along the nucleic acid phosphate backbone coordinates and was included only during the backbone diffusion stages to discourage diffused protein backbones from occupying the nucleic acid space.

New protein backbones extending the truncated TniQ C-terminus by 8-24 residues were generated using RFDiffusion (Watson et al. 2023). Diffusion trajectories were guided using hotspot residues defined on the dCas9 interaction surface (residues 1194, 1179, 1161, 1191, 1174, and 1360) and the TniQ scaffold (residues 250 and 253), producing several thousand TniQ backbone models with de novo C-terminal extensions per design campaign. The resulting backbone models were clustered according to the structural similarity of the diffused C-terminal extension using TM-align (Zhang and Skolnick 2005) with a TM-score threshold of 0.4, typically yielding 50-100 structural clusters per design campaign. Representative models from each cluster were selected based on interface shape complementarity and buried solvent-accessible surface area (dSASA), calculated using Rosetta (Alford et al. 2017; Leaver-Fay et al. 2011; Fleishman et al. 2011) (version 2024.09+release.06b3cf8).

Selected cluster representatives were subsequently subjected to 100 rounds of partial diffusion using RFDiffusion with residues 234 through the C-terminus designated as the diffused positions with `diffuser.partial_T=20` (Glögl et al. 2024). The final C-terminal position varied between residues 316 and 332 depending on the generated backbone length. The same dCas9 hotspot residues used during the initial diffusion stage were retained during partial diffusion. These steps generated a structurally diverse ensemble of plausible candidate backbones used for the initial NovoQ design set.

#### Sequence design

Before sequence design, the poly-glycine nucleic acid placeholders used during the backbone diffusion steps were removed and replaced with the original gRNA and target DNA coordinates from the starting model. Sequences were generated using LigandMPNN (Dauparas et al. 2025) (2 sequences per backbone model). Residues 1-233, corresponding to the N-terminal TniQ domain, were held fixed. Residues 234 through the C-terminus (which varied between residues 316 and 332 depending on NovoQ length), were fully designed in presence of the dCas9, gRNA, and target DNA. In Round 1, five surface-exposed polar residues on the dCas9 surface predicted to become buried upon NovoQ binding (K1148, K1161, K1191, K1192, and D1344; **Figure S4A**) were also allowed to vary during co-design with NovoQ.

#### Design filtering

Each design was first idealized using the `IdealizeMover` in PyRosetta, and residues within 10 Å of the dCas9–NovoQ interface were relaxed using the `FastRelax` mover in PyRosetta (Alford et al. 2017; Chaudhury et al. 2010) (version 2024.42+release.3366cf78a3). Designs were retained based on the following Rosetta-based interface and packing thresholds: Rosetta ddG/dSASA < –2, Rosetta ddG < –45, interface hydrophobicity > 60%, interface shape complementarity > 0.5, Rosetta holes < 0.2, Rosetta packstat > 0.47, buried unsatisfied atoms < 10, and dSASA > 2100. Passing designs were then predicted using Chai-1 (Team et al. 2024) as both NovoQ monomers and NovoQ–dCas9–gRNA–DNA complexes. Designs were further filtered according to the following thresholds: monomer Cα root mean square deviation (RMSD) of monomer < 3 Å, complex C-terminal extension RMSD < 4 Å, NovoQ displacement after aligning on dCas9 < 4 Å, mean RMSD across five Chai-1 predictions < 10 Å, monomer predicted template modeling (pTM) > 0.8, and complex interface pTM (ipTM) > 0.8.

Filtered sequences were clustered with MMSeqs2 (Steinegger and Söding 2017) at 99% identity, yielding 441 unique NovoQ designs. These were denoted as top-scoring design seeds (**Figure S6, blue**) and subjected to iterative redesign of the C-terminal domain of each predicted fold for up to 10 cycles (**Figure S6, pink**). Designs showing score improvement at each cycle were retained. Including iterative redesigns, a total of 2,000 NovoQ sequences were selected for Round 1 screening (**Supplementary Table 2A**), along with the five most frequently sampled dCas9 variants observed across the design output (**Supplementary Table 2B**). The Round 1 screening results are provided in **Supplementary Table 2C**.

#### Subsequent design rounds 2–3

Rounds 2 and 3 omitted the initial de novo backbone generation steps performed in Round 1 and instead used the top-performing Round 1 design, NovoQ_r1, as the starting backbone for subsequent redesign and optimization.

In the Round 2 dCas9 variant designs, NovoQ_r1 was subjected to partial diffusion for 5 and 10 diffusion steps (`diffuser.partial_T=5; diffuser.partial_T=10`) while retaining NovoQ_r1 sequence information. Interface residues on the dCas9 surface located within 4 Å of NovoQ_r1 in the computational model were redesigned using LigandMPNN. Resulting dCas9 design variants were filtered and iteratively redesigned using the same Rosetta and structure-prediction-based methods described above. The Round 2 dCas9 designed variant library consisted of 5 sequences designed directly following partial diffusion, 12 sequences from iterative redesign, and 72 sequences constructed by combinatorially sampling mutations identified from such designs. In total for Round 2, 89 unique dCas9 designed variants (**Supplementary Table 2D**) were evaluated against NovoQ_r1 (**Supplementary Table 2E**). The Round 2 screening results are provided in **Supplementary Table 2F**.

In Round 3, the de novo C-terminal domain of NovoQ_r1 (residues 234-320) was redesigned using LigandMPNN against 10 dCas9 variants: 7 top-performing dCas9 variants from the Round 2 screen and 3 additional model-guided dCas9 variants selected using ridge regression analysis (see below). Each dCas9 variant was structurally predicted again with NovoQ_r1, gRNA, and target DNA using AlphaFold3(“Accurate Structure Prediction of Biomolecular Interactions with AlphaFold 3 | Nature,” n.d.) to verify the models. The NovoQ C-terminal domain was further redesigned using LigandMPNN. Resulting NovoQ sequences were clustered at 99% sequence identity, scored with Rosetta, and re-folded with Chai-1 using the same methods as above. Five hundred NovoQ sequences (**Supplementary Table 2G**) were selected to maximize the previously described scores and were screened against the ten dCas9 variants (**Supplementary Table 2H**). The Round 3 screening results are provided in **Supplementary Table 2I**.

### Regression modeling and generation of additional dCas9 variants

Following the Round 2 dCas9 experimental screen, activity and specificity scores (see below for score definitions) were used to train separate ridge regression models estimating the contribution of individual dCas9 interface substitutions to the activity and specificity scores observed in the screen. Variant dCas9 sequences were encoded as binary feature vectors by one-hot encoding amino acid identity at each position. Each feature represented the presence or absence of a specific amino acid substitution. Activity and specificity were modeled independently using ridge regression. For each model, sequences were split into 80% training and 20% validation sets. The ridge regularization parameter α was optimized with RidgeCV(Pedregosa et al. 2011). Model performance was evaluated on the withheld validation set and showed agreement between the predicted and experimental values (*R^2^* = 0.9586, MSE = 0.0420 for the specificity model; *R^2^* = 0.9430, MSE = 0.1036 for the activity model).

To generate additional dCas9 variants for Round 3, fitted model coefficients were used to identify substitutions predicted to maximize improvements to specificity without a loss of activity (**Figure S10**). The additional dCas9 variants are listed in **Supplementary Table 2D.**

### Molecular Cloning

Q5 Hot Start High-Fidelity DNA Polymerase (NEB, M0494L) was used for polymerase chain reaction (PCR). All primers were purchased from Integrated DNA Technologies (IDT), and gene fragments were purchased from Twist Bioscience. All newly cloned plasmids were constructed either by Golden Gate assembly using BsaI-HFv2 (NEB, R3733L), BsmBI-v2 (NEB, R0739L), and T4 DNA ligase (NEB, M0202L) or by Gibson assembly using NEBuilder® HiFi DNA Assembly Master Mix (NEB, E2621L).

The native PmcCAST pHelper and pDonor plasmids were gifts from Feng Zhang (Addgene #168153 and #168162)(Saito et al. 2021). T7 promoters and terminators in the PmcCAST pHelper were replaced with Lac promoters and rrnB terminators to generate pHelper_NovoCAST plasmids, which express two separate transcriptional cassettes: a Cas effector (Cascade or dCas9 variants) and a transposition machinery (TnsABC and TniQ variants). To construct a non-replicable pDonor, the R6K origin of replication was inserted into the PmcCAST_pDonor-Kan plasmid, which carries a kanamycin resistance gene as the DNA cargo. This non-replicable PmcCAST_pDonor-Kan plasmid was used for all transposition assays in *E. coli*. Amino acid sequences of proteins used in this study are provided in **Supplementary Table 3**. DNA sequences of all plasmids and gRNAs are provided in **Supplementary Table 4.**

### DNA Library Generation

Custom entry vectors or PCR-amplified vector backbones were used to assemble each unique DNA library. To construct entry vectors, the region of interest in pHelper_NovoCAST was replaced with BsaI or BsmBI Golden Gate flanking sites (e.g., the TniQ coding region was replaced with BsaI flanking sites in the TniQ design libraries). The same Golden Gate flanking sites were added to both ends of each variant in a library, together with a unique DNA barcode identifier. To accommodate long variant sequences, the full-length variant sequences were fragmented into oligos shorter than 300 bp, which included unique primer binding sites using the OMEGA program (Freschlin et al. 2025), and the resulting oligo pools were purchased from Twist Bioscience.

Oligo pools were resuspended in nuclease-free water to a final concentration of 1 ng/µL and PCR-amplified. The purified amplicons were inserted into an entry vector using Golden Gate assembly at a 20:1 insert-to-vector molar ratio. The assembly was cleaned up and electroporated into NEB 10-beta electrocompetent *E. coli* (NEB, C3020K). Following a 1 h recovery at 37 °C, cells were plated on agar plates with 100 μg/mL carbenicillin. To ensure library representation, colonies were collected at >100X coverage per variant, and the pHelper library was extracted using ZymoPURE™ II Plasmid Midiprep Kit (Zymo Research, D4201).

### High-Throughput Screening of NovoCAST Designs

To ensure that each transformant received only a single unique pHelper, plasmid libraries were electroporated into cJP003, an engineered *E. coli* strain harboring an arabinose-inducible *ccdB* gene (Park et al. 2025), at a 0.5:1 plasmid-to-cell ratio. Cells were recovered at 37 °C with agitation for 1 h and plated on agar plates containing 100 μg/mL carbenicillin. Colonies sufficient to achieve >100X coverage per variant were collected and cultured as biological duplicates at 37 °C. Inoculation was performed at a 2% ratio in LB medium supplemented with 100 μg/mL carbenicillin.

Cells were harvested at mid-log phase (OD_600_ ≈ 0.6), washed three times, and resuspended in ice-cold 10% glycerol to make them electrocompetent. As input reference samples for the library, pHelper plasmids were extracted from aliquots of each resuspended replicate using Monarch® Spin Plasmid Miniprep Kit (NEB, T1110L). PmcCAST pDonor-Kan was electroporated into the resuspended cJP003 cells at a 5:1 plasmid-to-cell ratio. Cells were immediately resuspended in 1 mL of pre-warmed SOC medium and incubated at 37 °C with vigorous shaking.

After 2 h, the incubated cells were plated on agar plates with 100 μg/mL carbenicillin and 50 μg/mL kanamycin for the activity screen, or agar plates with 100 μg/mL carbenicillin, 50 μg/mL kanamycin, and 0.1% (w/v) L-arabinose for the specificity screen. Selected colonies were scraped at >100X coverage per variant, and pHelper plasmids from each screen were purified using the ZymoPURE™ II Plasmid Midiprep Kit.

To prepare amplicons for next-generation sequencing (NGS), target regions of the extracted pHelper libraries and input reference samples were PCR-amplified, followed by index PCR using Nextera XT Index Kit v2 Set A (Illumina, FC-131-2001). Indexed amplicons were sequenced on an in-house MiSeq or on a NovaSeq X at the Hartwell Center at St. Jude Children’s Research Hospital. To determine the relative integration activity and specificity of each NovoCAST variant, NGS data were analyzed using a previously described Enrich2-based pipeline(Park et al. 2025; Rubin et al. 2017). Activity and specificity scores were calculated based on the log_2_-fold changes in variant frequencies relative to the input reference following the activity and specificity screens. DNA concentrations were measured using the Qubit™ 1X dsDNA HS Assay Kit (Thermo Fisher, Q33231), and all amplicons were purified using AMPure XP Beads (Beckman Coulter, A63881). Primer sequences used for NGS are listed in **Supplementary Table 5**.

### E. coli in vivo Transposition Assay

Each NovoCAST pHelper variant was cloned and introduced into the cJP003 strain. Transformed cells were plated individually on agar plates with 100 μg/mL carbenicillin. Single colonies for each variant were grown in LB medium supplemented with 100 μg/mL carbenicillin at 37 °C until mid-log phase (OD_600_ ≈ 0.6). These cultures were processed in triplicate using the same high-throughput screening protocol. Integration activity and specificity were assessed by counting the number of colonies on agar plates with 100 μg/mL carbenicillin and 50 μg/mL kanamycin, with or without 0.1% (w/v) L-arabinose.

### Genome-wide Integration Profiling

Integration profiles of individual NovoCAST variants were characterized using tagmentation-based transposon insertion sequencing (TagTn-seq) (George et al. 2023). Following the *E. coli in vivo* transposition assay for each NovoCAST variant, over 500 colonies per variant were harvested from agar plates with 100 μg/mL carbenicillin and 50 μg/mL kanamycin, and genomic DNA (gDNA) was extracted using Wizard® Genomic DNA Purification Kit (Promega, A1125).

The extracted gDNA was tagmented using Nextera XT DNA Library Kit (Illumina, FC-131-1024) and the resulting fragments were PCR-amplified using PmcCAST left end (LE)-specific primers and an i7 primer (**Supplementary Table 5**). These amplicons were indexed with the Nextera XT Index Kit v2 Set A and sequenced on the MiSeq instrument.

### Analysis of Genome-wide integration profiling

Analysis of TagTn-seq data was performed using previously published methods (George et al. 2023) and custom Python scripts. Raw sequencing reads were merged and filtered for read quality using a minimum Phred threshold of 30. Reads containing the last 22 bp of the LE sequence (5’-CAAAGTTATGGGTAAAGTCACA-3’) were identified, and the adjacent 22 bp genomic sequences were extracted as transposon flanking reads. Flanking reads containing the intact PmcCAST pDonor-Kan sequence (5’-ATCCCAATGGCGCGCCGAGCTT-3’) were filtered out. The remaining flanking reads were then mapped back to the reference *E. coli* strain cJP003 genome using Bowtie2 (Langmead and Salzberg 2012). Only reads that mapped uniquely without any mismatches to the genome were used to generate the genome-wide integration profiles. Read position indexing with respect to the target site duplication and the orientation of the transposon insertion (t-LR vs. t-RL) was performed as described previously (George et al. 2023).

On-target reads were defined as flanking reads that mapped within a 100 bp window downstream of the PAM, whereas reads that mapped elsewhere in the genome were classified as off-target. Integration distances (d) were calculated between the insertion position and the target site PAM for on-target reads. Raw reads were normalized for each sample to visualize the genome-wide integration events.

For analysis of the transposition sequence preferences, only unique insertion sites were used to minimize amplification biases during sequencing. For each insertion site in the reference genome, a 400 bp sequence window was extracted (200 bp upstream and downstream). These sequence windows were aligned with respect to their insertion site at position 0 and then were used to generate position-specific frequency matrices to represent the frequency of each nucleotide. Sequence logos were plotted using LogoMaker (Tareen and Kinney 2020) to check for transposition sequence preferences and PAM enrichment across off-target integration events.

### Off-target candidate guide generation

Guide candidates were identified for off-target hot-spots with enriched integration in genome-wide integration profiling. Mapped insertion coordinates from tagmentation sequencing were filtered to retain reads with mapping quality scores ≥ 30. Insertion positions were counted and clusters were collapsed into single sites by merging sites within 5 bp, with each cluster represented by its most abundant insertion coordinate. Insertion sites with total read counts > 1000 were retained for downstream analysis.

Candidate protospacers were identified for each retained insertion site using three approaches. First, genomic sequences within ±120 bp of each integration site were scanned on both the forward and reverse strands to identify canonical PAM sites, and the corresponding protospacer sequences were extracted as the 20 bp immediately upstream of the PAM. Second, the original guide sequence was locally aligned to the insertion site-centered genomic window, and the best-scoring alignment was used to define a homologous genomic region from which the candidate protospacer was derived. Third, additional protospacers were identified by scanning a 50 bp window centered on the best locally aligned region for PAMs, followed by extraction of the corresponding 20 bp protospacers.

For downstream analysis, candidate protospacer sequences were restricted to those located 66-76 bp from an integration site with an associated PAM. Candidates were further filtered to retain those for which the PAM strand was opposite to the dominant insertion strand of a site, as determined from strand-resolved insertion counts. The resulting filtered candidate guides were used for visualization of guides targeting selected integration sites, including the *fliT* locus.

### Protein expression and purification

Proteins were encoded under the *lacO*-controlled T7 promoter and tagged with an N-terminal 6XHis-Strep-SUMO tag, with a ULP1 protease recognition site positioned immediately before the gene of interest to allow subsequent tag removal. The expression vectors (**Supplementary Table 4**) were introduced into One Shot BL21(DE3) chemically competent *E. coli* (Invitrogen, C600003), and the cells were cultured in 2XYT medium at 37 °C with shaking until OD_600_ reached 0.4-0.6. Protein overexpression was then induced by adding 0.25 mM isopropyl-β-D-thiogalactopyranoside (IPTG), followed by overnight incubation at 16 °C.

Cells were harvested and resuspended in His-lysis buffer (50 mM HEPES pH 7.5, 500 mM NaCl, 5% glycerol, and 1 mM DTT) supplemented with a protease inhibitor cocktail tablet (Thermo Scientific, A32963). Cells were then lysed by sonication on ice using 7 cycles of 5 s on and 10 s off pulses. The soluble fraction was collected by centrifugation at 16,000 rpm for 30 min at 4 °C and applied to Ni-NTA resin (Thermo Scientific, 88221) pre-equilibrated with His-lysis buffer. The resin was washed with 10 column volumes of His-wash buffer (50 mM HEPES pH 7.5, 500 mM NaCl, 5% glycerol, 50 mM imidazole, and 1 mM DTT). Target proteins were then eluted using His-elution buffer (50 mM HEPES pH 7.5, 500 mM NaCl, 5% glycerol, 1 mM DTT, and 500 mM imidazole). The eluted proteins were subjected to overnight digestion at 4 °C with in-house ULP1 protease to remove the tags.

Tag-cleaved fractions were pooled and diluted with heparin dilution buffer (50 mM Tris-HCl pH 7.5, 50 mM NaCl, 5% glycerol, and 1 mM DTT) to a final NaCl concentration of 200 mM. The diluted sample was loaded onto a HiTrap Heparin HP column (Cytiva, 17040601) pre-equilibrated with heparin low-salt buffer (50 mM Tris-HCl pH 7.5, 200 mM NaCl, 5% glycerol, and 1 mM DTT). Proteins were eluted using a linear salt gradient (200 mM to 1.1 M NaCl), generated by mixing the low-salt buffer with heparin high-salt buffer (50 mM Tris-HCl pH 7.5, 1.1 M NaCl, 5% glycerol, and 1 mM DTT). Elution was monitored by UV absorbance at 280 nm, and peak fractions were analyzed by SDS-PAGE. Pooled heparin fractions were concentrated and resolved by size-exclusion chromatography on a Superdex 200 10/300 GL column (Cytiva, 17517501) mounted on an ÄKTA Pure FPLC system and pre-equilibrated with gel filtration buffer (50 mM Tris-HCl pH 7.5, 500 mM NaCl, 5% glycerol, and 1 mM DTT). The peaks corresponding to the expected molecular weights of the target proteins were collected. Final protein purity was confirmed by SDS-PAGE. All proteins were purified using the same protocol unless stated otherwise.

PmcTnsC displayed multiple distinct peaks in the elution profile, indicating various oligomeric states, and only fractions corresponding to the monomeric form of PmcTnsC were pooled and concentrated. NovoQ_r3 was co-purified with untagged PmcTnsC, and fractions containing both proteins were collected.

### Preparation of DNA substrates

DNA substrates were designed to incorporate binding sites for dCas9_r2, NovoQ_r3 and TnsC based on integration profiles and were assembled using three oligonucleotide strands (**Supplementary Table 6**). The partial non-target strand is complementary to the complete target strand, and the desthiobiotinylated LUEGO strand is complementary to the 3’ region of the target strand. The three oligonucleotides were annealed at equimolar ratios in annealing buffer (10 mM Tris pH 7.5, 50 mM NaCl, and 1 mM EDTA). The mixture was denatured at 95 °C for 5 min and then cooled to 50 °C at a rate of 1 °C/min using a thermal cycler. All oligonucleotides were purified by high-performance liquid chromatography (HPLC) using an Agilent 1260 Infinity II system equipped with an AdvanceBio Oligonucleotide column (Agilent Technologies).

### Reconstitution of the recruitment complex and the DNA-pull-down

To reconstitute the recruitment complex, the annealed DNA substrate was mixed with *ccdB*_01 gRNA, dCas9_r2, PmcTnsC monomers, and the NovoQ_r3-PmcTnsC complex at a molar ratio of 1:1:1:0.5:1 in a final reaction volume of 200 μL. Reconstitution was carried out in reconstitution buffer (26 mM HEPES pH 7.5, 20 mM KCl, 2 mM MgCl_2_, 2% glycerol, 2 mM ATP, and 1 mM DTT), followed by incubation at 37 °C for 30 min.

DNA-bound complexes were isolated using a streptavidin-affinity pulldown. Streptavidin magnetic beads (Cytiva, 28985799) were washed three times with 500 μL of reconstitution buffer and incubated with the reconstituted sample for 1 h at room temperature under gentle rotation. Beads were washed three times with 500 μL of reconstitution buffer to remove unbound proteins.

Bound complexes were eluted in two sequential steps using 50 μL of elution buffer (reconstitution buffer supplemented with 10 mM biotin) per step, with a 30 min incubation on a rotator for each elution. Eluted fractions were analyzed by SDS–PAGE to confirm the presence and integrity of all protein components.

### Cryo-EM sample preparation and freezing of the recruitment complex

Graphene-coated grids were prepared according to the protocol developed previously (Ahn et al. 2023) with minor modifications. Briefly, a graphene sheet was allowed to settle onto the grids and was left overnight at room temperature to anneal gradually to the grids. The next day, the grids were baked at 150 °C for 30 min for firm annealing. To remove polymethyl methacrylate (PMMA), three rounds of acetone washes were performed, and the quality of graphene layering was verified using a Talos (Thermo Fisher Scientific).

The 1:1-diluted recruitment complex sample was applied on the carbon side of a graphene-coated grid and incubated for 30 s in the Mark IV Vitrobot chamber (Thermo Fisher), which was set to 4 °C and 100% humidity. Each grid was blotted for 3.5 s with a blot force of −7 and then plunged into liquid ethane cooled by liquid nitrogen.

### Cryo-EM imaging and image processing of transpososome complex

Vitrified samples of the recruitment complex were first screened using a Talos Arctica (Thermo Fisher) operated at 200 kV prior to large-scale data collection. Promising screened grids were imaged using a Titan Krios (300 kV, Thermo Fisher) equipped with a BioQuantum energy filter (Gatan) and a Falcon 4i direct electron detector (4k x 4k). Initially, 11,377 micrographs were collected using EPU (3.13.0.10277) at 160,000× magnification (0.724 Å per pixel) using image shift, with a nominal defocus range of −1.5 μm to −2.5 μm. Each movie was collected with a 1 s exposure with a total dose of 60 electrons per Å², fractionated into 60 frames. Frames were aligned using MotionCor2, and contrast transfer function (CTF) estimation was performed in cryoSPARC v4.7.1 for downstream image analysis. The workflow described below is shown in **Figure S14**.

Using the blob picker, 770,000 particles were initially picked, and the stack was subjected to 2D classification. Two-dimensional classification resulted in 200 classes, of which a few appeared to represent the recruitment complex. After selecting those classes, 6,000 particles were obtained and used to train Topaz. After obtaining a well-trained model, additional particles were picked and extracted from the micrographs. The new set of particles was also subjected to 2D classification, and the best classes were selected. These particles were used to further train the topaz model, and particles were extracted from the micrographs. The best-looking 2D classes were subjected to ab initio reconstruction. Ab initio reconstruction separated the particles into two classes, one containing 9,679 particles (free Cas9) and another containing 25,527 particles (the recruitment complex). The two classes were then subjected to non-uniform refinement separately. A similar pipeline was followed for a follow-up data collection with 13,405 micrographs. This dataset resulted in 5,951 particles corresponding to the recruitment complex. Particles from both datasets were merged, and heterogeneous refinement followed by non-uniform refinement was performed to remove junk particles. This eliminated 4,370 particles and produced a map of 3.29 Å resolution for the recruitment complex composed of 27,108 particles. Local refinement was performed to enhance the structural details, resulting in a final 3.2 Å map composed of 27,108 particles.

After merging the datasets, 2D classes with promising density for the TnsC ring were selected and processed separately. A total of 4,111 particles were subjected to ab initio reconstruction, non-uniform refinement, and local refinement, resulting in a map of 7.3 Å resolution in which three TnsC subunits could be observed.

Model building was performed using an AlphaFold3(“Accurate Structure Prediction of Biomolecular Interactions with AlphaFold 3 | Nature,” n.d.)-predicted model of NovoQ_r3, dCas9_r2, and TnsC with target DNA. The model was refined into the high-resolution density map using Rosetta FastRelax(Wang et al. 2016; Alford et al. 2017). Refinement was carried out in Phenix (v1.13). Model assessment was performed in Coot(Emsley et al. 2010). The final model was analyzed and visualized using ChimeraX (v1.11.1) and PyMOL (v3.1.7.2).

### Cointegrate analysis

*In vivo* transposition assays were performed in *E. coli* using NovoCAST_r4 and the *ccdB*_01 gRNA. Following Kan+Ara selection, more than 1,000 surviving colonies representing enriched on-target integrations were pooled by scraped for gDNA extraction using Monarch® Spin gDNA Extraction Kit (NEB, T3010L). The extracted gDNA was sent to Plasmidsaurus for whole-genome sequencing. The raw FASTQ files were analyzed to identify simple insertions and cointegrates. Raw reads containing the target site (TTTCGCGGTGGCTGAGATCAGCCACTTCTTCCCCGATAACGGAGACCGGCACACTGGCC) followed downstream by the PmcCAST right end (RE) sequence were identified as integration-containing reads. Among these reads, those containing the pDonor backbone sequence (CCTGATGCGGTATTTTCTCCTTACG) were classified as cointegrates. Reads lacking the pDonor backbone sequence were classified as simple insertions, or as ambiguous if the RE-downstream sequence was shorter than 200 bp.

### Computational stabilization of TnsAB

A strand transfer complex (STC) model of PmcCAST TnsAB tetramer bound to target and donor DNA was generated using AlphaFold3(“Accurate Structure Prediction of Biomolecular Interactions with AlphaFold 3 | Nature,” n.d.) and used as the starting structure for computational stabilization via structure-based sequence design. To preserve residues expected to be important for catalysis, DNA sequence specificity, and functional integrity, several classes of residues were excluded from redesign. Fixed residues included: 1. a predicted disordered loop spanning residues 856-898, 2. residues within 6 Å of bound DNA, 3. residues within 7 Å of either catalytic site (residues E62, D104, and K119, corresponding to the catalytic site of the TnsA region; D519, D588, and E624, corresponding to the catalytic site of the TnsB region), and 4. highly conserved residues identified with Consurf(Ashkenazy et al. 2016) with conservation grades of ≥ 8. These constraints yielded a total of 384 redesignable positions across the TnsAB STC model. As an additional stabilization strategy, redesign was further restricted to positions predicted by ThermoMPNN(Dieckhaus et al. 2024) to be destabilizing in each TnsAB conformation. The TnsAB STC tetramer contains two TnsAB protomers in an active conformation and two protomers in an inactive, stabilizing conformation. This additional design strategy reduced the number of redesignable positions to 43 residues.

TnsAB sequence variants were generated using LigandMPNN with per-residue sampling biases applied to the output logits prior to Softmax normalization(Dauparas et al. 2025). Bias values were assigned to favor either the wild-type amino acid or amino acids conserved across TnsAB homologs identified from sequence alignments. Designed variants were subsequently folded as full STC assemblies using AlphaFold3(“Accurate Structure Prediction of Biomolecular Interactions with AlphaFold 3 | Nature,” n.d.). Designed variants were prioritized for selection based on minimal mutation count with the following explicit filters: complex ipTM > 0.7, predicted clash score < 0.5, per-chain RMSD relative to the active conformation < 0.85 Å, and full STC RMSD < 5 Å. Six TnsAB designs satisfying these criteria and spanning 13-36 mutations were selected for experimental testing in human cells (**Supplementary Table 2J**).

### Mammalian cell culture

HEK293FT cells (Invitrogen, R70007) were cultured in Dulbecco’s Modified Eagle Medium (Gibco, 10566024) supplemented with 10% (v/v) fetal bovine serum (Corning, 35-010-CV), 1% (v/v) antibiotic-antimycotic (Gibco, 15240062), 1% (v/v) MEM non-essential amino acids (Gibco, 11140050), and 1 mM sodium pyruvate (Gibco, 11360070). Cells were incubated at 37 °C in a humidified incubator with 5 % of CO_2_.

### Episomal and Genomic Integration assays in human cells

For the episomal integration assay in human cells, 900,000 HEK293FT cells per well were co-transfected with 450 ng of pTnsABC, 450 ng of pCasQ, 850 ng of pTarget-*ccdB*, and 850 ng of pDonor using Lipofectamine 3000 transfection reagent (Invitrogen, L3000008) in 6-well plates. After 72 h of incubation, cells were harvested and washed three times with ice-cold phosphate-buffered saline (PBS). The plasmids were isolated from the cells using the Monarch® Spin Plasmid Miniprep Kit. The recovered plasmids were then submitted to Plasmidsaurus for sequencing, and targeted integration in pTarget was further validated by junction PCR using a LE-specific primer and a target site-specific primer.

For genomic integration assay in human cells, 900,000 HEK293FT cells per well were co-transfected with 450 ng of pTnsABC, 450 ng of pCasQ, and 1,350 ng of pDonor in 6-well plates. To minimize false-positive signals, 250 ng of pDonor carrying a puromycin resistance gene under the hPGK promoter was also co-transfected. At 24 h post-transfection, the cells were subjected to selection with 1 μg/ml puromycin (InvivoGen, ant-pr-1) for 72 h. Human gDNA was subsequently extracted using the Monarch® Spin gDNA Extraction Kit.

### Western Blot

HEK293FT cells were transfected with 2.5 μg of mammalian expression vectors encoding NovoCAST_r3 components either individually or in combination. All proteins were tagged with 3xFLAG tags. Cells were harvested 48 h post-transfection, washed twice with PBS, and lysed using M-PER™ mammalian protein extraction reagent (Thermo Fisher, 78503) supplemented with a protease inhibitor cocktail (Thermo Fisher, 87785). The soluble fraction of the lysates was collected by centrifugation, and protein concentrations were determined using a Pierce™ Dilution-Free™ rapid gold BCA protein assay (Thermo Fisher, A55862). Lysates were normalized to the same protein concentration and denatured in Laemmli sample buffer (Bio-Rad, 1610747) supplemented with DTT at 70 °C for 10 min. Proteins were separated by SDS-PAGE using 4–20% precast gels (Bio-Rad, 4561093) and transferred to nitrocellulose membranes (Thermo Fisher, IB33002). Membranes were blocked with Pierce™ clear milk blocking buffer (Thermo Fisher, 37587) and incubated with anti-FLAG antibodies (Sigma-Aldrich, F3165; 1:5,000 dilution) overnight at 4 °C. Membranes were then washed three times with TBS Tween 20 Buffer (Thermo Fisher, 28360) and incubated with HRP-conjugated secondary antibody (Invitrogen, A28177) in the wash buffer (1:10,000 dilution) at room temperature for 1 h. Following three washes with the wash buffer, protein bands were visualized using clarity western ECL substrate (Bio-Rad, 1705060) and imaged with a ChemiDoc imaging system (Bio-Rad). A loading control, β-actin, was detected using anti-β-actin antibodies (Invitrogen, MA1-140) after stripping the same membranes using stripping buffer (Thermo Fisher, 21059).

### Droplet digital PCR (ddPCR)

To quantify the on-target integration efficiency of NovoCAST in human cells, duplex ddPCR was performed on plasmids or gDNA isolated from integration assays in human cells. Primer and probe sets were designed to target a reference site (the pTarget backbone for the episomal integration assays, or *RPP30* for the genomic integration assays) and the integration junction (**Supplementary Table 5**).

ddPCR reactions were prepared using ddPCR supermix for probes (No dUTP) (Bio-Rad, 1863023). Each primer-probe set was added to the reaction mixture (20 µL total volume) for each sample, resulting in final concentrations of 900 nM for each primer and 250 nM for each probe. Either 0.1 pg of plasmid template or 20 ng of gDNA template was used as templates per reaction. To improve template accessibility, in-well restriction digestion was performed by adding 5 U of EcoRI-HF (NEB, R3101S) for plasmid templates and 5 U of NcoI-HF (NEB, R3193S) for gDNA templates directly to the reaction mixtures.

Droplets were generated using droplet generation oil (Bio-Rad, 1863005) according to the manufacturer’s instructions, sealed, and subjected to PCR amplification under the following conditions: 25 °C for 3 min; 95 °C for 10 min; 40 cycles of 94 °C for 30 s and 60 °C (genomic) or 62.5 °C (episomal) for 1 min; 98 °C for 10 min; followed by a 4 °C hold. A ramp rate of 2 °C/s was used for all steps. Droplets were read on a QX200 Droplet Reader (Bio-Rad), and data were analyzed using QX Manager software version 2.4 to quantify integration events. On-target integration frequency (%) was calculated as the ratio of the copy number of the integration junction detected by a FAM-labeled probe to that of the reference site detected by a HEX-labeled probe (fractional abundance, **Supplementary Table 7**).

---

**Figure S1.**
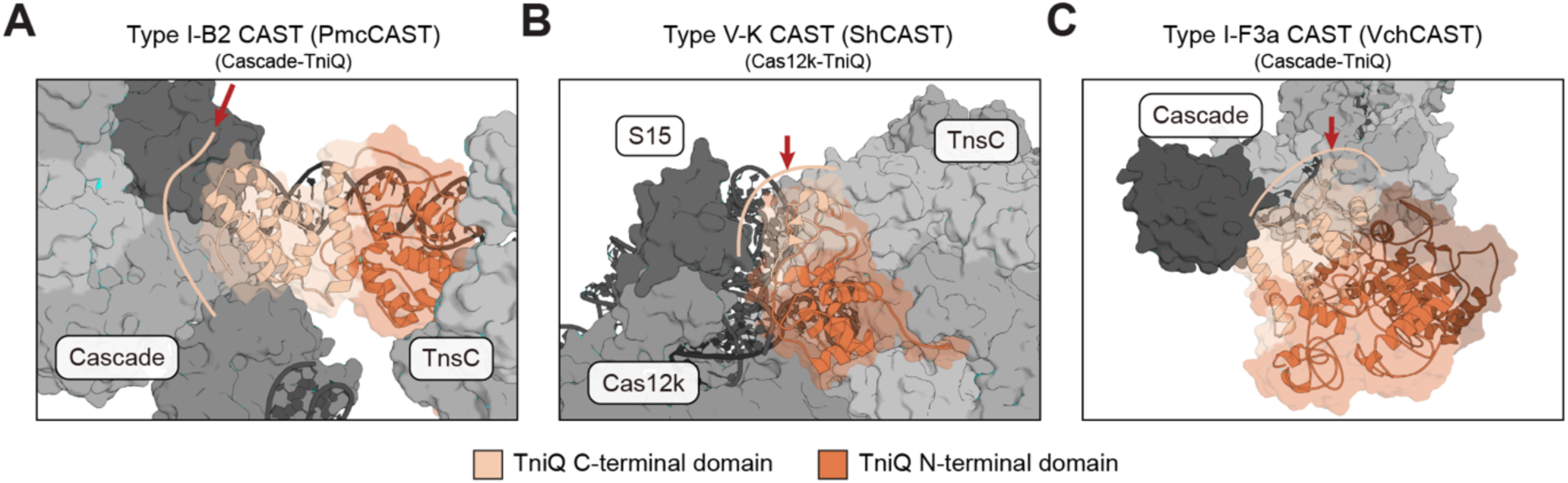
TniQ C-terminal domain interactions with diverse CRISPR effectors. Structures of representative CAST types shown at the TniQ-CRISPR effector interface. Structures are shown with colors that are consistent with those used in Figure 1B: TniQ is shown in orange (N-terminal domain) and beige (C-terminal domain) cartoons. Other components (gray, surface representation) are labeled. The red arrow and beige line indicate the location of the CRISPR effector-TniQ interface. **A.** Type I-B2 CAST (PmcCAST) TniQ-Cascade interface (PDB: 8FF4). Cascade is the CRISPR effector. **B**. Type V-K CAST (ShCAST) TniQ-Cas12k interface (PDB: 8EA3). S15 is a bacterial host factor that associates with the transposon-encoded CRISPR effector, Cas12k **C.** Type I-F3a CAST (VchCAST) TniQ-Cascade interface (PDB: 6PIJ). Cascade is the CRISPR effector.

**Figure S2.**
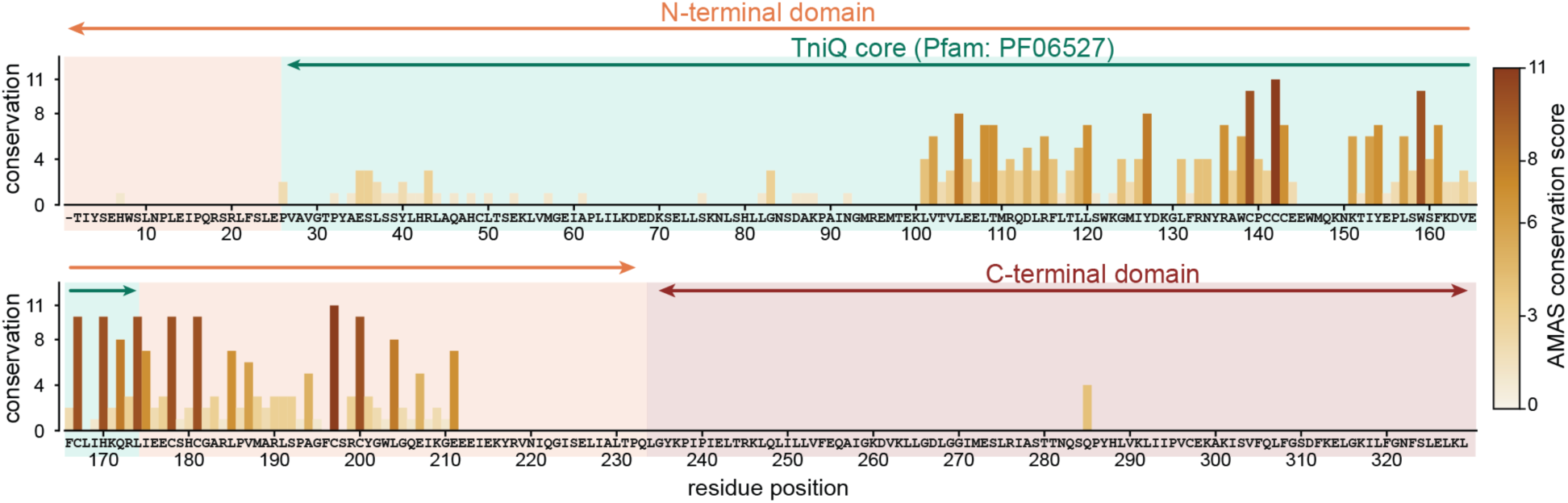
Conservation plot of CAST TniQ homologs across subtypes reveals conserved and variable regions, which are associated with different functional domains. Alignment-based mutational analysis (AMAS) conservation scores are shown on the y-axis in histogram format for each position in the multiple sequence alignment (MSA), with higher values indicating greater conservation. Colors correspond to the level of conservation, as indicated in the legend on the right. TniQ (Pfam: PF06527), corresponding to the Pfam HMM profile, is shaded green, with its range indicated by green arrows at the top of the plot. The N-terminal domain, which was held fixed during design, is shaded orange and indicated by orange arrows. The C-terminal domain is shaded red and indicated by red arrows.

**Figure S3.**
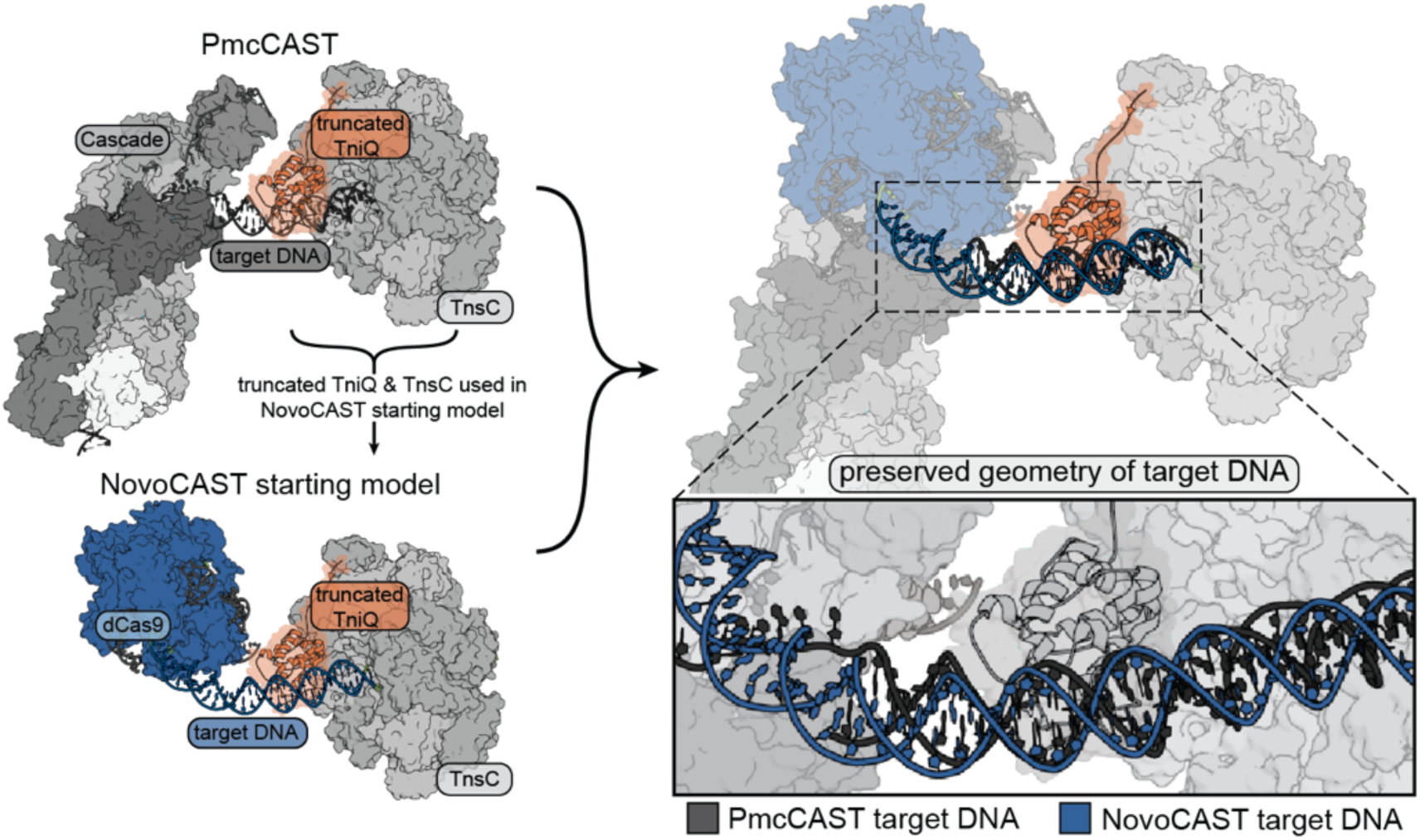
Construction of the starting point for design. Top left: Type I-B2 PmcCAST is shown (PDB: 8FF4) with truncated TniQ (orange). The color scheme and structural representation match those shown in Figure 2A-B. Bottom left: dCas9 (blue; PDB: 7OXA) is positioned by aligning its target DNA with that of PmcCAST, ensuring that NovoCAST recapitulates the native target DNA geometry required for transposition. Right: structural alignment of PmcCAST and the NovoCAST starting model. Target DNA (black/blue) shown in inset for clarity.

**Figure S4.**
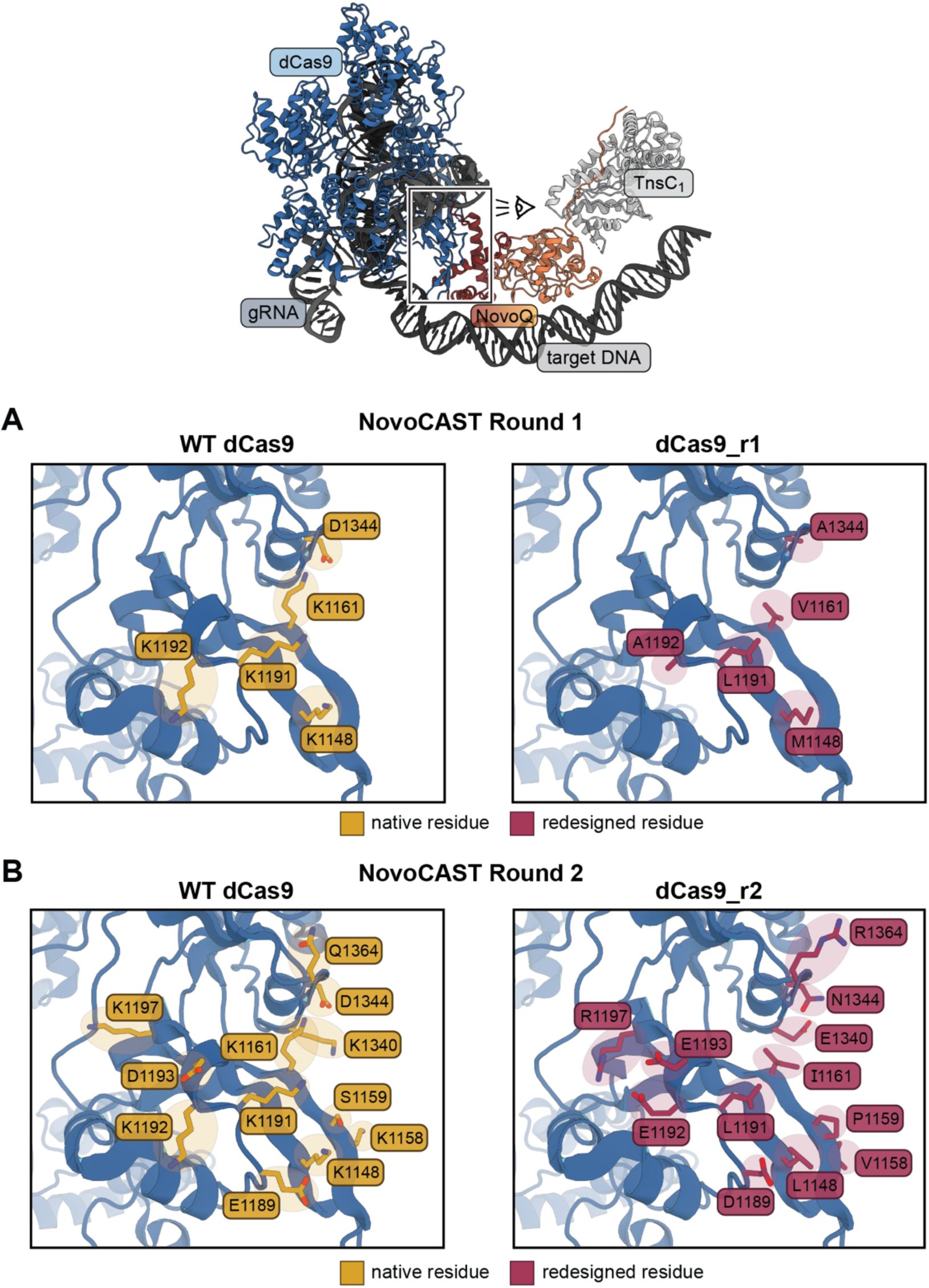
dCas9 positions mutated in Rounds 1 and 2 of NovoCAST designs. Top: overview of NovoCAST model. The boxed region indicates the designed dCas9-NovoQ interface that is highlighted in panels **A & B**. The eye symbol indicates the viewing direction. **A.** Round 1 of NovoCAST designs allowed for five positions on the WT-dCas9 surface (blue cartoon, PDB: 7OXA) to vary during design: K1148, K1161, K1191, K1192, and D1344 (yellow residues, left panel). Positions allowed to mutate are shown; sidechains are displayed in stick representation. The variant dCas9_r1 contains five mutations that create a more favorable binding surface for NovoQ recognition: K1148M, K1161V, K1191L, K1192A, and D1344A (magenta residues, right panel). **B**. Round 2 of NovoCAST designs, which redesigned dCas9_r1 residues that were within 4 Å of NovoQ_r1, led to the selection of dCas9_r2 which contains twelve surface mutations relative to WT-dCas9: K1148L, K1158V, S1159P, K1161I, E1189D, K1191L, K1192E, D1193E, K1197R, K1340E, D1344N, and Q1364R (WT-dCas9: left panel; dCas9_r2: right panel).

**Figure S5.**
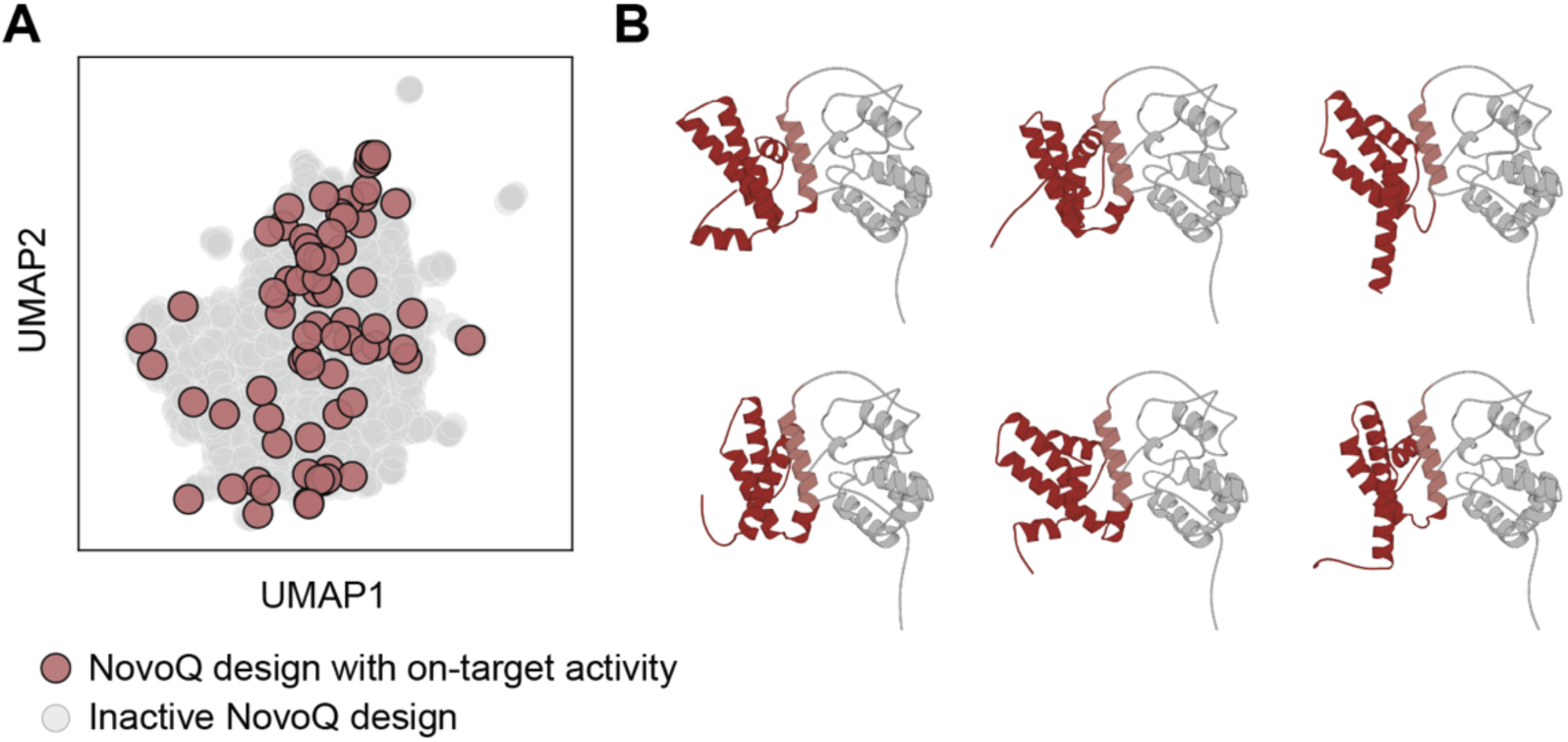
Sequence and structural diversity of NovoCAST Round 1 designs. **A.** UMAP of NovoQ ESM-2 sequence embeddings. Gray points represent all NovoQ sequences included in the initial screen. Light red points represent the 76 NovoQ sequences that resulted in quantifiable on-target integration. **B.** Representative subset of NovoQ folds predicted with Chai-1. The gray region represents the N-terminal domain. Red represents the *de novo* designed C-terminal domain. All NovoQ designs are shown aligned to the N-terminal domain.

**Figure S6.**
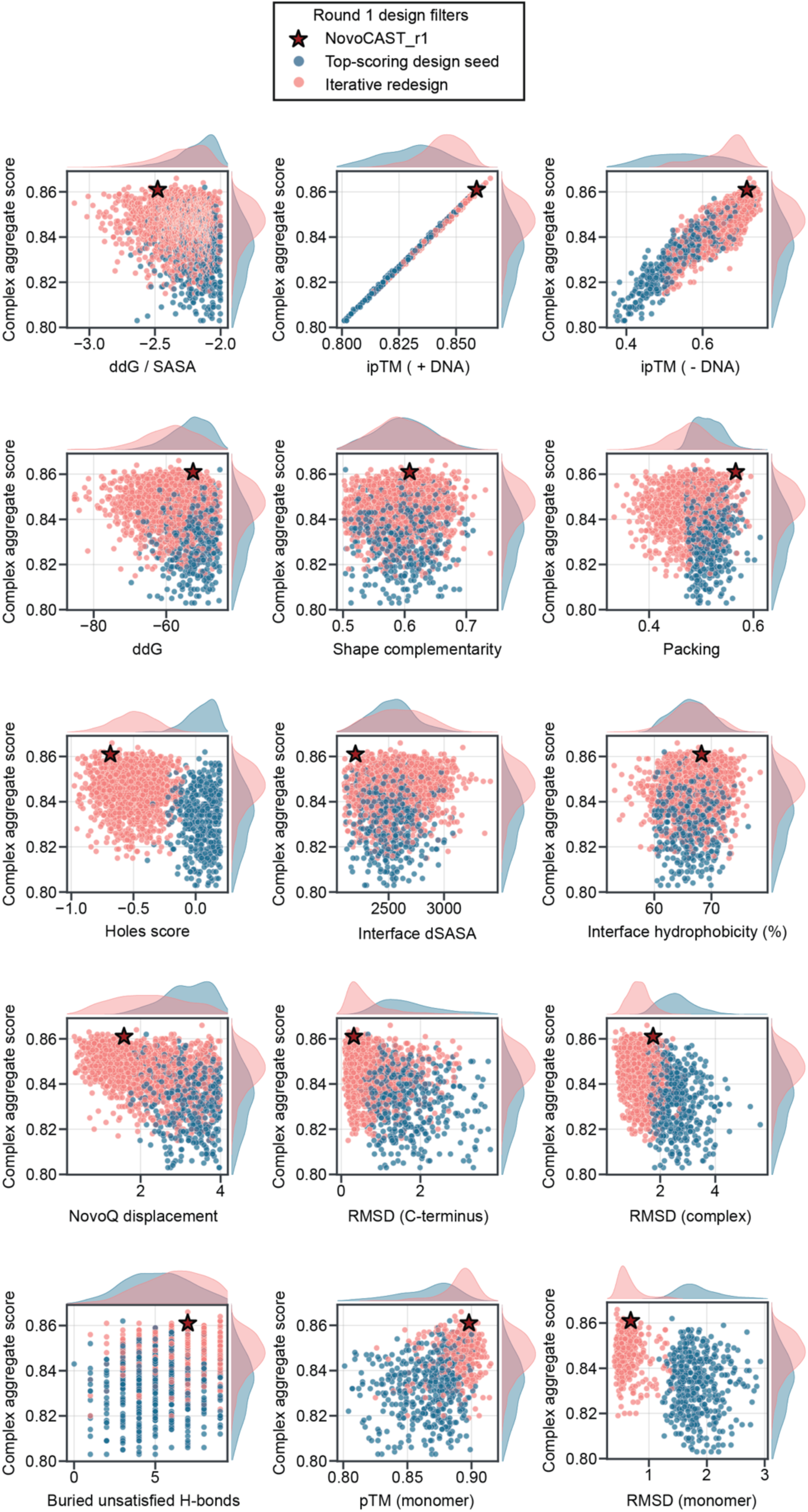
Distribution of design scores for NovoQ sequences that were prioritized for experimental screening. Complex aggregate score (y-axis) is the score output from Chai-1 and was used as a reference for the individual scoring filters. Blue population corresponds to top scoring seed designs; pink population corresponds to the designs selected after iterative redesigning until convergence. Kernel Density Estimates (KDEs) are displayed on the top and right axes to show the distribution of scores. NovoCAST_r1 score in each plot is indicated by a red star. Details of the scoring metrics can be found in the Methods and Supplementary Materials.

**Figure S7.**
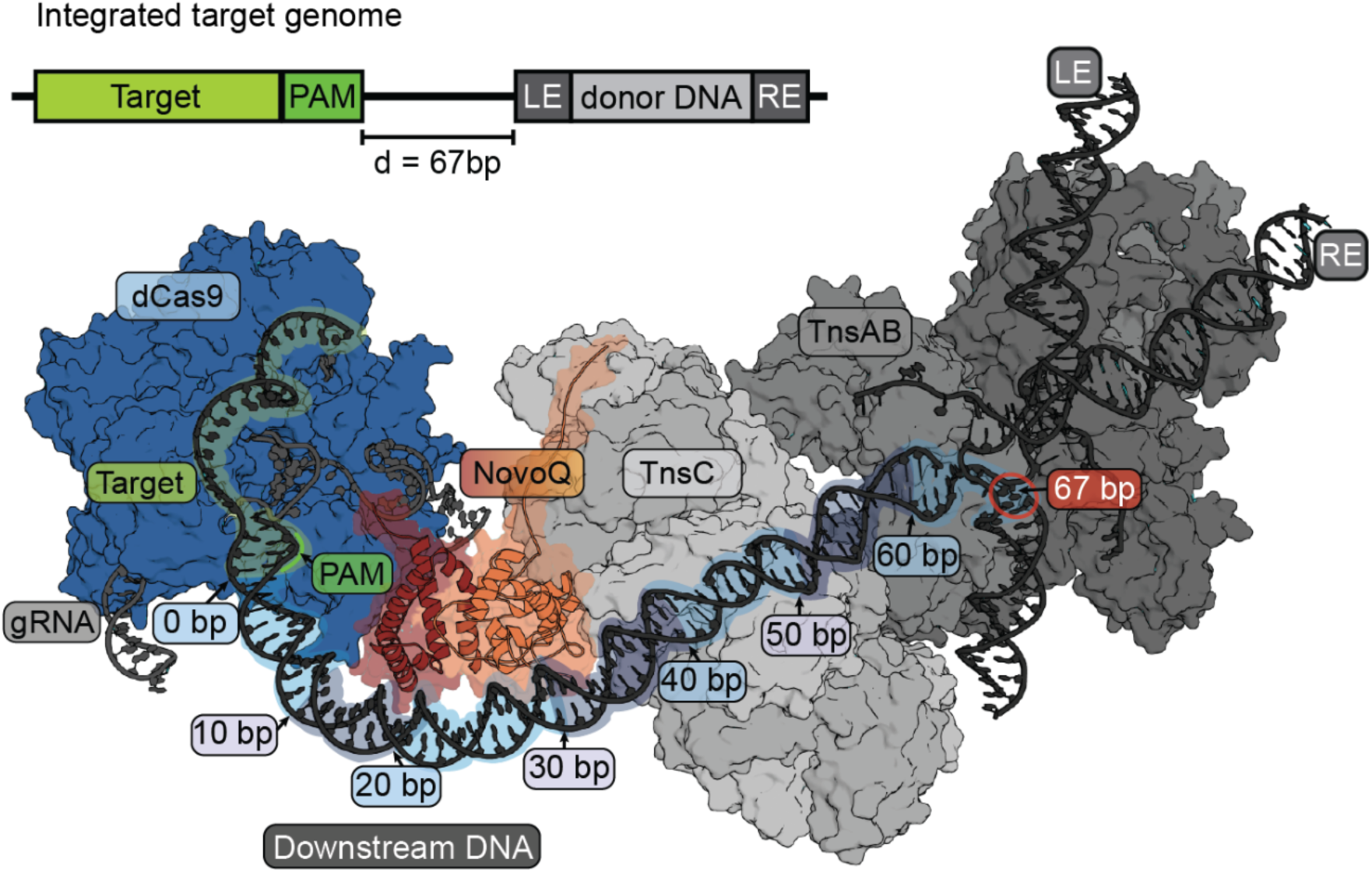
Expected NovoCAST integration complex architecture. Expected integration product (top), as defined by the modeled integration complex architecture (bottom). Target for gDNA is colored light green, PAM motif shown in darker shade of green. LE, transposon left end; RE, transposon right end; d, distance from the PAM to the integration site (bp). Colors follow the convention established in Figure 2. The transposition machinery was modeled using the cryo-EM structure of PmcCAST TnsABC (PDB: 9BW1) aligned onto an AlphaFold3 predicted model of the NovoQ_r3 – dCas9_r2 complex. Target DNA is labeled every 10 bp, starting from the end of the PAM motif. The red circle at 67 bp on the right indicates the predicted integration site.

**Figure S8.**
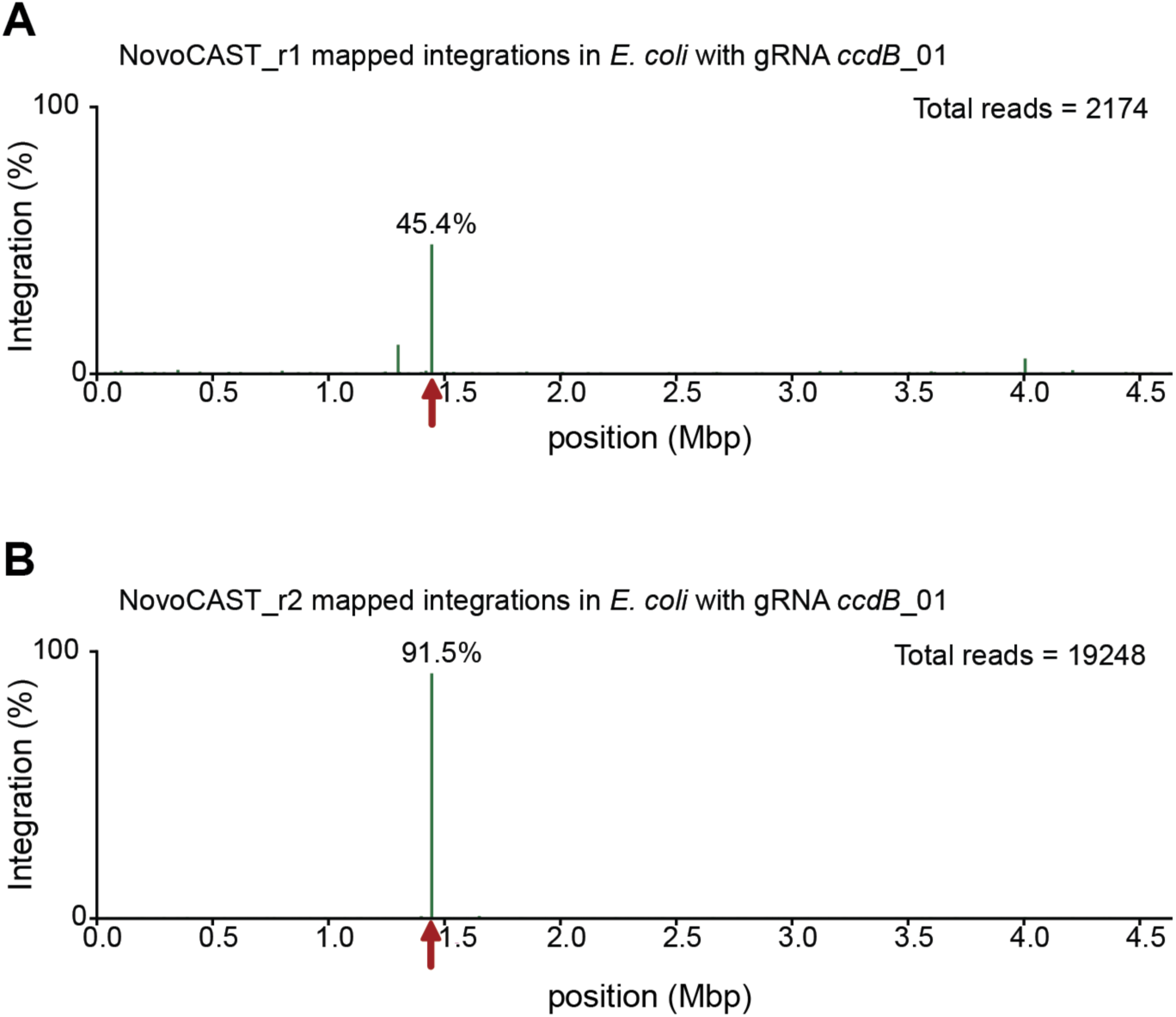
Genome-wide integration profile of NovoCAST Rounds 1 and 2 in *E. coli* targeting *ccdB.* **A.** Genome-wide distribution of mapped NovoCAST_r1 integration events in *E. coli* using *ccdB*_01 gRNA. The targeted genomic position is highlighted with a red arrow, with 45.4% of insertions occurring on target. **B.** Genome-wide distribution of mapped NovoCAST_r2 integration events in *E. coli* using the same gRNA. The targeted genomic position is highlighted with a red arrow, with 91.5% of insertions occurring at the target site.

**Figure S9.**
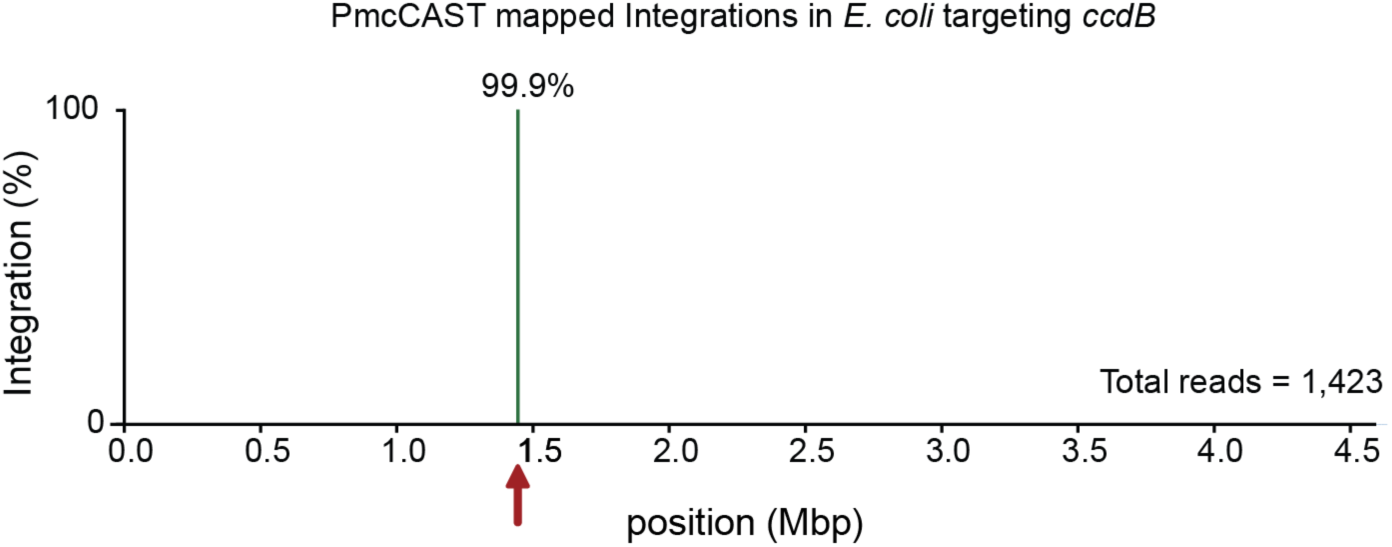
Genome-wide integration profile of PmcCAST in *E. coli* targeting *ccdB.* Genome-wide distribution of mapped PmcCAST integration events in *E. coli* using the best-performing Pmc_*ccdB* gRNA (Supplementary Table S4). The targeted genomic position is indicated with the red arrow.

**Figure S10.**
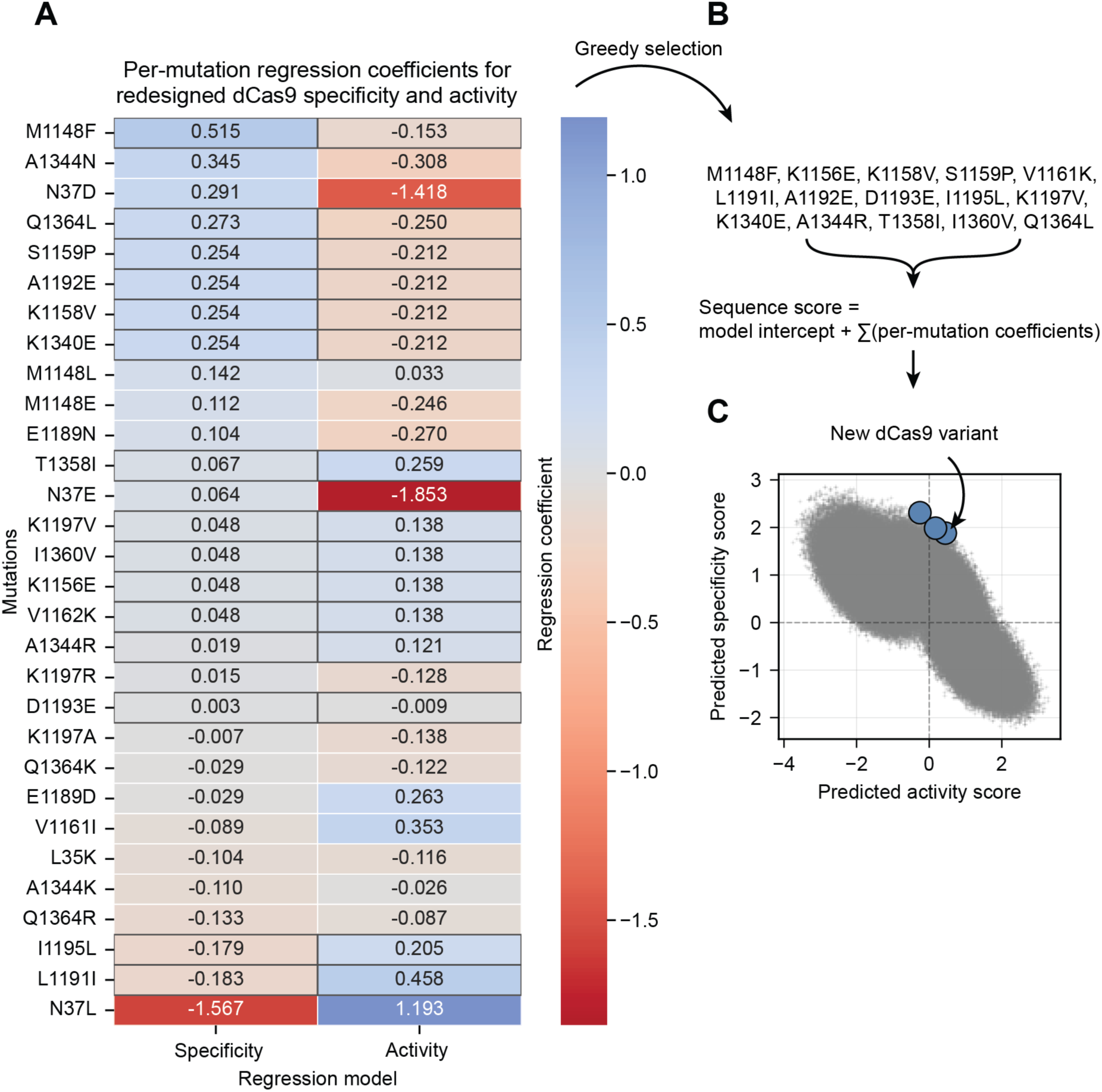
Ridge regression-guided design of dCas9 variants with improved specificity. **A.** Heatmap of per-mutation regression coefficients from two ridge regression models trained on Round 2 experimental data (see Methods). Rows are individual dCas9 mutations from Round 2, and columns correspond to specificity and activity models, colored by the magnitude of the regression coefficient (red to blue). **B.** Greedy selection of mutations using the specificity coefficients in panel A. The predicted score is the model intercept and sum of per-mutation coefficients. **C.** The predicted specificity (y-axis) and activity (x-axis) scores for sampled dCas9 variants are shown in gray and the three regression dCas9 variants that maximize specificity scores without loss of activity are highlighted in blue.

**Figure S11.**
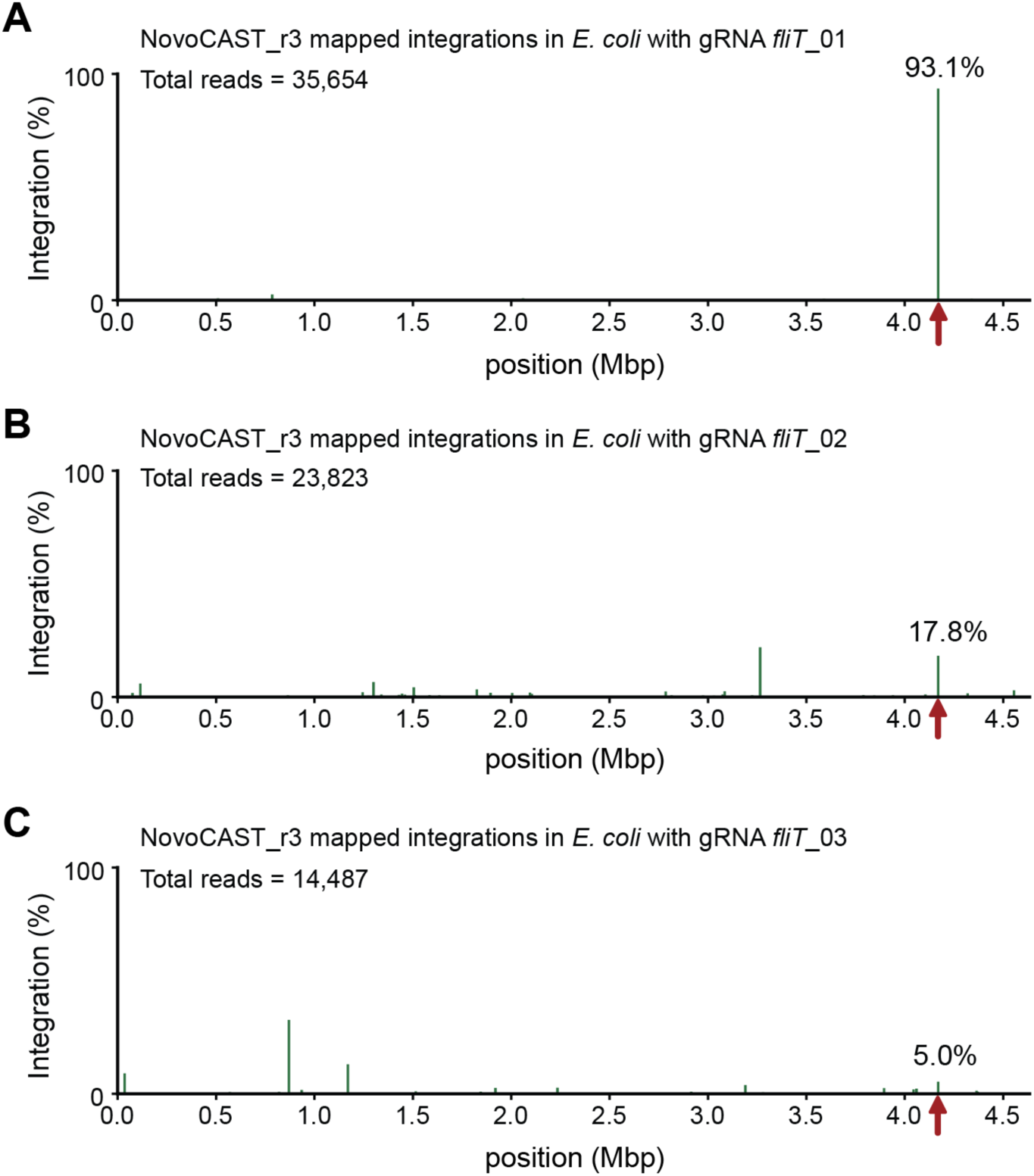
Genome-wide integration profile of NovoCAST_r3 in *E. coli* targeting *fliT.* Genome-wide distribution of mapped NovoCAST_r3 integration events for three dCas9 gRNAs (*fliT*_01, _02, and _03) targeting the flagellar gene *fliT*. Integration events are normalized by read count. The expected target site is indicated by red arrows. **A.** *fliT*_01 shows 93.1% on-target integration. **B**. *fliT*_02 shows 17.8% on-target integration. **C**. *fliT*_03 shows 5.0% on-target integration.

**Figure S12.**
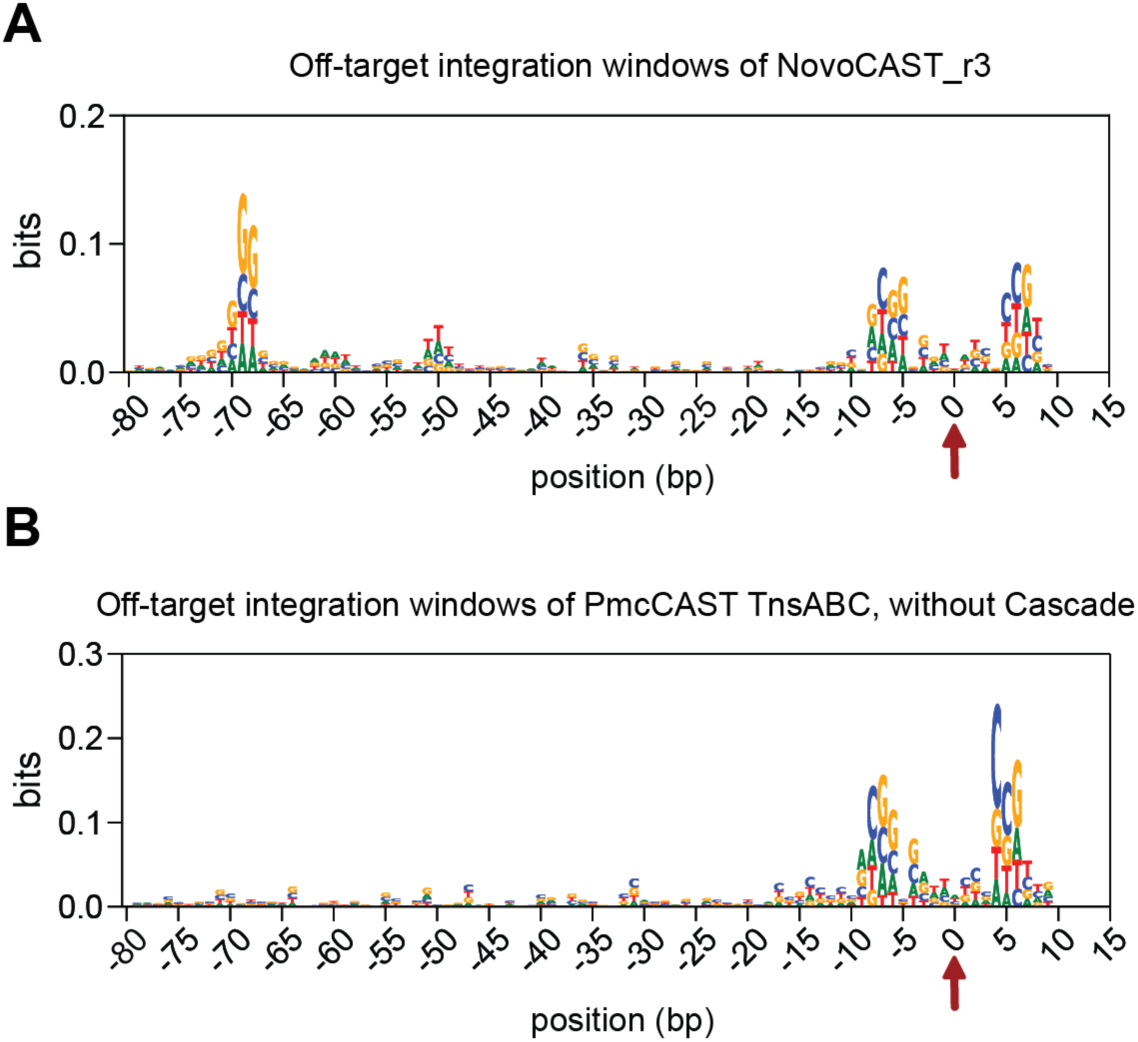
Off-target integration sequence preferences of NovoCAST_r3 compared with PmcCAST. Sequence logos showing nucleotide enrichment (bits) surrounding off-target integration events detected by genome-wide integration profiling in *E. coli.* Positions are plotted relative to the insertion site (position 0; red arrow). **A**. Sequence logo showing cumulative off-target integration events detected for NovoCAST_r3 across three *fliT*-targeting gRNAs. Modest nucleotide enrichment is observed near the insertion site (positions −10 to +10) and near the expected NovoCAST 5’-NGG-3’ PAM (position −70). The enrichment of nucleotides corresponding to the NovoCAST PAM occurs at the expected distance specified by the NovoCAST architecture. **B.** Sequence logo showing cumulative off-target integration events detected for PmcCAST TnsAB + TnsC (without CRISPR effectors). Sequence preferences at the insertion site (positions −10 to +10) closely match those observed in NovoCAST_r3, and, as expected, there is no enrichment of a 5’-NGG-3’ PAM 70 bp from the insertion site. These results indicate that NovoCAST retains the intrinsic target-site sequence features of the native CAST transposition machinery (PmcCAST TnsAB + TnsC) and imposes additional sequence preferences that correspond to the inclusion of dCas9 as the new CRISPR effector.

**Figure S13.**
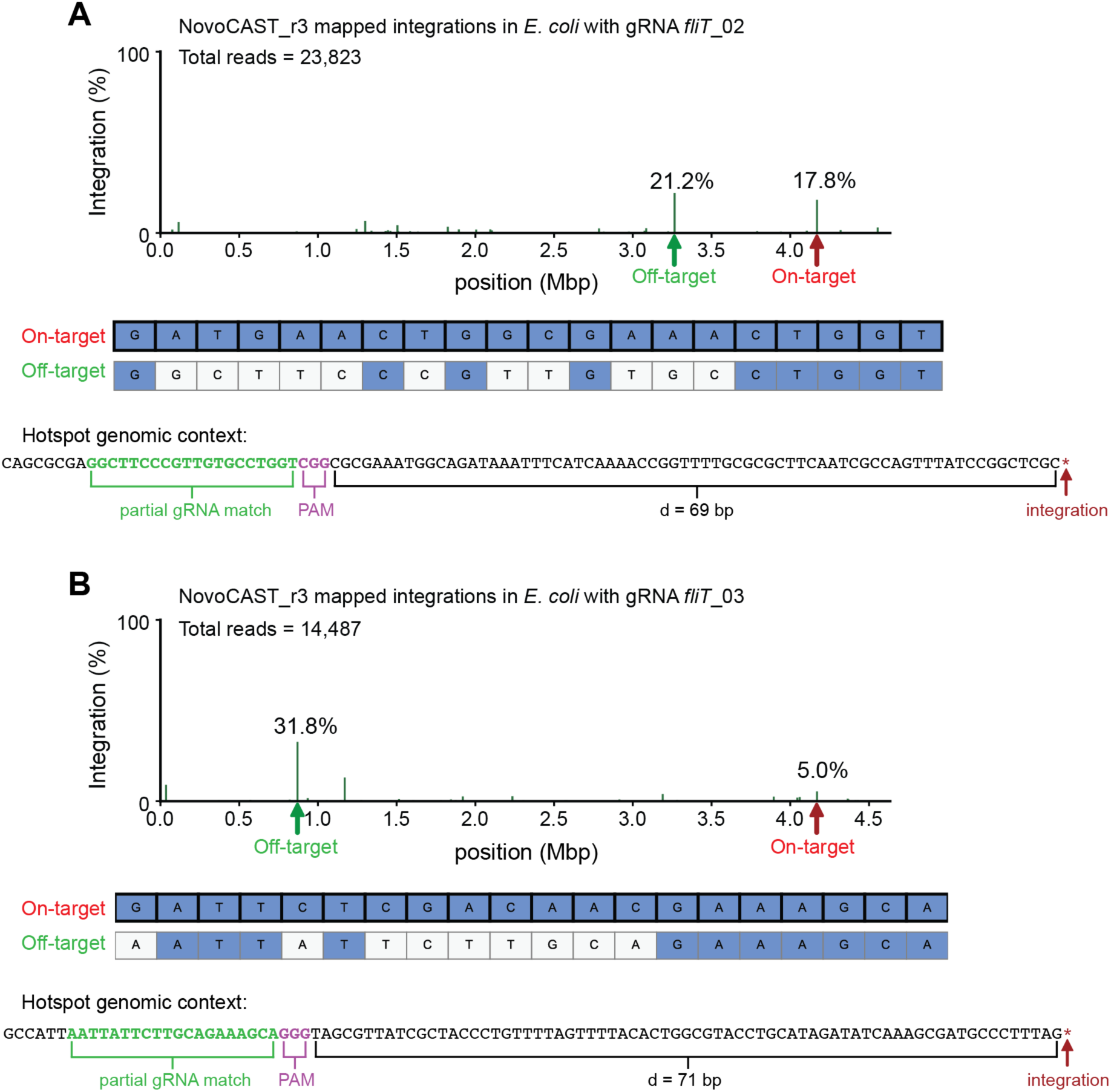
Off-target integration sites for *fliT*_02 and *fliT*_03 show partial gRNA matches. **A.** *fliT*_02, which was only 17.8% on-target (red arrow), had 21.2% of integration events occurring at an off-target genomic location (green arrow) containing a partial gRNA match. On-target and off-target spacers are shown below. Matches are colored blue. **B.** *fliT*_03, which was only 5.0% on-target, had 31.8% of integration events also occurring at an off-target genomic location containing a partial gRNA match.

**Figure S14.**
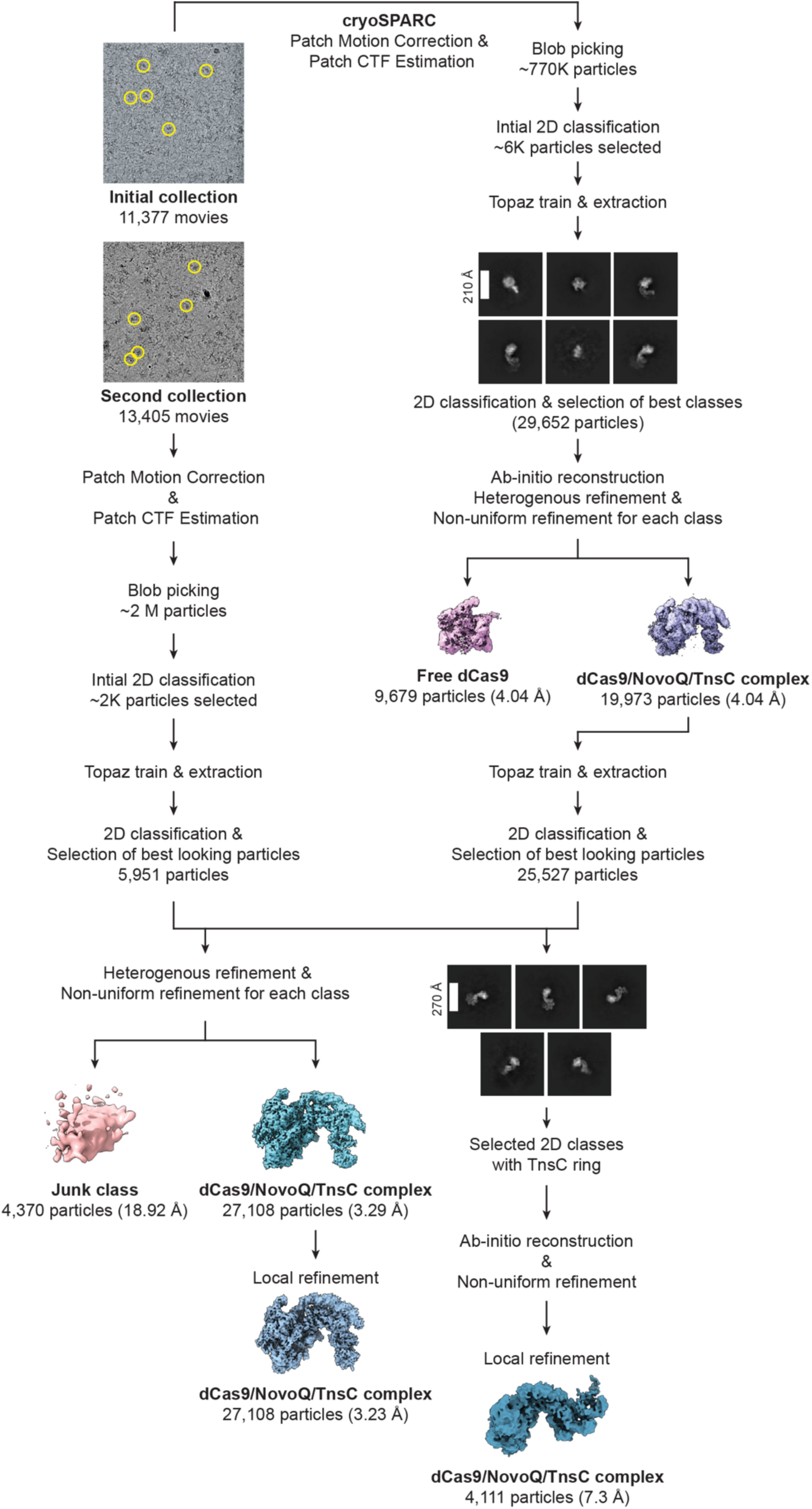
Cryo-EM data processing and NovoCAST reconstruction. Cryo-EM image processing workflow. Roughly two million particles were picked after patch motion correction and were subjected to iterative rounds of two-dimensional classification, and the best particles out of 2D classification were used for Topaz training. Representative micrographs are shown with representative particle picks (yellow circles). Representative 2D classes are shown; the white line indicates the scale bar (270 Å). Heterogeneous refinement separated free dCas9 particles from dCas9-NovoQ-TnsC complexes. Final refinement of the dCas9-NovoQ-TnsC complex yielded a reconstructed map at 3.23 Å resolution from 27,108 particles. A subset of particles containing a more complete TnsC ring was further processed by ab initio reconstruction and local refinement, yielding a lower-resolution map (7.3 Å) from 4,111 particles.

**Figure S15.**
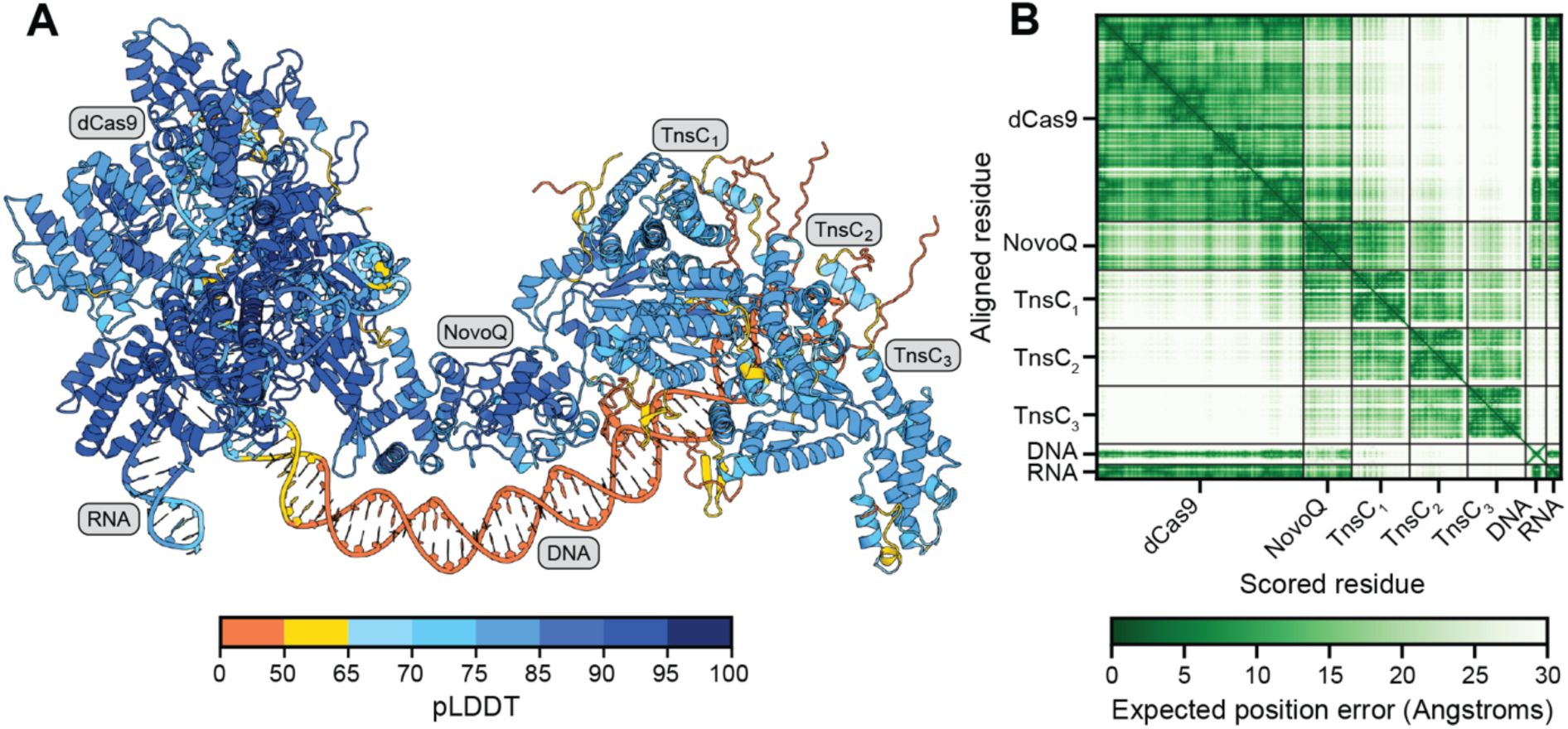
Predicted AlphaFold3 model of the NovoQ–dCas9–TnsC complex used to fit the Cryo-EM map. **A.** AlphaFold3 (AF3) prediction of the NovoCAST_r3 complex containing NovoQ_r3, dCas9_r2, three TnsC protomers (TnsC1–TnsC3), gRNA, and target DNA. The model is colored by per-residue confidence (pLDDT) according to AF3, with dark blue indicating higher confidence and orange indicating lower confidence regions of the predicted complex. This complex (NovoQ_r3–dCas9_r2–TnsC1) was used for fitting into the high-resolution cryo-EM map during model reconstruction. **B.** Predicted aligned error (PAE) matrix corresponding to the model shown in panel A. Dark green regions indicate low PAE, whereas lighter regions indicate high PAE indicative of increased uncertainty.

**Figure S16.**
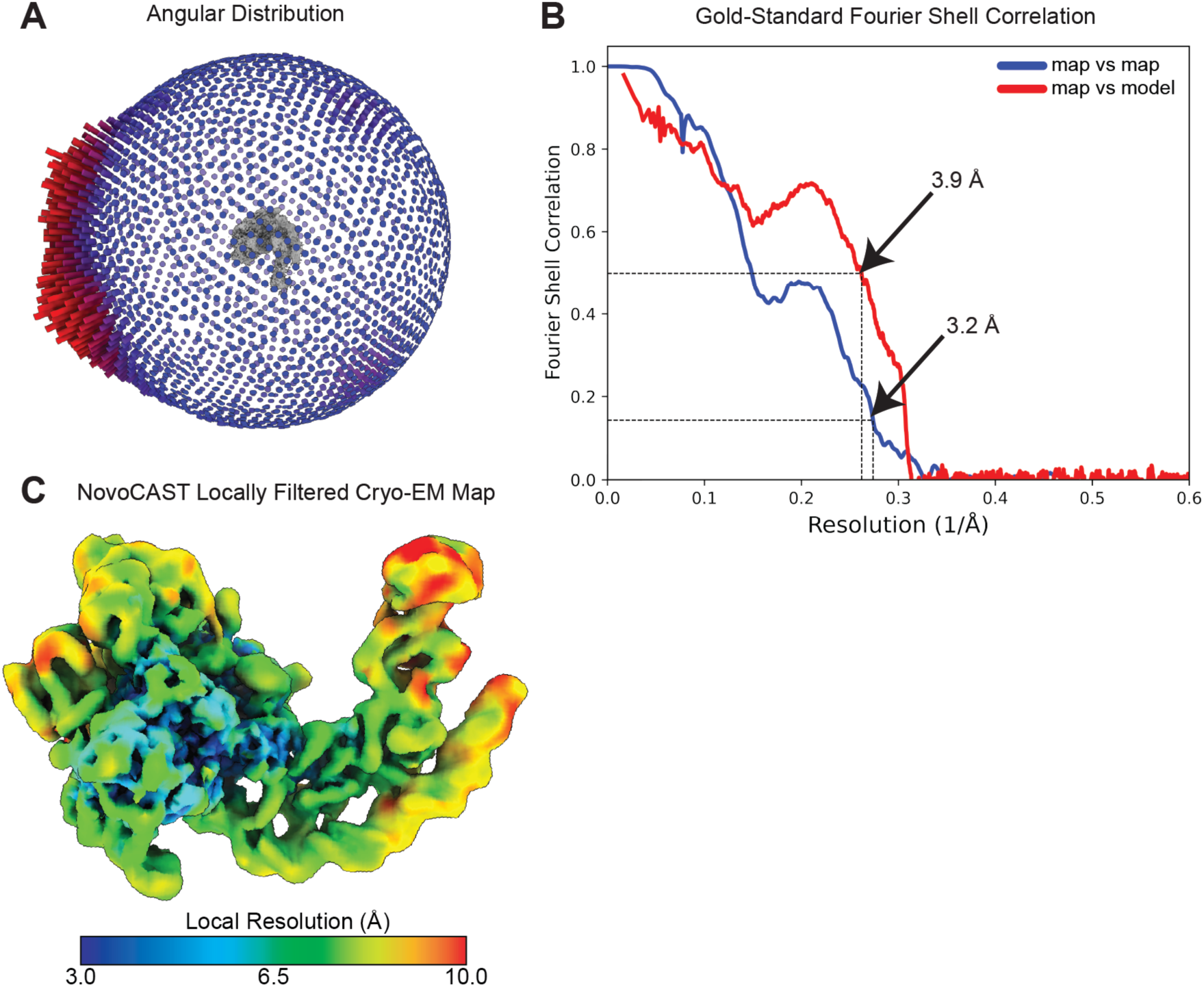
Cryo-EM data validation and locally filtered map of the NovoQ–dCas9–TnsC complex. **A.** Angular distribution plot of all NovoQ–dCas9–TnsC particles that contributed to the final map. The map and the angular distribution plot have the same orientation. The height and color (from blue to red) of the cylinders on the surface of the sphere are proportional to the number of particles in each view. **B.** Fourier shell correlation (FSC) curves for the final reconstruction. The bottom line indicates 0.143 standard cut-off for estimating overall resolution. The top line indicates 0.5 cutoff for estimating the model-map resolution for the predicted AlphaFold3 model and the high-resolution cryo-EM map. **C.** The local estimated resolution for each map voxel is colored and shown on a filtered version of the high-resolution cryo-EM map. Local resolution varies from 3 to 10 Å, the legend below indicates the resolution value. Lower-resolution regions correspond to more flexible portions of the density map, particularly at the distal ends of the target DNA furthest from dCas9 and the TnsC protomer.

**Figure S17.**
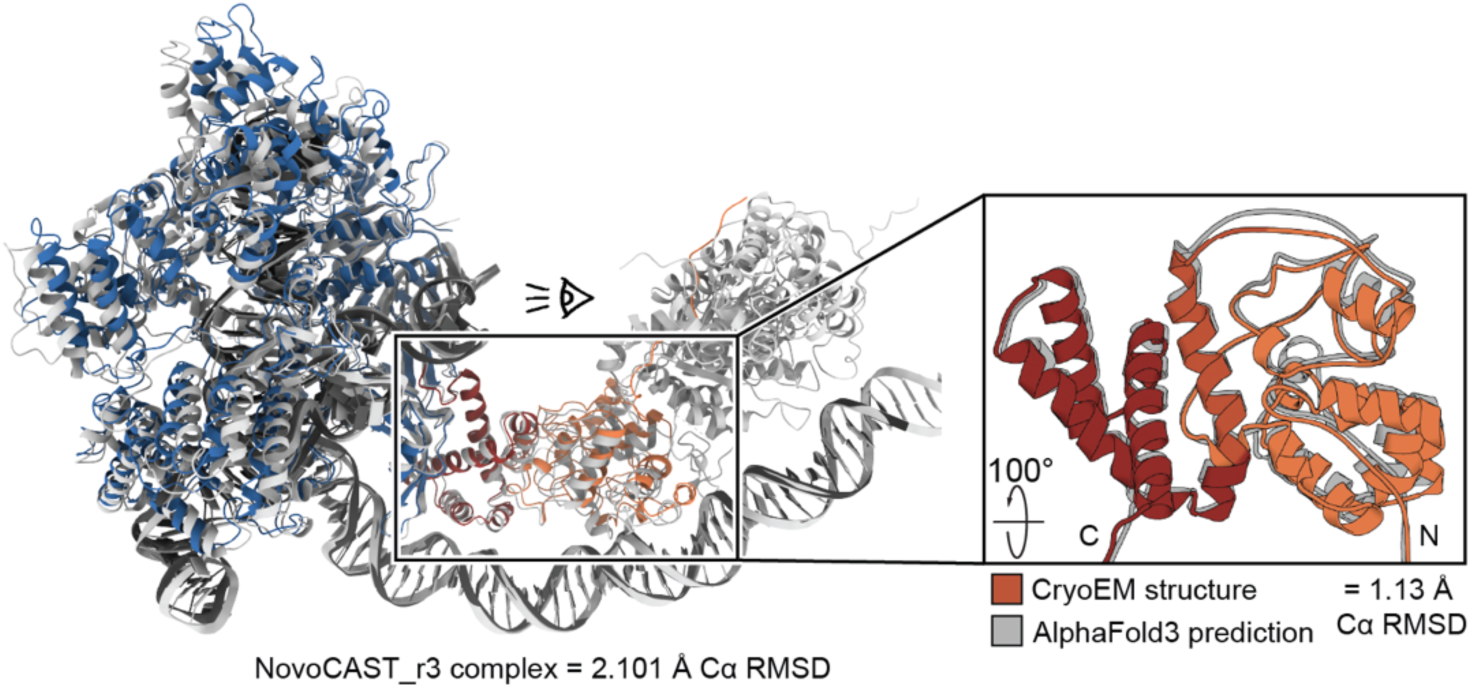
The NovoQ_r3–dCas9_r2 cryo-EM model compared to the NovoQ_r3–dCas9_r2 design prediction. The refined model of NovoQ_r3–dCas9_r2 from the reconstructed map at 3.23 Å is colored (NovoQ_r3, orange and red; dCas9_r2, blue) and aligned to the AlphaFold3 prediction of NovoQ_r3–dCas9_r2 (light gray). The global Cα RMSD between the NovoQ_r3–dCas9_r2 prediction and the refined model (left) is 2.101 Å. The Cα RMSD for NovoQ_r3 alone is 1.113 Å (right panel).

**Figure S18.**
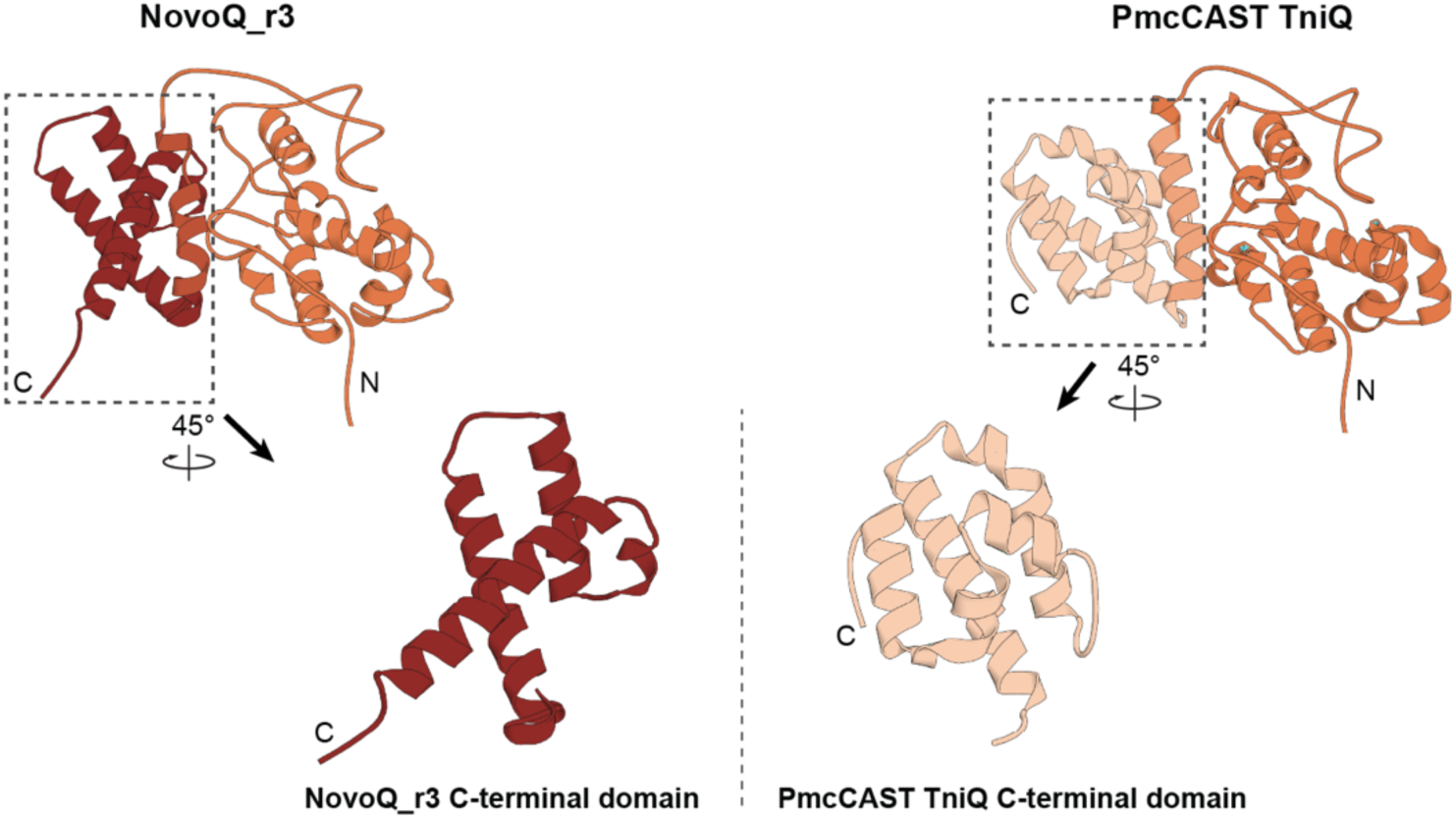
Structural comparison between the de novo NovoQ_r3 C-terminal domain and the native PmcCAST TniQ C-terminal domain. Overall structures on the top row depict NovoQ_r3 (left) and PmcCAST TniQ (right). The bottom row shows the corresponding C-terminal domains highlighted in the dashed boxes above. Both extracted C-terminal domains are rotated by 45 degrees for visual clarity. The TM-score between the C-terminal domains of NovoQ_r3 and PmcCAST TniQ is 0.43 over 87 structurally aligned residues. N- and C-termini of TniQ are labeled N and C, respectively. The N-terminal domains are colored orange, the C-terminal domain of PmcCAST TniQ is colored beige, and the *de novo* C-terminal domain of NovoQ_r3 is colored red.

**Figure S19.**
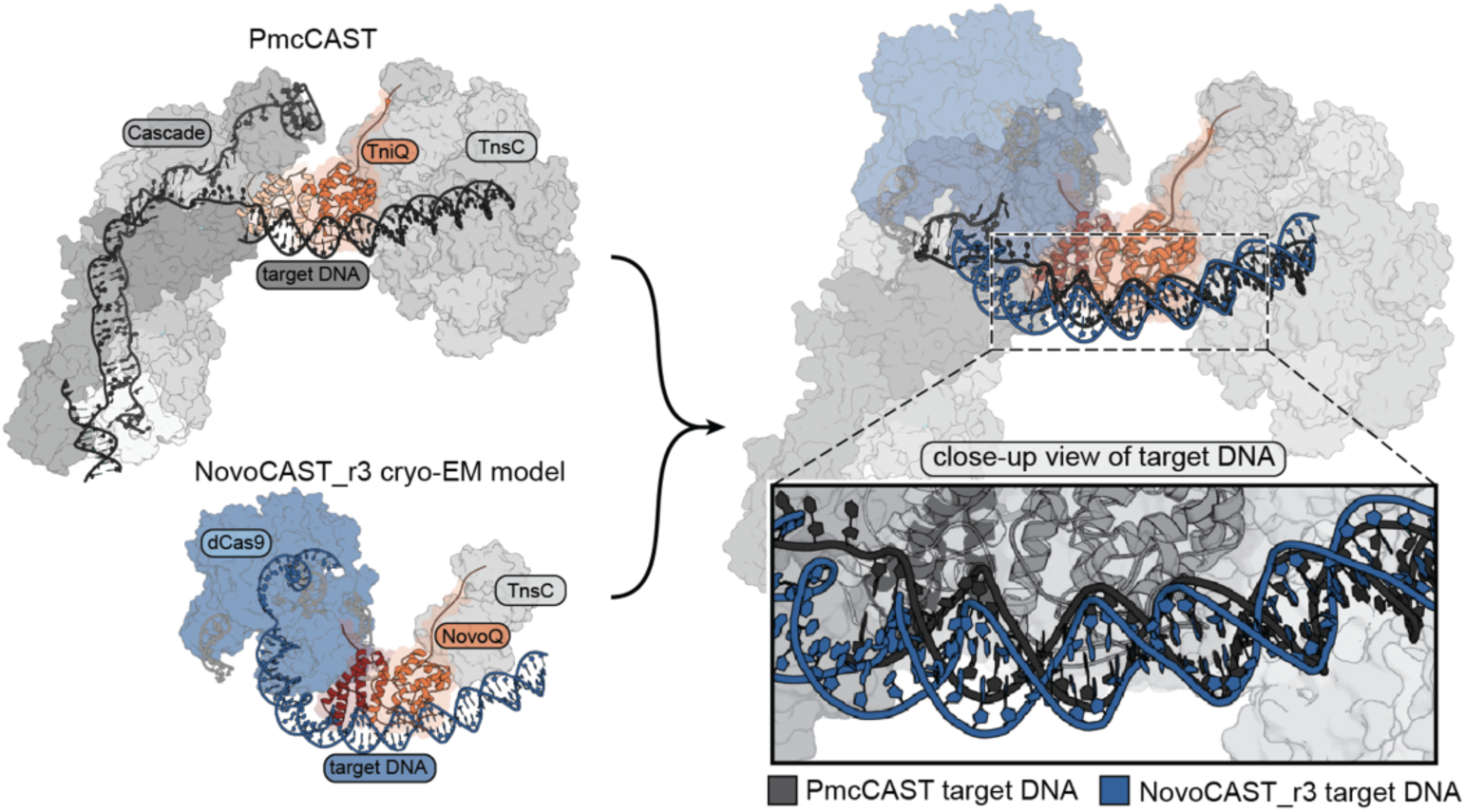
NovoCAST_r3 target DNA distortions resemble those observed in PmcCAST. Top left: Type I-B2 PmcCAST is shown (PDB: 8FF4), highlighting the target DNA (dark gray). Bottom left: the experimentally determined atomic model of NovoCAST_r3 is displayed, highlighting the target DNA (dark blue). Both structures are shown in views that are aligned along the TniQ N-terminal domain (orange). Right: superposition of PmcCAST and NovoCAST_r3. The target DNA of both PmcCAST (dark gray) and NovoCAST_r3 (dark blue) is shown in a close-up view (boxed inset), showing a similar DNA curvature in both systems.

**Figure S20.**
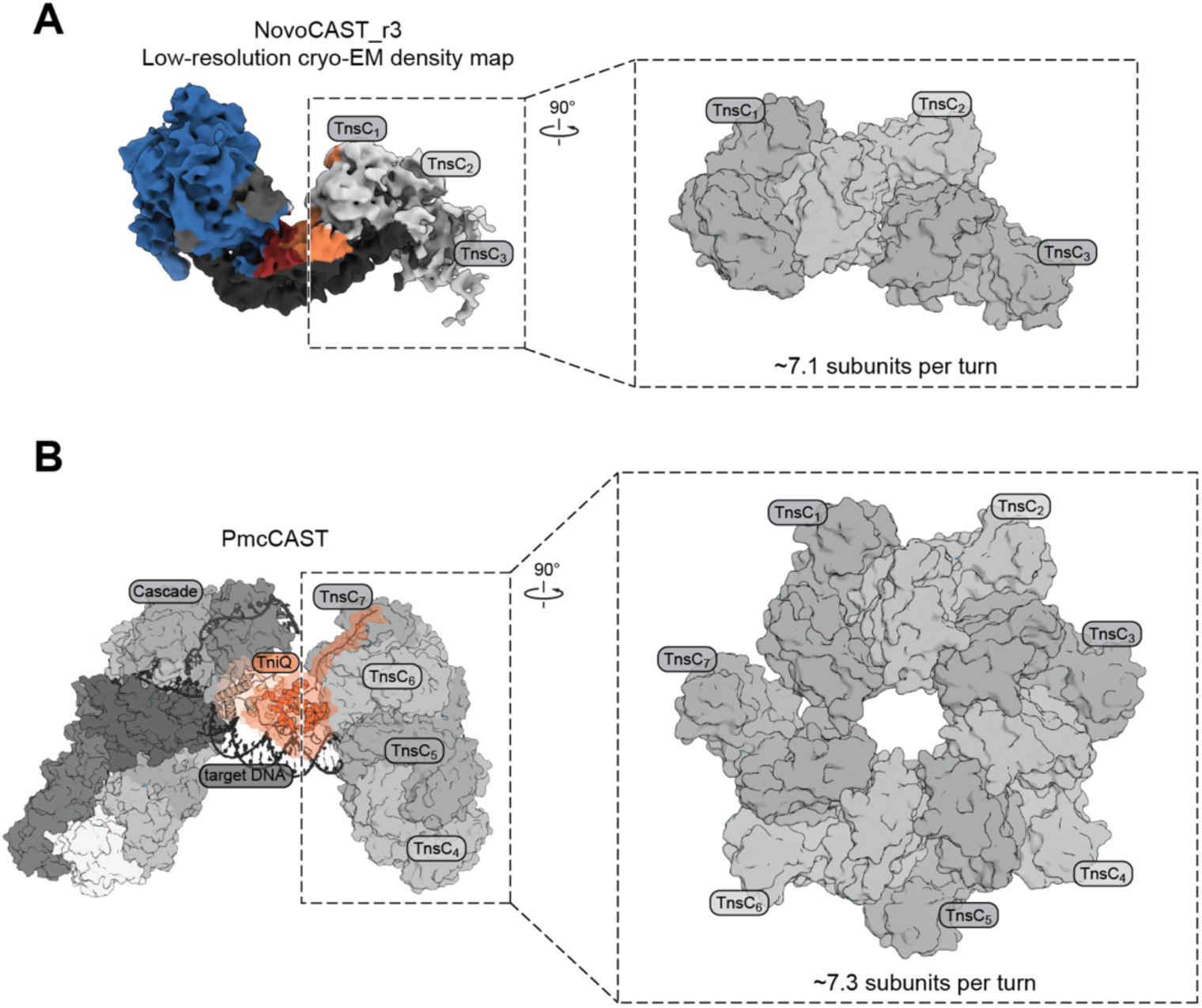
TnsC protomers match the canonical heptameric TnsC assembly. **A.** Left: Cryo-EM map of NovoCAST_r3. TnsC protomers are labeled. Right: Docked TnsC protomers shown in an orthogonal viewing direction (rotation indicated). Helical parameters from the docked TnsC protomers correspond to ∼7.1 protomer subunits per turn. **B.** PmcCAST assembly (PDB: 8FF4). Orthogonal views (rotation and axis indicated) of the TnsC heptamer, helical parameters computed from this arrangement correspond to ∼7.3 protomer subunits per turn. The slightly different helical parameters observed in this arrangement correspond to the gap between TnsC_1_ and TnsC_7_, which breaks the symmetry of the heptameric arrangement.

**Figure S21.**
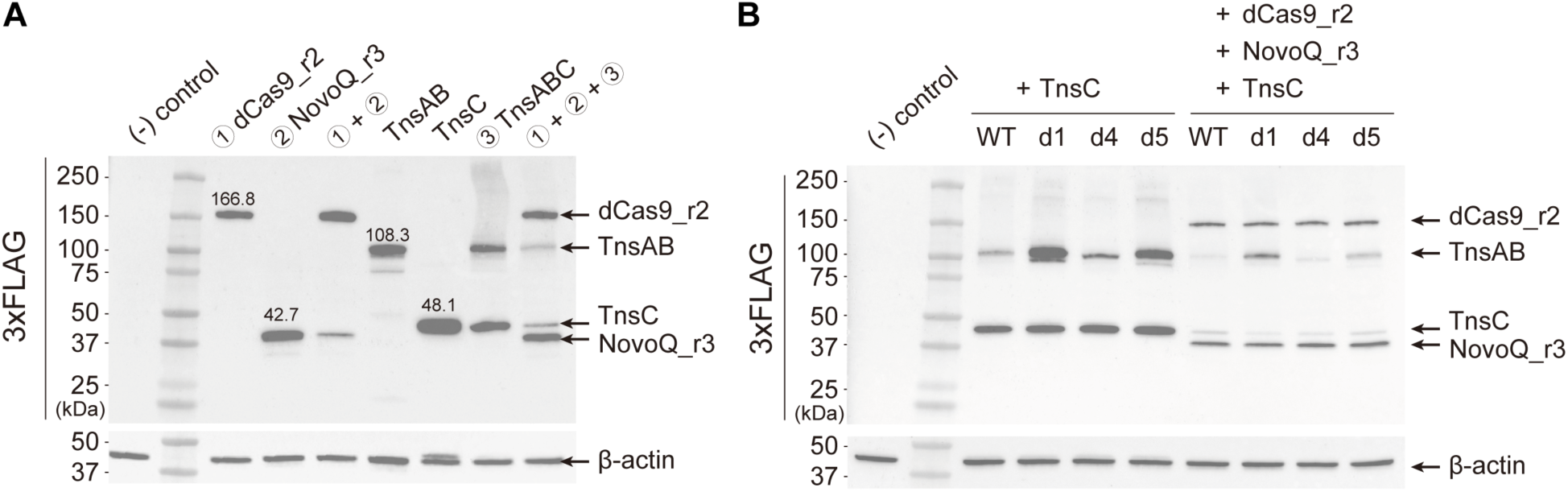
Expression of TnsAB redesigns in human cells. **A.** Western blot of NovoCAST_r3 components detected with anti-FLAG antibodies. **B.** Western blot of TnsAB redesigns co-expressed with NovoCAST_r3 components, detected using anti-FLAG antibodies. WT, wild-type PmcCAST TnsAB, d1, TnsAB_d1; d4, TnsAB_d4; d5, TnsAB_d5. All lanes include a TnsAB variant, samples differ depending on whether only TnsC was co-transfected (left) or if all components were co-transfected (right). For both panels, the (-) control represents no transfection. Molecular weights of the protein ladder (Lane 2) are indicated on the left side. Proteins are indicated by arrows, and their molecular weights (kDa) are indicated on the membrane. β-actin was used as a loading control and detected with anti-β-actin antibodies.

**Figure S22.**
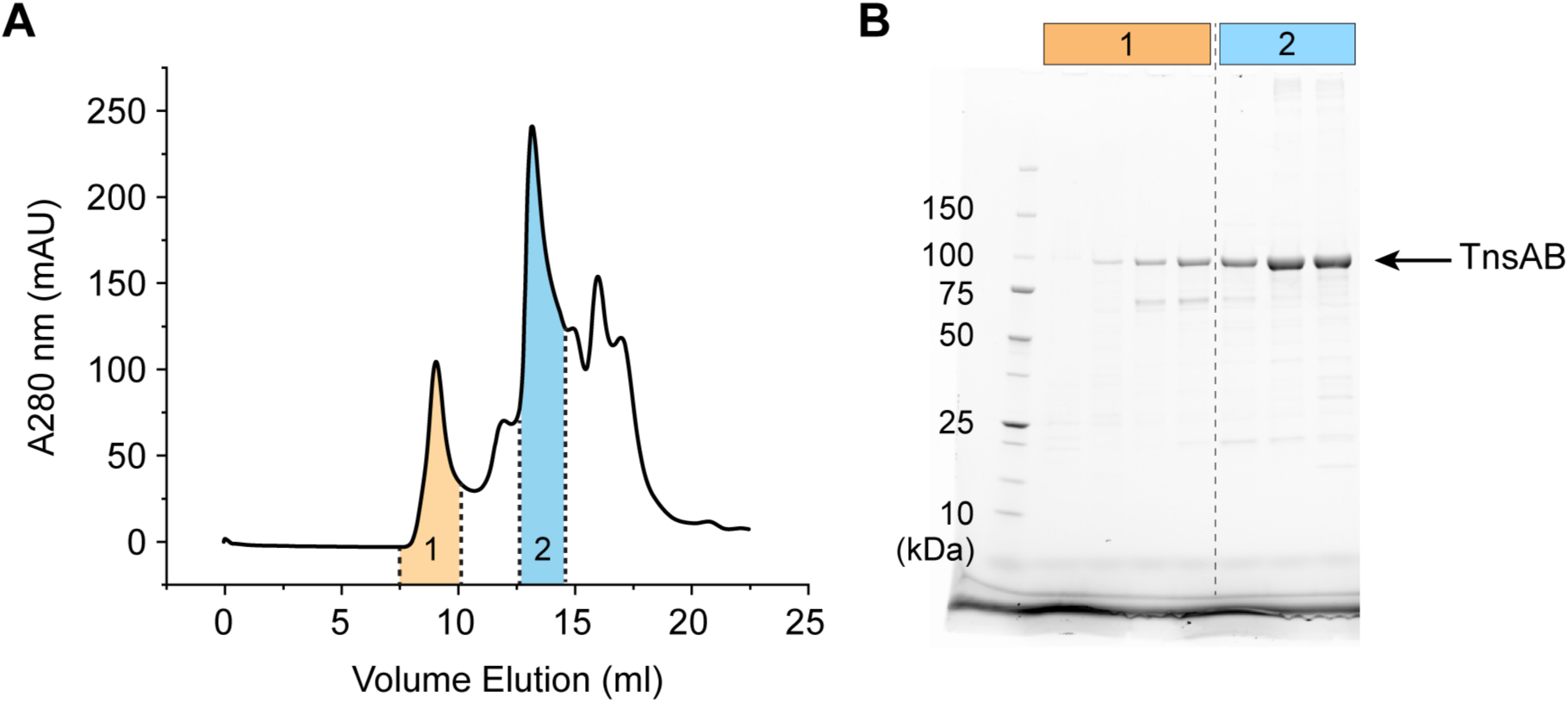
Size exclusion chromatography of purified TnsAB. **A.** UV/Vis chromatogram of TnsAB size exclusion chromatography. TnsAB was overexpressed, purified using a heparin column, and subjected to size exclusion chromatography on a Superdex 200 Increase 10/300 GL column. Fractions 1 (orange) and 2 (blue) are indicated on the graph. Fraction 1 shows a significant signal in the void volume, suggesting a tendency of PmcCAST TnsAB to aggregate. **B.** Coomassie-stained SDS-PAGE of fractions 1 and 2. Molecular weights of the protein ladder (lane 1) are indicated on the left, and the lanes corresponding to each fraction are indicated at the top.

**Figure S23.**
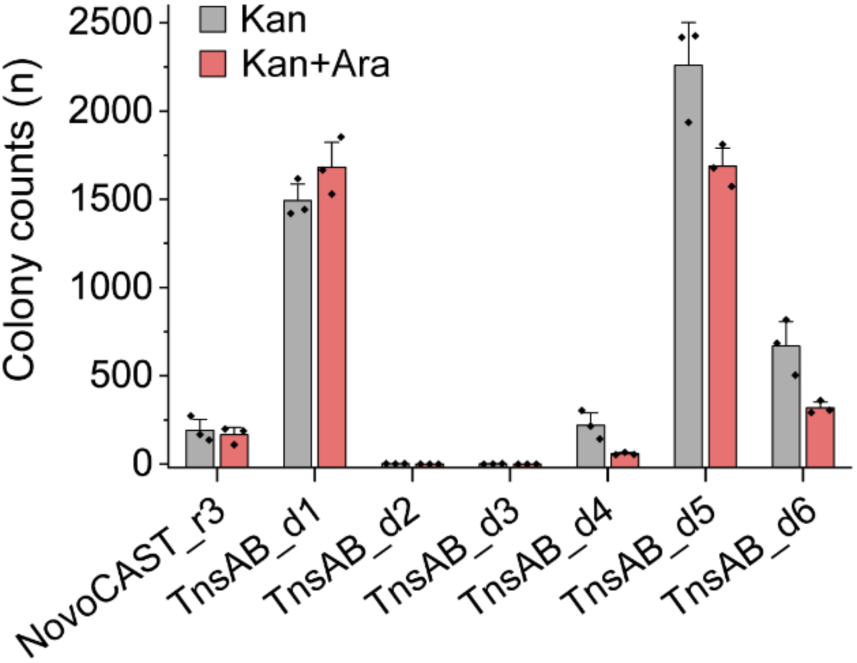
Redesign of TnsAB enhances integration activity of NovoCAST in *E. coli*. Integration efficiency of TnsAB redesigns was measured using an *in vivo* transposition assay in *E. coli*. Bars with black diamonds represent colony counts from Kan selection (overall activity), and bars with red diamonds represent colony counts from Kan+Ara selection (on-target activity). All data represent the mean ± standard deviation; n = 3 for each bar.

**Figure S24.**
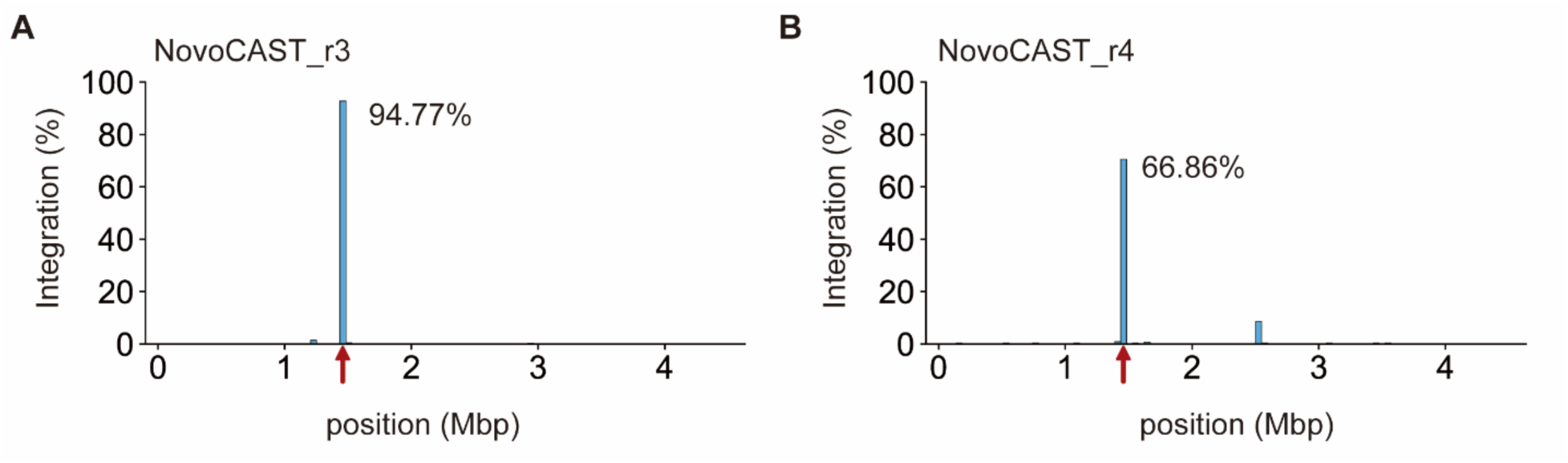
Comparison of integration profiles between NovoCAST_r3 and NovoCAST_r4. Integration profiles of **A.** NovoCAST_r3 and **B.** NovoCAST_r4 (corresponding to NovoCAST_r3 + TnsAB_d5) are shown. All integration sites were analyzed using genome-wide integration profiling and mapped onto the *E. coli* genome. The target site guided by the *ccdB*_01 gRNA is indicated by a red arrow, and the percentage of integrations at the target site is shown next to each histogram. The sample analyzed in panel **A** was generated independently from the sample analyzed in Figure 2M **& 3A**.

**Figure S25.**
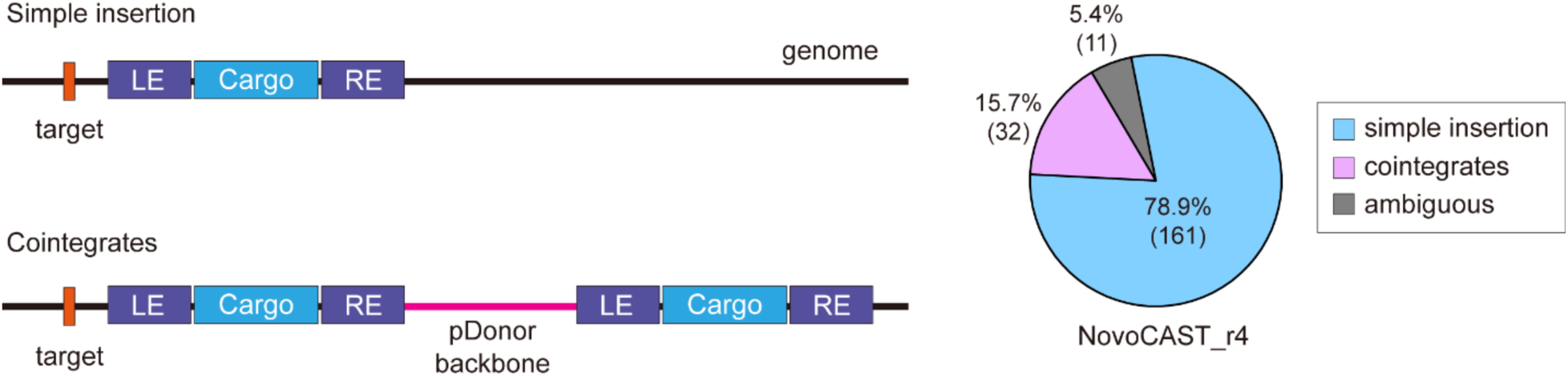
Cointegration analysis of NovoCAST_r4 in *E. coli*. Schematics of simple insertion and cointegrate products are shown in the left panel. In the cointegrate product, the pDonor backbone (pink line) is integrated into the genome (black line) along with duplicated donor DNA (LE-Cargo-RE). The proportions of simple insertions (blue), cointegrates (pink), and ambiguous reads (gray) produced by NovoCAST_r4 are shown as a pie chart in the right panel, with the number of sequencing reads indicated in parentheses. Reads that could not be classified as either simple insertions or cointegrates due to the insufficient read length were classified as ambiguous.

**Supplementary Table S8:**
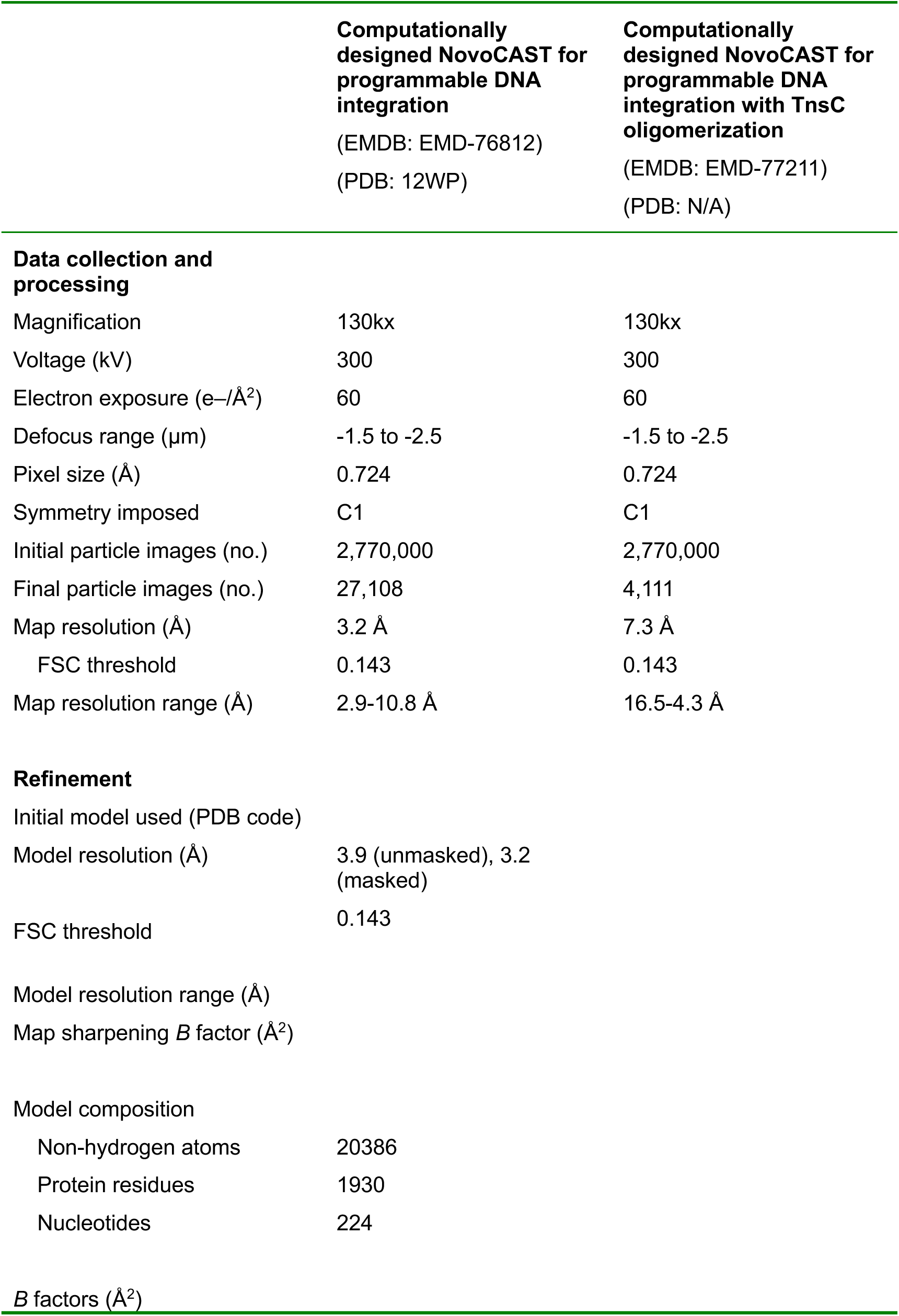
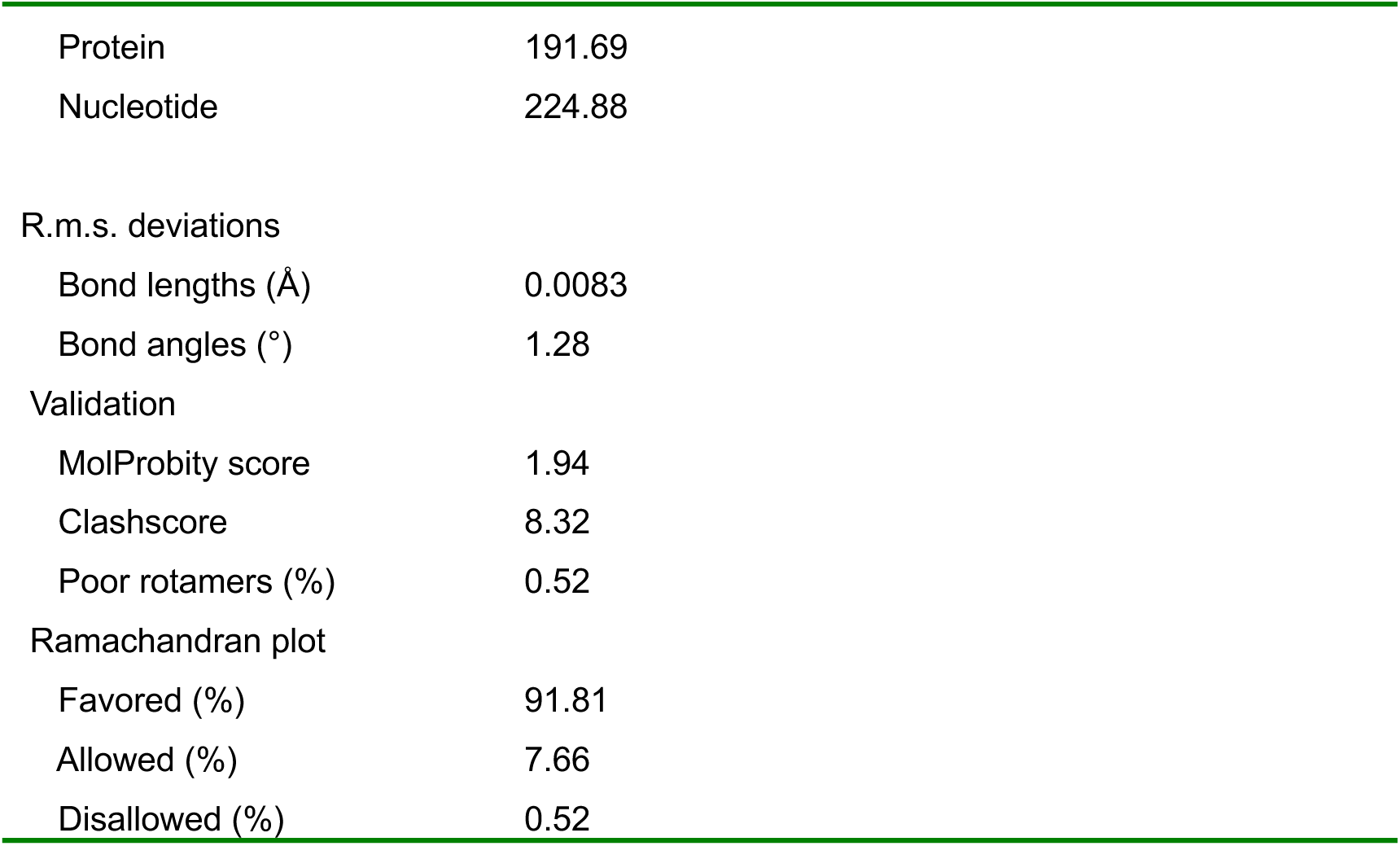
Cryo-EM data collection, refinement, and validation statistics.

|  | Computationally designed NovoCAST for programmable DNA integration<br>(EMDB: EMD-76812)<br>(PDB: 12WP) | Computationally designed NovoCAST for programmable DNA integration with TnsC oligomerization<br>(EMDB: EMD-77211)<br>(PDB: N/A) |
| --- | --- | --- |
| **Data collection and processing** |  |  |
| Magnification | 130kx | 130kx |
| Voltage (kV) | 300 | 300 |
| Electron exposure (e-/Å²) | 60 | 60 |
| Defocus range (µm) | -1.5 to -2.5 | -1.5 to -2.5 |
| Pixel size (Å) | 0.724 | 0.724 |
| Symmetry imposed | C1 | C1 |
| Initial particle images (no.) | 2,770,000 | 2,770,000 |
| Final particle images (no.) | 27,108 | 4,111 |
| Map resolution (Å) | 3.2 Å | 7.3 Å |
| FSC threshold | 0.143 | 0.143 |
| Map resolution range (Å) | 2.9-10.8 Å | 16.5-4.3 Å |
| **Refinement** |  |  |
| Initial model used (PDB code) |  |  |
| Model resolution (Å) | 3.9 (unmasked), 3.2 (masked) |  |
| FSC threshold | 0.143 |  |
| Model resolution range (Å) |  |  |
| Map sharpening *B* factor (Å²) |  |  |
| **Model composition** |  |  |
| Non-hydrogen atoms | 20386 |  |
| Protein residues | 1930 |  |
| Nucleotides | 224 |  |
| ***B* factors (Å²)** |  |  |

|  |  |
| --- | --- |
| Protein | 191.69 |
| Nucleotide | 224.88 |
| R.m.s. deviations |  |
| Bond lengths (Å) | 0.0083 |
| Bond angles (°) | 1.28 |
| Validation |  |
| MolProbity score | 1.94 |
| Clashscore | 8.32 |
| Poor rotamers (%) | 0.52 |
| Ramachandran plot |  |
| Favored (%) | 91.81 |
| Allowed (%) | 7.66 |
| Disallowed (%) | 0.52 |

## References

1. “Accurate Structure Prediction of Biomolecular Interactions with AlphaFold 3 | Nature.” n.d. Accessed March 11, 2026. https://www.nature.com/articles/s41586-024-07487-w.

2. Alford, Rebecca F., Andrew Leaver-Fay, Jeliazko R. Jeliazkov, et al. 2017. “The Rosetta All-Atom Energy Function for Macromolecular Modeling and Design.” Journal of Chemical Theory and Computation 13 (6): 3031–48. 10.1021/acs.jctc.7b00125.

3. Altae-Tran, Han, Soumya Kannan, F. Esra Demircioglu, et al. 2021. “The Widespread IS200/IS605 Transposon Family Encodes Diverse Programmable RNA-Guided Endonucleases.” Science (New York, N.Y.) 374 (6563): 57–65. 10.1126/science.abj6856.

4. Anzalone, Andrew V., Peyton B. Randolph, Jessie R. Davis, et al. 2019. “Search-and-Replace Genome Editing without Double-Strand Breaks or Donor DNA.” Nature 576 (7785): 149–57. 10.1038/s41586-019-1711-4.

5. Arinkin, Vladimir, Georgy Smyshlyaev, and Orsolya Barabas. 2019. “Jump Ahead with a Twist: DNA Acrobatics Drive Transposition Forward.” Current Opinion in Structural Biology 59 (December): 168–77. 10.1016/j.sbi.2019.08.006.

6. Cong, Le, F. Ann Ran, David Cox, et al. 2013. “Multiplex Genome Engineering Using CRISPR/Cas Systems.” Science (New York, N.Y.) 339 (6121): 819–23. 10.1126/science.1231143.

7. Craig, Nancy Lynn. 2015. Mobile DNA III. ASM press.

8. Dauparas, Justas, Gyu Rie Lee, Robert Pecoraro, et al. 2025. “Atomic Context-Conditioned Protein Sequence Design Using LigandMPNN.” Nature Methods 22 (4): 717–23. 10.1038/s41592-025-02626-1.

9. Davey, James A., and Roberto A. Chica. 2012. “Multistate Approaches in Computational Protein Design.” Protein Science: A Publication of the Protein Society 21 (9): 1241–52. 10.1002/pro.2128.

10. Doench, John G., Nicolo Fusi, Meagan Sullender, et al. 2016. “Optimized sgRNA Design to Maximize Activity and Minimize Off-Target Effects of CRISPR-Cas9.” Nature Biotechnology 34 (2): 184–91. 10.1038/nbt.3437.

11. Durrant, Matthew G., Nicholas T. Perry, James J. Pai, et al. 2024. “Bridge RNAs Direct Programmable Recombination of Target and Donor DNA.” Nature 630 (8018): 984–93. 10.1038/s41586-024-07552-4.

12. Eyquem, Justin, Jorge Mansilla-Soto, Theodoros Giavridis, et al. 2017. “Targeting a CAR to the TRAC Locus with CRISPR/Cas9 Enhances Tumour Rejection.” Nature 543 (7643): 113–17. 10.1038/nature21405.

13. Frangoul, Haydar, David Altshuler, M. Domenica Cappellini, et al. 2021. “CRISPR-Cas9 Gene Editing for Sickle Cell Disease and β-Thalassemia.” The New England Journal of Medicine 384 (3): 252–60. 10.1056/NEJMoa2031054.

14. Gaudelli, Nicole M., Alexis C. Komor, Holly A. Rees, et al. 2017. “Programmable Base Editing of A•T to G•C in Genomic DNA without DNA Cleavage.” Nature 551 (7681): 464–71. 10.1038/nature24644.

15. George, Jerrin Thomas, Christopher Acree, Jung-Un Park, et al. 2023. “Mechanism of Target Site Selection by Type V-K CRISPR-Associated Transposases.” Science (New York, N.Y.) 382 (6672): eadj8543. 10.1126/science.adj8543.

16. Haiko, Johanna, and Benita Westerlund-Wikström. 2013. “The Role of the Bacterial Flagellum in Adhesion and Virulence.” Biology 2 (4): 1242–67. 10.3390/biology2041242.

17. Halling, S. M., and N. Kleckner. 1982. “A Symmetrical Six-Base-Pair Target Site Sequence Determines Tn10 Insertion Specificity.” Cell 28 (1): 155–63. 10.1016/0092-8674(82)90385-3.

18. Halpin-Healy, Tyler S., Sanne E. Klompe, Samuel H. Sternberg, and Israel S. Fernández. 2020. “Structural Basis of DNA Targeting by a Transposon-Encoded CRISPR–Cas System.” Nature 577 (7789): 271–74. 10.1038/s41586-019-1849-0.

19. Hoffmann, Florian T., Minjoo Kim, Leslie Y. Beh, et al. 2022. “Selective TnsC Recruitment Enhances the Fidelity of RNA-Guided Transposition.” Nature 609 (7926): 384–93. 10.1038/s41586-022-05059-4.

20. Hoose, Alex, Richard Vellacott, Marko Storch, Paul S. Freemont, and Maxim G. Ryadnov. 2023. “DNA Synthesis Technologies to Close the Gene Writing Gap.” Nature Reviews Chemistry 7 (3): 144–61. 10.1038/s41570-022-00456-9.

21. Hsieh, Shan-Chi, and Joseph E. Peters. 2024. “Natural and Engineered Guide RNA–Directed Transposition with CRISPR-Associated Tn7-Like Transposons.” Annual Review of Biochemistry 93 (1): 139–61. 10.1146/annurev-biochem-030122-041908.

22. Ikeda, Arisa, Wataru Fujii, Koji Sugiura, and Kunihiko Naito. 2019. “High-Fidelity Endonuclease Variant HypaCas9 Facilitates Accurate Allele-Specific Gene Modification in Mouse Zygotes.” Communications Biology 2: 371. 10.1038/s42003-019-0627-8.

23. Jinek, Martin, Krzysztof Chylinski, Ines Fonfara, Michael Hauer, Jennifer A. Doudna, and Emmanuelle Charpentier. 2012. “A Programmable Dual-RNA–Guided DNA Endonuclease in Adaptive Bacterial Immunity.” Science 337 (6096): 816–21. 10.1126/science.1225829.

24. Juhas, Mario, Lewis D. B. Evans, Joe Frost, et al. 2014. “Escherichia Coli Flagellar Genes as Target Sites for Integration and Expression of Genetic Circuits.” PLOS ONE 9 (10): e111451. 10.1371/journal.pone.0111451.

25. Kleinstiver, Benjamin P., Vikram Pattanayak, Michelle S. Prew, et al. 2016. “High-Fidelity CRISPR-Cas9 Nucleases with No Detectable Genome-Wide off-Target Effects.” Nature 529 (7587): 490–95. 10.1038/nature16526.

26. Kleinstiver, Benjamin P., Michelle S. Prew, Shengdar Q. Tsai, et al. 2015. “Engineered CRISPR-Cas9 Nucleases with Altered PAM Specificities.” Nature 523 (7561): 481–85. 10.1038/nature14592.

27. Klompe, Sanne E., Phuc L. H. Vo, Tyler S. Halpin-Healy, and Samuel H. Sternberg. 2019. “Transposon-Encoded CRISPR–Cas Systems Direct RNA-Guided DNA Integration.” Nature 571 (7764): 219–25. 10.1038/s41586-019-1323-z.

28. Kosicki, Michael, Kärt Tomberg, and Allan Bradley. 2018. “Repair of Double-Strand Breaks Induced by CRISPR–Cas9 Leads to Large Deletions and Complex Rearrangements.” Nature Biotechnology 36 (8): 765–71. 10.1038/nbt.4192.

29. Kovač, Adrian, Csaba Miskey, Michael Menzel, Esther Grueso, Andreas Gogol-Döring, and Zoltán Ivics. 2020. “RNA-Guided Retargeting of Sleeping Beauty Transposition in Human Cells.” eLife 9 (March): e53868. 10.7554/eLife.53868.

30. Kuduvalli, Prasad N., Jason E. Rao, and Nancy L. Craig. 2001. “Target DNA Structure Plays a Critical Role in Tn7 Transposition.” The EMBO Journal 20 (4): 924–32. 10.1093/emboj/20.4.924.

31. Kuscu, Cem, Sevki Arslan, Ritambhara Singh, Jeremy Thorpe, and Mazhar Adli. 2014. “Genome-Wide Analysis Reveals Characteristics of off-Target Sites Bound by the Cas9 Endonuclease.” Nature Biotechnology 32 (7): 677–83. 10.1038/nbt.2916.

32. Lampe, George D., Rebeca T. King, Tyler S. Halpin-Healy, et al. 2024. “Targeted DNA Integration in Human Cells without Double-Strand Breaks Using CRISPR-Associated Transposases.” Nature Biotechnology 42 (1): 87–98. 10.1038/s41587-023-01748-1.

33. Liu, Jason, Daniela S. Aliaga Goltsman, Lisa M. Alexander, et al. 2025. “Integration of Therapeutic Cargo into the Human Genome with Programmable Type V-K CAST.” Nature Communications 16 (1): 2427. 10.1038/s41467-025-57416-2.

34. Mali, Prashant, Luhan Yang, Kevin M. Esvelt, et al. 2013. “RNA-Guided Human Genome Engineering via Cas9.” Science (New York, N.Y.) 339 (6121): 823–26. 10.1126/science.1232033.

35. Montaño, Sherwin P., Ying Z. Pigli, and Phoebe A. Rice. 2012. “The μ Transpososome Structure Sheds Light on DDE Recombinase Evolution.” Nature 491 (7424): 413–17. 10.1038/nature11602.

36. Pandey, Smriti, Xin D. Gao, Nicholas A. Krasnow, et al. 2025. “Efficient Site-Specific Integration of Large Genes in Mammalian Cells via Continuously Evolved Recombinases and Prime Editing.” Nature Biomedical Engineering 9 (1): 22–39. 10.1038/s41551-024-01227-1.

37. Park, Jung-Un, Amy Wei-Lun Tsai, Tiffany H. Chen, Joseph E. Peters, and Elizabeth H. Kellogg. 2022. “Mechanistic Details of CRISPR-Associated Transposon Recruitment and Integration Revealed by Cryo-EM.” Proceedings of the National Academy of Sciences of the United States of America 119 (32): e2202590119. 10.1073/pnas.2202590119.

38. Park, Jung-Un, Amy Wei-Lun Tsai, Eshan Mehrotra, et al. 2021. “Structural Basis for Target Site Selection in RNA-Guided DNA Transposition Systems.” Science 373 (6556): 768–74. 10.1126/science.abi8976.

39. Park, Jung-Un, Amy Wei-Lun Tsai, Alexandrea N. Rizo, et al. 2023. “Structures of the Holo CRISPR RNA-Guided Transposon Integration Complex.” Nature 613 (7945): 775–82. 10.1038/s41586-022-05573-5.

40. Park, Seong Guk, Jung-Un Park, Esteban Dodero-Rojas, John A. Bryant Jr, Geetha Sankaranarayanan, and Elizabeth H. Kellogg. 2025. “Comprehensive Profiling of Activity and Specificity of RNA-Guided Transposons Reveals Opportunities to Engineer Improved Variants.” Nucleic Acids Research 53 (18): gkaf917. 10.1093/nar/gkaf917.

41. Peters, Joseph E., Kira S. Makarova, Sergey Shmakov, and Eugene V. Koonin. 2017. “Recruitment of CRISPR-Cas Systems by Tn7-like Transposons.” Proceedings of the National Academy of Sciences of the United States of America 114 (35): E7358–66. 10.1073/pnas.1709035114.

42. Saito, Makoto, Alim Ladha, Jonathan Strecker, et al. 2021. “Dual Modes of CRISPR-Associated Transposon Homing.” Cell 184 (9): 2441–2453.e18. 10.1016/j.cell.2021.03.006.

43. Schargel, Richard D., Laura Chacon Machado, Shravanika Kumaran, Jordan E. Thesier, Alba Guarné, and Joseph E. Peters. 2025. “De Novo Engineered Guide RNA-Directed Transposition with TnpB-Family Proteins Reveals Features of Naturally Evolved Systems.” Preprint, bioRxiv, August 27. 10.1101/2025.08.27.672191.

44. Schmitz, Michael, Irma Querques, Seraina Oberli, Christelle Chanez, and Martin Jinek. 2022. “Structural Basis for the Assembly of the Type V CRISPR-Associated Transposon Complex.” Cell 185 (26): 4999–5010.e17. 10.1016/j.cell.2022.11.009.

45. Strecker, Jonathan, Alim Ladha, Zachary Gardner, et al. 2019. “RNA-Guided DNA Insertion with CRISPR-Associated Transposases.” Science 365 (6448): 48–53. 10.1126/science.aax9181.

46. Sumida, Kiera H., Reyes Núñez-Franco, Indrek Kalvet, et al. 2024. “Improving Protein Expression, Stability, and Function with ProteinMPNN.” Journal of the American Chemical Society 146 (3): 2054–61. 10.1021/jacs.3c10941.

47. Suzuki, Keiichiro, Yuji Tsunekawa, Reyna Hernandez-Benitez, et al. 2016. “In Vivo Genome Editing via CRISPR/Cas9 Mediated Homology-Independent Targeted Integration.” Nature 540 (7631): 144–49. 10.1038/nature20565.

48. Team, Chai Discovery, Jacques Boitreaud, Jack Dent, et al. 2024. “Chai-1: Decoding the Molecular Interactions of Life.” Preprint, bioRxiv, October 11. 10.1101/2024.10.10.615955.

49. Tou, Connor J., Benno Orr, and Benjamin P. Kleinstiver. 2023. “Precise Cut-and-Paste DNA Insertion Using Engineered Type V-K CRISPR-Associated Transposases.” Nature Biotechnology 41 (7): 968–79. 10.1038/s41587-022-01574-x.

50. Vakulskas, Christopher A., Daniel P. Dever, Garrett R. Rettig, et al. 2018. “A High-Fidelity Cas9 Mutant Delivered as a Ribonucleoprotein Complex Enables Efficient Gene Editing in Human Hematopoietic Stem and Progenitor Cells.” Nature Medicine 24 (8): 1216–24. 10.1038/s41591-018-0137-0.

51. Walker, Matt W. G., Sanne E. Klompe, Dennis J. Zhang, and Samuel H. Sternberg. 2023. “Novel Molecular Requirements for CRISPR RNA-Guided Transposition.” Nucleic Acids Research 51 (9): 4519–35. 10.1093/nar/gkad270.

52. Wang, Shukun, Clinton Gabel, Romana Siddique, Thomas Klose, and Leifu Chang. 2023. “Molecular Mechanism for Tn7-like Transposon Recruitment by a Type I-B CRISPR Effector.” Cell 186 (19): 4204–4215.e19. 10.1016/j.cell.2023.07.010.

53. Watson, Joseph L., David Juergens, Nathaniel R. Bennett, et al. 2023. “De Novo Design of Protein Structure and Function with RFdiffusion.” Nature 620 (7976): 1089–100. 10.1038/s41586-023-06415-8.

54. Witte, Isaac P., George D. Lampe, Simon Eitzinger, et al. 2025. “Programmable Gene Insertion in Human Cells with a Laboratory-Evolved CRISPR-Associated Transposase.” Science 388 (6748): eadt5199. 10.1126/science.adt5199.

55. Wu, Xuebing, David A. Scott, Andrea J. Kriz, et al. 2014. “Genome-Wide Binding of the CRISPR Endonuclease Cas9 in Mammalian Cells.” Nature Biotechnology 32 (7): 670–76. 10.1038/nbt.2889.

56. Yarnall, Matthew T. N., Eleonora I. Ioannidi, Cian Schmitt-Ulms, et al. 2023. “Drag-and-Drop Genome Insertion of Large Sequences without Double-Strand DNA Cleavage Using CRISPR-Directed Integrases.” Nature Biotechnology 41 (4): 500–512. 10.1038/s41587-022-01527-4.

57. Yin, Zhiqi, Mikalai Lapkouski, Wei Yang, and Robert Craigie. 2012. “Assembly of Prototype Foamy Virus Strand Transfer Complexes on Product DNA Bypassing Catalysis of Integration.” Protein Science: A Publication of the Protein Society 21 (12): 1849–57. 10.1002/pro.2166.

58. Yin, Zhiqi, Ke Shi, Surajit Banerjee, et al. 2016. “Crystal Structure of the Rous Sarcoma Virus Intasome.” Nature 530 (7590): 362–66. 10.1038/nature16950.

59. Zhang, Xiaozhu, Briana Van Treeck, Connor A. Horton, et al. 2025. “Harnessing Eukaryotic Retroelement Proteins for Transgene Insertion into Human Safe-Harbor Loci.” Nature Biotechnology 43 (1): 42–51. 10.1038/s41587-024-02137-y.

## References

60. “Accurate Structure Prediction of Biomolecular Interactions with AlphaFold 3 | Nature.” n.d. Accessed March 11, 2026. https://www.nature.com/articles/s41586-024-07487-w.

61. Ahn, Eungjin, Byungchul Kim, Soyoung Park, et al. 2023. “Batch Production of High-Quality Graphene Grids for Cryo-EM: Cryo-EM Structure of Methylococcus Capsulatus Soluble Methane Monooxygenase Hydroxylase.” ACS Nano 17 (6): 6011–22. 10.1021/acsnano.3c00463.

62. Alford, Rebecca F., Andrew Leaver-Fay, Jeliazko R. Jeliazkov, et al. 2017. “The Rosetta All-Atom Energy Function for Macromolecular Modeling and Design.” Journal of Chemical Theory and Computation 13 (6): 3031–48. 10.1021/acs.jctc.7b00125.

63. Altschul, S. F., T. L. Madden, A. A. Schäeer, et al. 1997. “Gapped BLAST and PSI-BLAST: A New Generation of Protein Database Search Programs.” Nucleic Acids Research 25 (17): 3389–402. 10.1093/nar/25.17.3389.

64. Ashkenazy, Haim, Shiran Abadi, Eric Martz, et al. 2016. “ConSurf 2016: An Improved Methodology to Estimate and Visualize Evolutionary Conservation in Macromolecules.” Nucleic Acids Research 44 (W1): W344–50. 10.1093/nar/gkw408.

65. Chaudhury, Sidhartha, Sergey Lyskov, and Jeerey J. Gray. 2010. “PyRosetta: A Script-Based Interface for Implementing Molecular Modeling Algorithms Using Rosetta.” Bioinformatics 26 (5): 689–91. 10.1093/bioinformatics/btq007.

66. Dauparas, Justas, Gyu Rie Lee, Robert Pecoraro, et al. 2025. “Atomic Context-Conditioned Protein Sequence Design Using LigandMPNN.” Nature Methods 22 (4): 717–23. 10.1038/s41592-025-02626-1.

67. Dieckhaus, Henry, Michael Brocidiacono, Nicholas Z. Randolph, and Brian Kuhlman. 2024. “Transfer Learning to Leverage Larger Datasets for Improved Prediction of Protein Stability Changes.” Proceedings of the National Academy of Sciences 121 (6): e2314853121. 10.1073/pnas.2314853121.

68. Emsley, P., B. Lohkamp, W. G. Scott, and K. Cowtan. 2010. “Features and Development of Coot.” Acta Crystallographica Section D: Biological Crystallography 66 (Pt 4): 486–501. 10.1107/S0907444910007493.

69. Finn, Robert D., Jody Clements, and Sean R. Eddy. 2011. “HMMER Web Server: Interactive Sequence Similarity Searching.” Nucleic Acids Research 39 (suppl_2): W29–37. 10.1093/nar/gkr367.

70. Fleishman, Sarel J., Andrew Leaver-Fay, Jacob E. Corn, et al. 2011. “RosettaScripts: A Scripting Language Interface to the Rosetta Macromolecular Modeling Suite.” PLOS ONE 6 (6): e20161. 10.1371/journal.pone.0020161.

71. Freschlin, Chase R., Kevin K. Yang, and Philip A. Romero. 2025. “Scalable and Cost-Eeicient Custom Gene Library Assembly from Oligopools.” Preprint, bioRxiv, March 22. 10.1101/2025.03.22.644747.

72. George, Jerrin Thomas, Christopher Acree, Jung-Un Park, et al. 2023. “Mechanism of Target Site Selection by Type V-K CRISPR-Associated Transposases.” Science (New York, N.Y.) 382 (6672): eadj8543. 10.1126/science.adj8543.

73. Glögl, Matthias, Aditya Krishnakumar, Robert J. Ragotte, et al. 2024. “Target-Conditioned Dieusion Generates Potent TNFR Superfamily Antagonists and Agonists.” Science 386 (6726): 1154–61. 10.1126/science.adp1779.

74. Guindon, Stéphane, Jean-François Dufayard, Vincent Lefort, Maria Anisimova, Wim Hordijk, and Olivier Gascuel. 2010. “New Algorithms and Methods to Estimate Maximum-Likelihood Phylogenies: Assessing the Performance of PhyML 3.0.” Systematic Biology 59 (3): 307–21. 10.1093/sysbio/syq010.

75. Katoh, Kazutaka, and Daron M. Standley. 2013. “MAFFT Multiple Sequence Alignment Software Version 7: Improvements in Performance and Usability.” Molecular Biology and Evolution 30 (4): 772–80. 10.1093/molbev/mst010.

76. Langmead, Ben, and Steven L. Salzberg. 2012. “Fast Gapped-Read Alignment with Bowtie 2.” Nature Methods 9 (4): 357–59. 10.1038/nmeth.1923.

77. Larralde, Martin. 2022. “Pyrodigal: Python Bindings and Interface to Prodigal, an Eeicient Method for Gene Prediction in Prokaryotes.” Journal of Open Source Software 7 (72): 4296. 10.21105/joss.04296.

78. Leaver-Fay, Andrew, Michael Tyka, Steven M. Lewis, et al. 2011. “ROSETTA3: An Object-Oriented Software Suite for the Simulation and Design of Macromolecules.” Methods in Enzymology 487: 545–74. 10.1016/B978-0-12-381270-4.00019-6.

79. Letunic, Ivica, and Peer Bork. 2021. “Interactive Tree Of Life (iTOL) v5: An Online Tool for Phylogenetic Tree Display and Annotation.” Nucleic Acids Research 49 (W1): W293–96. 10.1093/nar/gkab301.

80. Li, Weizhong, and Adam Godzik. 2006. “Cd-Hit: A Fast Program for Clustering and Comparing Large Sets of Protein or Nucleotide Sequences.” Bioinformatics (Oxford, England) 22 (13): 1658–59. 10.1093/bioinformatics/btl158.

81. Park, Seong Guk, Jung-Un Park, Esteban Dodero-Rojas, John A. Bryant Jr, Geetha Sankaranarayanan, and Elizabeth H. Kellogg. 2025. “Comprehensive Profiling of Activity and Specificity of RNA-Guided Transposons Reveals Opportunities to Engineer Improved Variants.” Nucleic Acids Research 53 (18): gkaf917. 10.1093/nar/gkaf917.

82. Pedregosa, Fabian, Gaël Varoquaux, Alexandre Gramfort, et al. 2011. “Scikit-Learn: Machine Learning in Python.” The Journal of Machine Learning Research 12 (null): 2825–30.

83. Rubin, Alan F., Hannah Gelman, Nathan Lucas, et al. 2017. “A Statistical Framework for Analyzing Deep Mutational Scanning Data.” Genome Biology 18 (1): 150. 10.1186/s13059-017-1272-5.

84. Russel, Jakob, Rafael Pinilla-Redondo, David Mayo-Muñoz, Shiraz A. Shah, and Søren J. Sørensen. 2020. “CRISPRCasTyper: Automated Identification, Annotation, and Classification of CRISPR-Cas Loci.” The CRISPR Journal 3 (6): 462–69. 10.1089/crispr.2020.0059.

85. Rybarski, James R., Kuang Hu, Alexis M. Hill, Claus O. Wilke, and Ilya J. Finkelstein. 2021. “Metagenomic Discovery of CRISPR-Associated Transposons.” Proceedings of the National Academy of Sciences 118 (49): e2112279118. 10.1073/pnas.2112279118.

86. Saito, Makoto, Alim Ladha, Jonathan Strecker, et al. 2021. “Dual Modes of CRISPR-Associated Transposon Homing.” Cell 184 (9): 2441–2453.e18. 10.1016/j.cell.2021.03.006.

87. Schrödinger, LLC. n.d. The PyMOL Molecular Graphics System. V. 3.0.

88. Steinegger, Martin, and Johannes Söding. 2017. “MMseqs2 Enables Sensitive Protein Sequence Searching for the Analysis of Massive Data Sets.” Nature Biotechnology 35 (11): 1026–28. 10.1038/nbt.3988.

89. Tareen, Ammar, and Justin B. Kinney. 2020. “Logomaker: Beautiful Sequence Logos in Python.” Bioinformatics 36 (7): 2272–74. 10.1093/bioinformatics/btz921.

90. Team, Chai Discovery, Jacques Boitreaud, Jack Dent, et al. 2024. “Chai-1: Decoding the Molecular Interactions of Life.” Preprint, bioRxiv, October 11. 10.1101/2024.10.10.615955.

91. Wang, Ray Yu-Ruei, Yifan Song, Benjamin A. Barad, Yifan Cheng, James S. Fraser, and Frank DiMaio. 2016. “Automated Structure Refinement of Macromolecular Assemblies from Cryo-EM Maps Using Rosetta.” eLife 5 (September): e17219. 10.7554/eLife.17219.

92. Watson, Joseph L., David Juergens, Nathaniel R. Bennett, et al. 2023. “De Novo Design of Protein Structure and Function with RFdieusion.” Nature 620 (7976): 1089–100. 10.1038/s41586-023-06415-8.

93. Zhang, Yang, and Jeerey Skolnick. 2005. “TM-Align: A Protein Structure Alignment Algorithm Based on the TM-Score.” Nucleic Acids Research 33 (7): 2302–9. 10.1093/nar/gki524.

